# Near-Infrared Turn-On Fluorogenic Probe for Versatile Detection of Inorganic Polyphosphates

**DOI:** 10.64898/2026.06.19.733421

**Authors:** Kenji Torii, Rūta Gerasimaitė, Gražvydas Lukinavičius

## Abstract

Inorganic polyphosphate (polyP) is a ubiquitous phosphate biopolymer involved in diverse cellular processes. Despite its significance, selective detection of polyP remains challenging because of its simple and highly charged structure. Here, we report a near-infrared (NIR) fluorogenic turn-on chemosensor for selective polyP detection and imaging, SiX-DPA-Zn. The probe combines a silicon-xanthene (SiX) fluorophore with a zinc(II)-coordinated 2,2′-dipicolylamine (DPA-Zn^2+^) recognition unit and shows more than 100-fold selectivity for inorganic polyP over ADP and ATP. SiX-DPA-Zn enables quantitative detection of polyP at micromolar concentrations in microplate assays and stains a broad range of polyP species, starting from tripolyphosphate, in polyacrylamide gels. In HEK293 cells expressing *Escherichia coli* polyphosphate kinase 1, the probe visualizes intracellular polyP and enables quantitative analysis of polyP levels in relation to nuclear proteins for example fibrillarin and nucleolin. Stimulated emission depletion (STED) microscopy further revealed subdiffraction-sized polyP granules within polyP aggregates. SiX-DPA-Zn is the first near-infrared (NIR) fluorogenic chemosensor for polyP that is compatible with multiple detection platforms, including microplate assays, polyacrylamide gel staining, confocal and super-resolution STED microscopy.

## Introduction

Inorganic polyphosphate (polyP), composed of chains of tens to thousands of orthophosphates, plays numerous roles in all forms of life, ranging from bacteria to mammalian^1–4^. For example, it functions as an energy and phosphate storage^5–7^, metal chelator^8,9^, buffer to alkaline^10^, protein protecting chaperone^11–13^, regulator of gene expression^14^, and protector against DNA damage^15^. PolyP is also associated with blood clotting^16^ and inflammation^17,18^. More recently, some studies have implicated polyP in neurodegeneration, including Parkinson’s and Alzheimer’s disease, and amyotrophic lateral sclerosis^19,20^. Given the broad and rapidly growing interest in polyP research, the development of chemical sensors capable of selectively detecting and visualizing polyP in biological systems has emerged as increasingly important direction in the field^21,20,4^.

Despite the high demand for polyP sensors, only a few probes that can sense polyP have been reported so far. The first example is 4,6-diamidino-2-phenylindole (DAPI), which is commonly used as a DNA stain (emission at around 450 nm). It is also used as a polyP sensor, detecting the red-shifted yellow emission (525–550 nm) that occurs upon binding to polyP^22,23^. However, DAPI exhibits similar yellow emission after binding to RNA (around 500 nm)^24,25^ and inositol pyrophosphates (around 550 nm)^26^ which reduces its specificity for polyP. JC-D dyes are recognized as the second generation of polyP sensors, aiming at higher specificity than DAPI^27^. Nevertheless, JC-D dyes exhibit strongly reduced fluorescence in physiological salt conditions due to weakened ionic dye–polyphosphate interactions, which severely limits reliable intracellular polyP staining^27^. Most recently, an excited-state intramolecular proton transfer (ESIPT)-based fluorescent probe for polyphosphate detection has been reported, enabling sensitive measurements in solution^28^. However, its applicability in biological systems remains unverified. Currently, the most widely used approach for visualizing polyphosphate (polyP) in mammalian cells employs polyphosphate-binding domain (PPBD), which binds polyP with high affinity and specificity^29–32^. It is typically fused with an epitope tag (e.g., Xpress tag) for immunostaining using primary and fluorophore-conjugated secondary antibodies. This conventional procedure is labor-intensive and time-consuming, and the multiple fixation and washing steps may also cause loss or redistribution of short-chain polyP. Hence, there is a strong need for the development of a chemosensor that enables facile detection of polyP in biological systems.

Several studies have shown that binuclear 2,2′-dipicolylamine (DPA)–metal (particularly Zn^2+^) complexes function as phosphate-binding motifs and have been widely used in chemosensors for various phosphate-containing substances^33–37^, nucleoside polyphosphates^38–45^ and pyrophosphates (PPi)^46–49,41^. We hypothesized that coupling a silicon-substituted xanthene fluorophore with a DPA– Zn^2+^ recognition motif would provide an effective platform for polyP sensing by tuning the electronic properties of the fluorophore and facilitate a turn-on response upon interaction with highly charged inorganic polyphosphates. Thus, we synthesized a novel inorganic polyP-selective near infra-red (NIR) emissive probe designated as SiX-DPA-Zn. Notably, SiX-DPA-Zn exhibits more than 100-fold higher selectivity for inorganic polyP, with micromolar affinity, while showing only weak binding to organic phosphates such as ATP and ADP (millimolar range). This selectivity enables quantitative determination of polyP in *Saccharomyces cerevisiae* lysates and monitoring enzymatic polyP degradation by alkaline phosphatase (ALP) and exopolyphosphatase (PPX) in microplate-based assays. In addition, SiX-DPA-Zn allows visualization of polyP in polyacrylamide gels, including short-chain species (<15 phosphate units) that are poorly detected by conventional dyes such as DAPI, JC-D7, and toluidine blue O (TB-O)^21^. Importantly, the probe is applicable to cellular imaging, enabling visualization of polyP in mammalian cells expressing *E. coli* polyphosphate kinase 1 (EcPPK) by confocal and STED microscopy.

## Results and Discussion

### Design and synthesis of SiX-DPA (-Zn)

In designing the sensor, we focused on a nucleoside polyphosphate turn-on probe incorporating two 2,2’-dipicolylamine (DPA)–Zn^2+^ complexes onto fluorescein (hereafter referred to as X-DPA-Zn), as reported by Hamachi et al^39,40^. In X-DPA-Zn, the oxygen atom bridged by the two DPA-Zn units forms a covalent bond with the C9 position of the xanthene scaffold, rendering the molecule non-fluorescent. Interaction with nucleoside di(tri)phosphates such as ADP (ATP), which contain two or more orthophosphate units, disrupts this C–O bond, leading to strong fluorescence^39^.

Encouraged by this intriguing property of the DPA-Zn complex, we hypothesized that inorganic polyP could also be recognized using a similar approach. To this end, we designed SiX-DPA-Zn, operating in the red fluorescence channel, which is less subject to interference by autofluorescence of cell components than blue and green channels employed by DAPI and other polyP probes. SiX-DPA-Zn consists of two DPA-Zn units and a silicon-substituted xanthene scaffold, with each DPA-Zn unit directly incorporated at the 1 and 8 positions of the fluorophore, as illustrated in **Figure 1A**. SiX-DPA-Zn was synthesized in 10 steps (9 pots) to afford the product in high purity (>99%) with an overall yield of 1.1%, shown in **Figure 1B**. The most critical step is the conjugation of DPA to the silicon-substituted xanthene. We found that a one-pot reaction, combining an Appel reaction followed by S_N_2 amination, afforded a good yield. This is likely attributable to the high instability of the dibrominated intermediate, which undergoes substantial decomposition during reaction workup and purification. Careful handling is also required during the purification of SiX-DPA, as it is easily oxidized in methanol. When purified using methanol, we observed a non-negligible byproduct in LC– MS with a mass increase of 16 and 34 Da, indicating the addition of one oxygen atom and an additional water molecule at the meso position (C9) of the xanthene core (**Figure S1**). Other studies have also reported that the Si-pyronin structure is unstable during purification^50^ or when stored in methanol^51^. This oxidation was successfully prevented by using acetonitrile during purification. After purification and lyophilization, the product purity was confirmed to be greater than 99% based on absorbance measurements at both 254 and 650 nm (**Figure S1**). The purified compound was stored in dimethyl sulfoxide (DMSO) at –80 °C for subsequent experiments. Detailed synthetic procedures and complete characterization data for all intermediates and final compound are provided in the Supporting Information.

**Figure 1.**
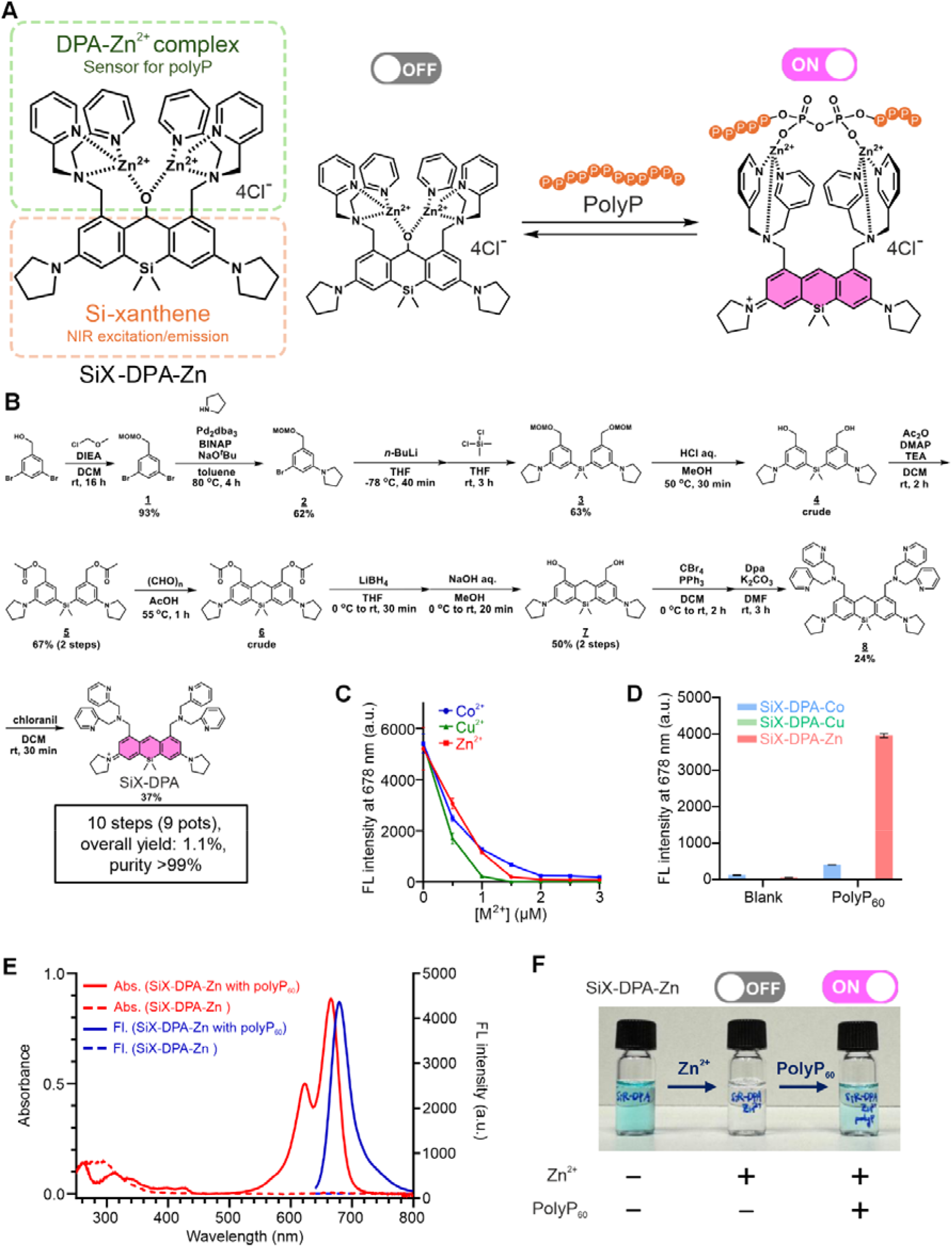
Molecular design, synthesis, and photophysical characterization of SiX-DPA. **(A)** Molecular design and proposed polyphosphate (polyP) sensing mechanism of SiX-DPA-Zn. **(B)** Synthetic route to SiX-DPA. **(C)** Fluorescence titration of SiX-DPA (1.0 µM) with ZnCl_2_, CuCl_2_ or CoCl_2_ in 20 mM HEPES buffer (pH 7.5). **(D)** Fluorescence response of SiX-DPA-Co, SiX-DPA-Cu, and SiX-DPA-Zn (1.0 µM) to addition of 100 µM polyP_60_. All data represented as mean ± s.d. of three independent replicates. **(E)** Absorption and fluorescence spectra of SiX-DPA-Zn (10 μM for absorbance measurement; 1.0 μM for fluorescence measurement) in the absence and presence of polyP_60_ (100 eq.) in 20 mM HEPES buffer (pH 7.5). **(F)** Color transition of 10 µM SiX-DPA solutions. From left to right: SiX-DPA alone, SiX-DPA with Zn^2+^, and SiX-DPA with Zn^2+^ plus polyP (1 mM).

### Photophysical properties of SiX-DPA and SiX-DPA-M

Purified SiX-DPA exhibits strong NIR absorption (*λ*_ex,max_ = 666 nm, *ε*= 1.1×10^5^ M^−1^cm^−1^) and fluorescence emission (*λ*_*em,max*_ = 678 nm, QY = 0.24, *τ* = 0.23 and 2.52 ns) in aqueous solution (**Figure S2** and **Table S1**). We evaluated effect of various divalent and trivalent metal ions and found that addition of two equivalents of Co^2+^, Cu^2+^, or Zn^2+^ resulted in substantial fluorescence quenching (**Figure S3, Figure 1C**). We propose that it is initiated by covalent bond formation between the C9 position of the xanthene scaffold and an oxygen atom bridging two DPA-Zn units (**Figure S4**). A similar mechanism has also been reported for X-DPA-Zn^39^. This quenched state (SiX-DPA-M) remains stable at pH>7.0 (**Figure S5**). However, both SiX-DPA-Zn and SiX-DPA-Co exhibit fluorescence recovery under acidic conditions starting from pH < 6.0, with complete recovery observed at pH 4.0. This is likely due to protonation of the DPA moiety, which decreases its affinity for zinc and cobalt ions^52,53^.We also evaluated the fluorescence stability of SiX-DPA across a range of pH values and observed a gradual decrease in fluorescence beginning at pH 8.5 (**Figure S5**). This can be attributed to nucleophilic attack of hydroxyl group (OH^-^) at the meso position, followed by subsequent oxidation. Therefore, we concluded that the fluorescence turn-off system operates effectively within the pH range of 7.0–8.0, which encompasses physiological conditions. Unless otherwise noted, all in vitro polyP detection experiments were performed in HEPES buffer at pH 7.5.

Next, we examined if SiX-DPA-M fluorescence intensity change can be used for polyP detection. SiX-DPA-Zn exhibited a markedly higher absorbance and fluorescence response to polyP_60_ than the other metal complexes (**Figure 1D,E)**. Its photophysical properties was similar to the metal-free SiX-DPA precursor, although it showed a threefold lower fluorescence quantum yield and a shorter fluorescence lifetime (**Table S1**). The color change of the SiX-DPA solution upon addition of Zn^2+^ and its subsequent recovery following supplementation with polyP_60_ could be readily observed as illustrated in **Figure 1F**. Therefore, SiX-PDA-Zn is a polyP sensor and we have investigated its properties further.

### Determination of SiX-DPA-Zn affinity to various phosphate-containing compounds

After identifying potential of SiX-DPA-Zn to detect polyP, we investigated its specificity by comparing its affinity to polyP and other phosphate-containing compounds. As analytes, we examined PolyP of different chain lengths (average lengths of 14 (polyP_14_), 60 (polyP_60_), and 130 (polyP_130_)), pyrophosphate (PPi) triphosphate (PPPi same as polyP_3_) and organic polyphosphates – adenosine diphosphate (ADP), adenosine triphosphate (ATP), and nucleic acids (calf thymus DNA and total yeast RNA). To examine possibility to use X-DPA-Zn as a polyP sensor in green channel, we also investigated it in parallel.

Titration experiments with SiX-DPA-Zn demonstrated significant differences in affinity between inorganic polyPs and other phosphate-containing substances. PPi showed the highest affinity of 2.4 µM while longer polyPs of all chain lengths (polyP_3_, polyP_14_, polyP_60_, and polyP_130_) exhibited slightly lower affinities with the chain length having little effect (*K*_*D*_ = 21, 16, 28, and 33 □ µM, respectively). ADP, ATP, and RNA showed considerably lower affinity (*K*_*D*_ > 1 □ mM). No fluorescence increase was observed upon addition of DNA (**Figure 2A** and **Table S2**). X-DPA-Zn showed similar affinity to all polyPs as SiX-DPA-Zn and negligible binding to nucleic acids. As previously reported^39^, it showed high affinity (*K*_*D*_ < 1 µM) to ATP and ADP (**Figure 2B**, and **Figure S6**). Moreover, SiX-DPA-Zn exhibited a larger fluorescence enhancement (*F*_*max*_/*F*_*0*_ = ~200-fold) upon binding to polyPs, compared to X-DPA-Zn (*F*_*max*_/*F*_*0*_ = ~10-fold) (**Table S2 and 3**). These results suggest that SiX-DPA-Zn can produce better contrast, is more specific to polyP and therefor is a better polyP sensor than X-DPA-Zn.

**Figure 2.**
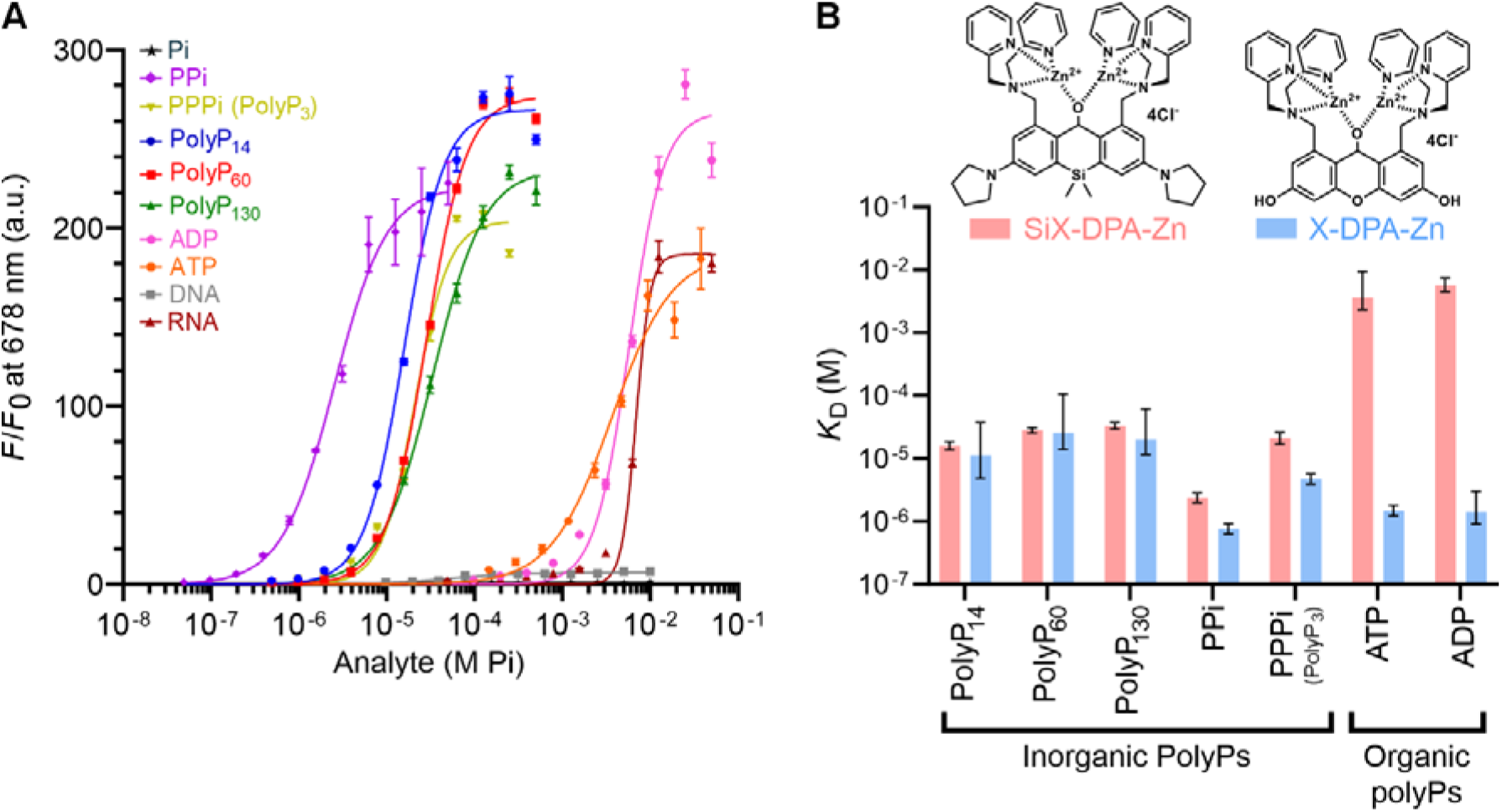
Binding affinity and selectivity of SiX-DPA-Zn toward phosphate-containing compounds. **(A)** Fluorescence titration curves of SiX-DPA-Zn (1.0 µM) with polyPs of different chain lengths (14, 60, and 130), PPi, PPPi (polyP_3_), ATP, ADP, DNA, and RNA. *λ*_*ex/em*_ = 620/678 nm. Fluorescence responses are shown as normalized intensities (F/F_0_), where F_0_ represents the fluorescence intensity in the absence of analyte. Concentrations of all phosphate-containing compounds are expressed as phosphate equivalents (M Pi) to allow straightforward comparison between different species. **(B)** Comparison of apparent dissociation constants (*K*_*D*_) of SiX-DPA-Zn and X-DPA-Zn toward polyPs, PPi, PPPi (polyP_3_), ATP, and ADP. *K*_*D*_ is calculated from Hills equation fitting. All data represented as mean ± s.d. of three independent replicates.

We hypothesize that the increased selectivity of SiX-DPA-Zn compared to X-DPA-Zn for polyP arises from altered electronic properties and improved scaffold preorganization, which together reduce binding to ATP and ADP. Replacement of the bridging oxygen at the 10-position with silicon changes the electronic structure of the fluorophore due to the lower electronegativity and higher polarizability of silicon. This modification alters charge distribution and may influence Zn(II)–DPA coordination and phosphate recognition. In addition, substitution with silicon leads to a small (~0.3 Å) increase in the distance between the 1- and 8-substituent positions, consistent with the larger atomic radius of silicon and longer Si–C bonds (**Figure S7**). This subtle geometric change may affect the preorganization of the binding sites.

After establishing ability of SiX-DPA-Zn to detect polyP, we examined its applicability in different methods used to characterize polyP. This includes measuring of polyP in solutions and in yeast extracts, visualizing polyP in PAA gels and staining polyP in fixed eukaryotic cells expressing *Ec*PPK1 for conventional and super-resolution fluorescence microscopy and cell sorting. Where applicable, we compared SiX-DPA-Zn performance to that of DAPI and JC-D7.

### Monitoring polyP degradation in vitro by polyphosphatases

To examine utility of fluorogenic SiX-DPA-Zn response for polyP quantification in vitro, we built calibration curves and found linear relationship between polyP concentration and fluorescence in 2-10 µM range (**Figure S8**). For SiX-DPA-Zn, the detection limit (0.5 µM polyP) was higher than that of DAPI^23^ and JC-D7^54^ (0.125 µM) (**Figure S8** and **S9**).

Several points should be considered when performing quantitative analysis of polyP using SiX-DPA-Zn. First, the apparent affinity of SiX-DPA-Zn for polyP decreases in the presence of high concentrations of phosphate, likely due to competition for probe binding (**Figure S10**). Second, ethylenediaminetetraacetic acid (EDTA), which is often used to quench metal-depending enzymatic reactions, can deprive Zn^2+^ of SiX-DPA-Zn with apparent affinity of 9 µM (**Figure S11**). This leads to fluorescence increase that might cause overestimation of polyP concentration. Third, the fluorescence intensity is influenced by magnesium ions, which are often present in biological buffers. This effect is presumably due to magnesium ions forming complexes with polyP^55,56^, thereby hindering its interaction with SiX–DPA–Zn. We also examined the effect of proteins using BSA as a standard protein and found that BSA did not impair the interaction between SiX-DPA-Zn and polyP **(Figure S10**). Therefore, we strongly recommend constructing calibration curves under identical buffer conditions, particularly with the same magnesium concentration and avoid using metal chelating agents (**Figure S12**). We also note that 0.1% Triton X-100 was added to HEPES buffer to maintain probe dispersion and ensure reliable fluorescence measurements in the following experiments (**Figure 3**).

**Figure 3.**
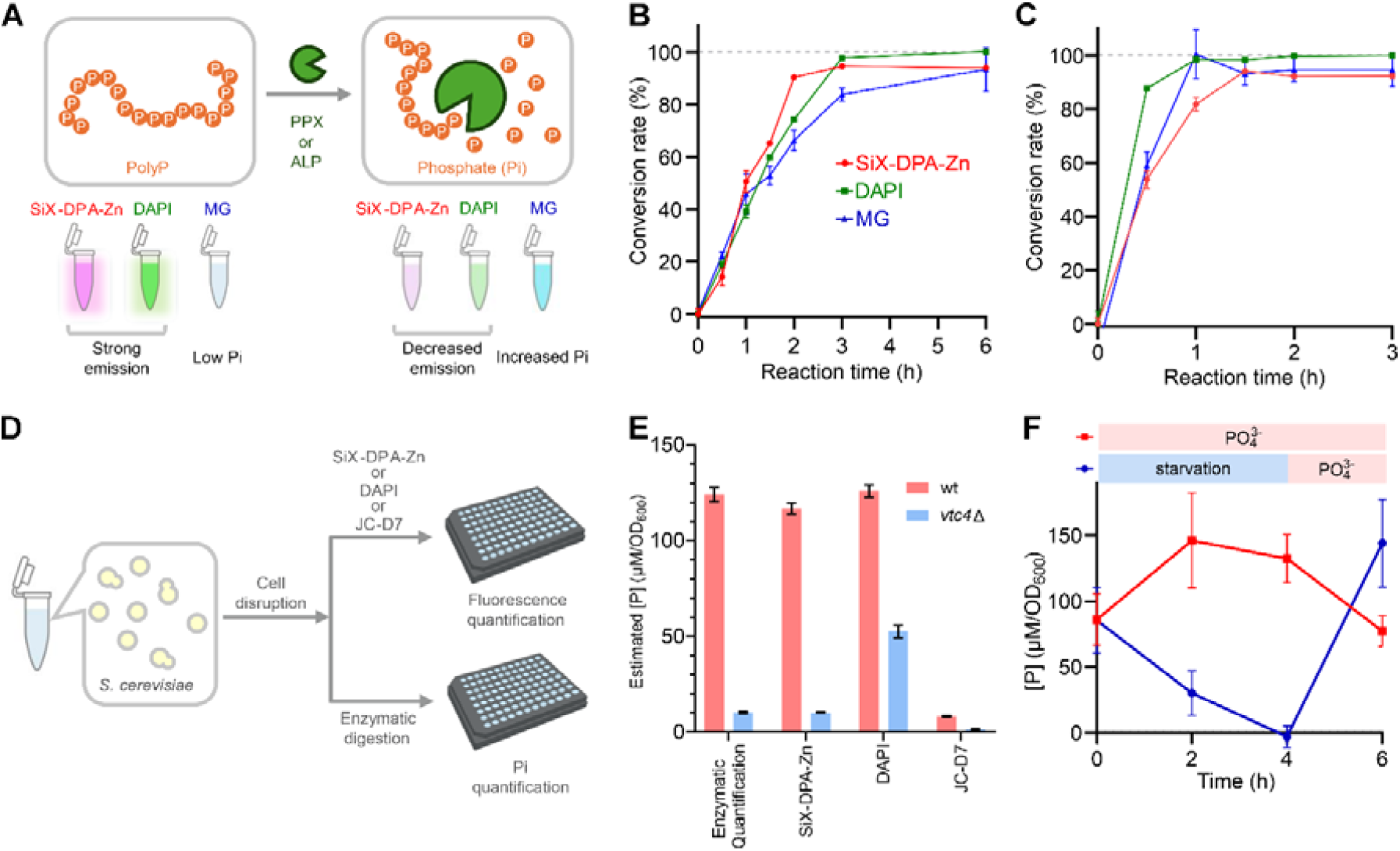
PolyP quantification using SiX-DPA-Zn. **(A)** Schematic illustration of enzymatic polyphosphate (polyP) degradation and its analysis using SiX-DPA-Zn fluorescence, DAPI fluorescence, and the malachite green (MG) phosphate assay. Time-dependent degradation of polyP_130_ (1 mM) monitored using end-point SiX-DPA-Zn assay during treatment with (**B**) exopolyphosphatase (PPX, 500 ng mL^− 1^) or (**C**) alkaline phosphatase (ALP, 50 µg mL^− 1^). Before fluorescence measurements, reaction mixtures were diluted 50-fold with 20 mM HEPES buffer. In case of SiX-DPA-Zn, the solution was additionally supplemented with 0.1% Triton X-100 in order to prevent probe aggregation. **(D)** Schematic illustration of polyP quantification in yeast (*S. cerevisiae*) extracts. **(E)** Determination of polyP concentration in stationary-phase wild-type yeast and the vacuolar transporter chaperone 4 knockout mutant (*vtc4*Δ) extracts using SiX-DPA-Zn, DAPI, JC-D7, and enzymatic quantification. PolyP concentrations were normalized to OD_600_. **(F)** Monitoring of polyP concentration under phosphate starvation. Red and Blue plot lines indicate Pi-starved and control cultures, respectively. After 4 hours of starvation, 10 mM KH_2_PO_4_ was added to the phosphate-depleted culture. All data represented as mean ± s.d. of three independent biological replicates.

After establishing conditions for polyP quantification with SiX-DPA-Zn, we monitored polyP digestion by *E. coli* exopolyphosphatase (PPX) and alkaline phosphatase (ALP)^57^ in the end-point assay (**Figure 3A**). We prepared a polyP_130_ solution, incubated it with each enzyme in the appropriate reaction buffer, and quantified polyP levels at defined time points using calibration curves generated with SiX-DPA-Zn or DAPI (**Figure S13A, B, D**, and **E**). Possible interference from the phosphate generated in the reaction and magnesium required for enzyme activity was counteracted by diluting the reaction mixture before probe addition and producing calibration curves with the identical buffer composition as the samples (**Figure S12**). At the same time, we determined concentration of released inorganic phosphate with malachite green (MG) assay^58^ (**Figure S13C** and **F**). All three methods yielded similar reaction time-courses (**Figure 3B and 3C**), indicating that SiX-DPA-Zn can be used to track polyP concentrations like DAPI and MG assay.

### Quantitative analysis of polyP in *Saccharomyces cerevisiae* cell extracts

To evaluate the applicability of SiX-DPA-Zn to analyze polyP from natural sources, we measured the concentration of polyP in yeast extracts (**Figure 3D**). As a control, we used *vtc4Δ* knockout mutant lacking polyphosphate synthase Vtc4, which contains negligible levels of polyP^59^. SiX-DPA-Zn was compared with other polyP sensitive dyes (DAPI and JC-D7) and with the enzymatic gold-standard method for polyP quantification, in which polyP is hydrolized by polyphosphatase and pyrophosphatase and the concentration of the relesed Pi easured by colorimetric reaction ^60,61^. After cell disruption with glass beads, the supernatant (cell lysate) was diluted to the appropriate concentration range for each assay (final concentration of 2–20 µM for the dye-based methods and 20–200 µM for the enzymatic method). The polyP concentration was then determined using calibration curves generated for each method. Enzymatic assay and SiX-DPA-Zn found similar polyP levels in wt yeast and low residual levels in *vtc4****Δ*** mutant (**Figure 3E**). Consistent to high sensitivity to salt, JC-D7 heavily underestimated the polyP concentration^27^. PolyP levels measured with DAPI in wt strain were similar, however this method reported unexpectedly high polyP in *vtc4****Δ***. RNase treatment of the lysates reduced fluorescent signal of DAPI in both samples and lead to undestimation of polyP levels in wt yeast (**Figure S14A**). The latter might be explaind by less efficient DAPI interaction with short polyPs^62^. Importantly, RNase treatment had no effect on SiX-DPA-Zn signal, suggesting negligible interference from RNA (**Figure S14B**).

Next, we used SiX-DPA-Zn to monitor polyP degradation and resynthesis during phosphate-starvation-replenishment cycle (**Figure 3F, Figure S15**). PolyP levels dropped to undetectable during 4 h of starvation and recovered to even higher levels within 2 hours upon phosphate addition to the medium.

Altogether these results demonstrate that SiX-DPA-Zn enables rapid and precise quantification of polyP concentration in *S. cerevisiae* without the enzymes and with no interference from RNA.

### Staining polyP in polyacrylamide gels

Assessment of polyP size distribution by polyacrylamide gel electrophoresis (PAGE) is an essential technique in investigating polyP metabolism, biological functions and the activity of polyP-synthesizing and -degrading enzymes. We compared SiX-DPA-Zn performance in staining polyP in gels to performance of other commonly used dyes – DAPI (positive and negative detection), JC-D7 and toluidine blue (TB-O)^63^. To this end, we ran four identical 30% gels with 10 nmol/well of different polyP species and stained them with the four dyes. To stain the gel with SiX-DPA-Zn, we incubated the gel with 10 µM probe in HEPES buffer supplemented with 20% ethanol and 2% glycerol for 30 min. We used the published protocols for other dyes (see Methods section for details). Upon excitation with 630 nm light, SiX-DPA-Zn-stained gel revealed fluorescent polyP smear (**Figure 4A**) with chain length distribution similar, but not identical to those produced by other methods (**Figure 4B**). The most striking difference was ability of SiX-DPA-Zn to stain polyPs as short as polyP_3_ that are not detectable with DAPI, JC-D7, or TB-O. This result is consistent with previous reports showing that DAPI and TB-O fail to detect polyP with fewer than 15 phosphate units^21^. To the best of our knowledge, JC-D7 has not been used for staining polyPs in gels before. In our hands, JC-D7 revealed similar polyP size distribution as DAPI and TB-O and also failed to detect short-chain polyPs.

**Figure 4.**
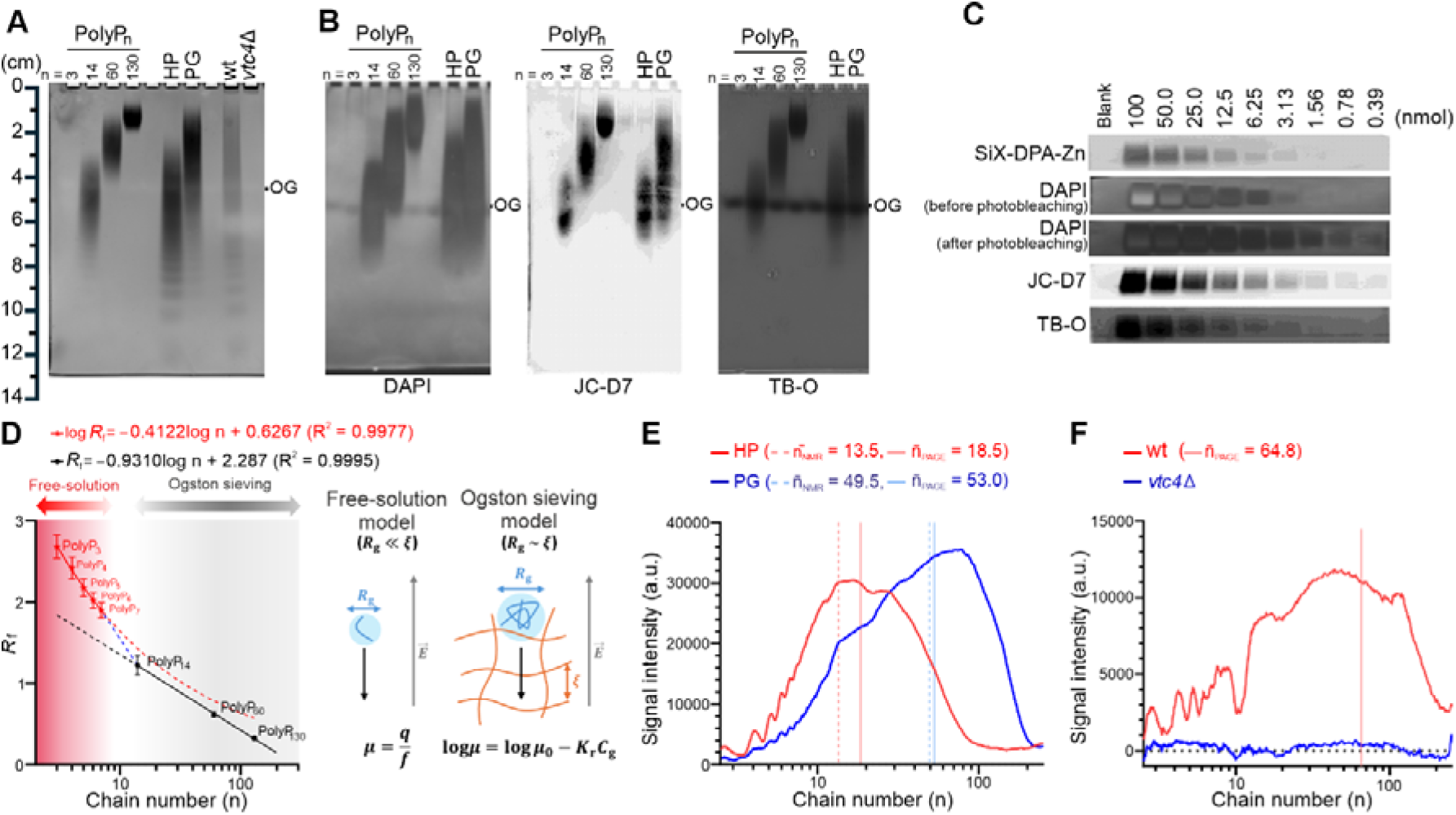
Polyphosphate (polyP) analysis by polyacrylamide gel electrophoresis (PAGE). **(A)** Detection of separated polyP species by SiX-DPA-Zn staining in 30% PAGE gels. Synthetic polyPs (polyP_3_, polyP_14_, polyP_60_, and polyP_130_), hexametaphosphate (HP), phosphate glass (PG), and RNA/polyP extracts from *S. cerevisiae* wild-type (wt) and knockout (*vtc4Δ*) cells were respectively separated by gel electrophoresis. OG is Orange G dye. **(B)** Comparison of staining performance for resolved polyP species in 30% PAGE gels using DAPI (negative staining), JC-D7, and toluidine blue (TB-O). OG represents the migration front of Orange G. **(C)** Various amounts of polyP60 (0.39–100 nmol) separated in 15% PAGE were stained with SiX-DPA-Zn, DAPI (before/after photobleaching), JC-D7, and TB-O. The images were compressed vertically for visualization. Full gels shown in Figure S17. **(D)** The relationship between relative mobility (*R*_*f*_) and chain number of polyP (n) determined using polyP_3_ and the resolved polyphosphate bands (polyP_4–7_) of HP (red line). Linear regression of the logarithm of chain number (n) versus relative mobility (*R*_*f*_) was performed using polyP_14_, polyP_60_, and polyP_130_ (black line). The blue dashed line connects the *R*_*f*_ values of polyP_7_ and polyP_14_ plotted against log n. The *R*_*f*_ values are expressed as the mean ± standard deviation from three independent experiments. Right panel shows schematic comparison of free-solution transport to long-chain Ogston sieving regimes. Free-solution mobility is governed by the effective charge (q) and friction coefficient (**f)**, related to the polymer radius of gyration (*R*_*g*_). Sieving mobility is governed by the retardation coefficient (*K*_*r*_) and gel mesh size (ξ). **(E, F)** Estimated chain-length distributions of **(E)** HP and PG, and **(F)** polyP extracts from *S. cerevisiae* wild-type (wt) and *vtc4*Δ cells calibrated using polyP_3_, polyP_14_, polyP_60_, and polyP_130_ standards. Dashed pale red and blue lines indicate average chain lengths determined by ^31^P NMR and corresponding solid lines indicate values estimated from PAGE analysis.

SiX-DPA-Zn successfully stained polyP extracted from *S. cerevisiae*. Strong polyP signal was observed in samples from wild-type cells, whereas no signal was detected in the *vtc4*Δ strain (**Figure 4A**). A weak, slowly-migrating discrete bands present in both samples completely disappeared upon RNase treatment, indicating that, similarly to DAPI, SiX-DPA-Zn can detect co-isolated RNA (**Figure S16**).

The sensitivity of polyP detection by SiX-DPA-Zn staining was comparable to DAPI before photobleaching and TB-O (~1.6 nmol), but lower than DAPI after photobleaching and JC-D7 (<0.4 nmol) **(Figure 4C** and **S17**). Despite this, SiX-DPA-Zn offers a clear advantage of fast and simple staining procedure without prolonged washing and photobleaching. Notably, it enables direct visualization of polyPs with a degree of polymerization below 10 on PA gels, which has not been previously achieved, except for methods requiring acid hydrolysis prior to methyl green staining^64^.

Ability to detect polyP_3_ provides an unambiguous reference and enables direct determination of the exact chain lengths of very short polyPs. Thereby we resolved individual polyP_3–7_ bands in HP sample. (**Figure 4A**). The relative mobility (Rf) of each band was calculated as the migration distance relative to the Orange G (OG) dye front. Rf values were then plotted against polyP chain length based on the Ogston sieving model (**Figure 4D**)^65–67^. Long-chain polyPs followed the expected sieving behavior. In contrast, short-chain polyPs (polyP_3-7_) required a different fitting model because their migration is better described by free-solution behavior, where the gel mesh has little effect. Both calibration curves showed excellent linearity with R^2^ > 0.995 based on at least three independent gels (**Figure 4D** and **Tables S4** and **S5**).

Based on the regressions obtained in **Figure 4D**, the distributions of the degree of polymerization for the polyphosphate samples HP and PG were determined and compared to those obtained by ^31^P NMR (**Figure 4E**). We found average chain lengths 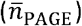 of 18.5 for HP and 53.0 for PG. The average chain lengths of HP 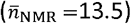 and PG 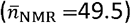 were determined by ^31^P NMR (**Figure S18**)^68,21^. The values from PAGE were slightly higher than those from NMR. This difference arises because fluorescence intensity reflects the total amount of bound probe and thus the mass of polyP rather than the number of chains. As a result, PAGE analysis yields a weight-average distribution, whereas NMR provides a number-average value, leading to an apparent shift toward longer chain lengths. The chain-length distribution of the polyP extract from *S. cerevisiae* was also analyzed in the same manner, yielding an average chain length of 64.8 (**Figure 4F**). The signal in the ^31^P NMR spectra was too weak to determine the average chain length for this sample. This demonstrates that our method is well suited for analyzing a broad range of polyP samples, including those not easily accessible by ^31^P NMR.

### PolyP visualization in mammalian cells

To assess the feasibility to use SiX-DPA-Zn for visualizing polyP by fluorescence microscopy, we stained a recently developed model T-Rex-293 cell line expressing *E*.*coli* polyphosphate kinase 1 (EcPPK1) under tetracycline repressor (TetR)^32^. Endogenous polyP levels in mammalian cells are low and thus challenging to detect. In this system, doxycycline induces expression of *E*.*coli* transgene, which leads to dramatically increased polyP levels. After 24 h of induction, cells were fixed, stained with SiX-DPA-Zn for 20 min, and imaged by fluorescence confocal microscope. Doxycycline-induced cells showed strong punctate fluorescence signals in the nucleus while little staining could be detected in noninduced controls (**Figure 5A**). A similar staining pattern was observed with red-shifted polyP-specific DAPI fluorescence (**Figure S19A**). We also tested JC-D7, a commercially available polyP-sensitive fluorophore^27^. However, no detectable fluorescence signal was observed in either induced or noninduced cells (**Figure S20**), consistent with the previous reports showing limited applicability of JC-D series dyes in mammalian cells^32^.

**Figure 5.**
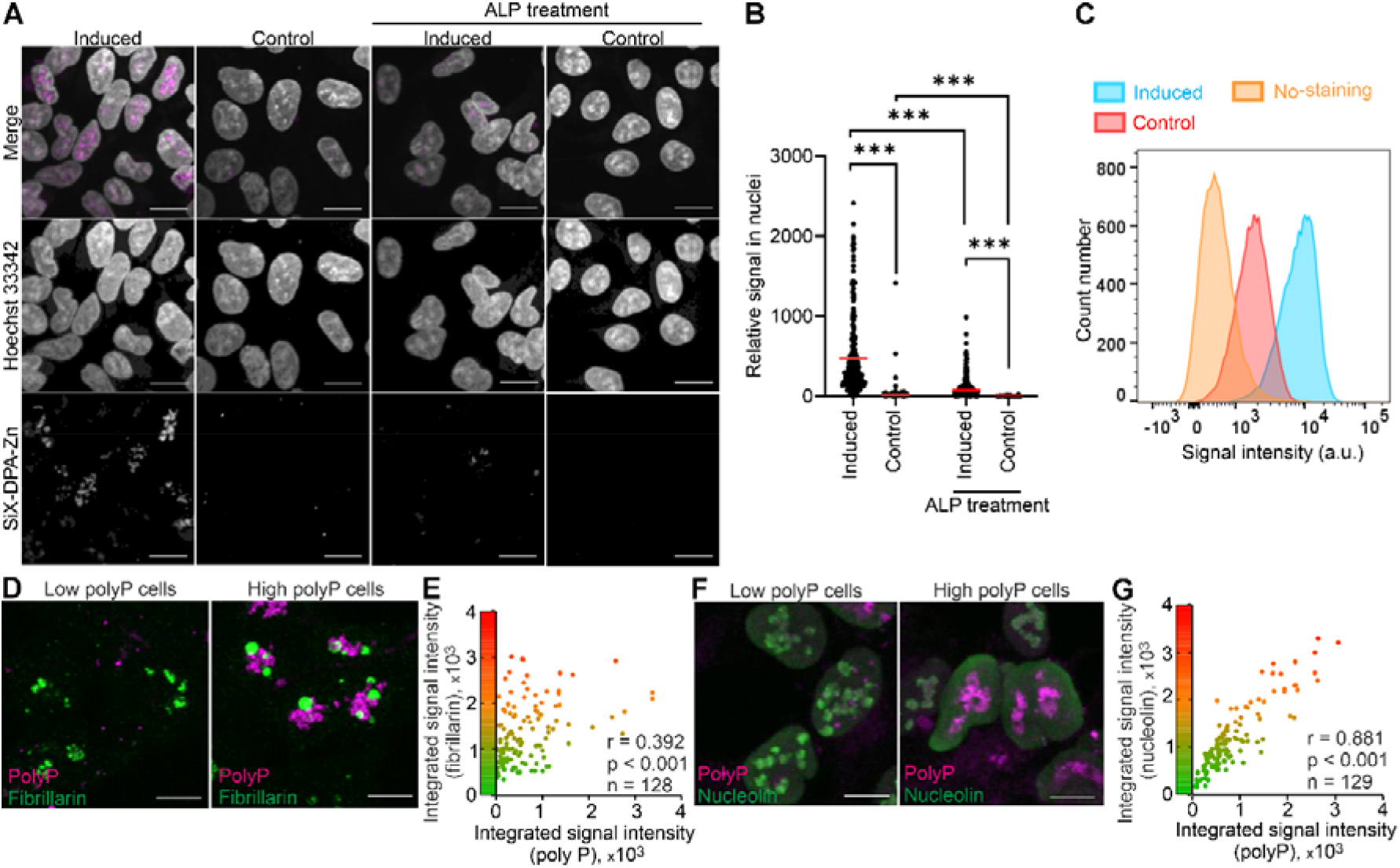
PolyP analysis in TREx-PPK cells using SiX-DPA-Zn. **(A)** Confocal images of TREx-PPK cells with and without doxycycline induction for 24 h. PolyP was stained with SiX-DPA-Zn (5.0 µM, magenta) and nuclei with Hoechst 33342 (1.0 µM, gray). PolyP depletion was achieved by alkaline phosphatase (ALP, 1.0 mg mL^− 1^) treatment for 1 h prior to analysis. Scale bar, 20 µm. **(B)** Quantification of nuclear polyP intensity from imaging data. The number of analyzed nuclei was 235 (induced, no ALP), 236 (control, no ALP), 205 (induced, with ALP), and 186 (control, with ALP). Red line represents the mean. The nucleus area was detected using Hoechst 33342 signal. *** - *p*<0.0001. **(C)** Flow cytometric analysis of single-cell polyP levels (excitation 633 nm, emission 670/14 nm). Cell numbers analyzed: 27,601 (induced), 26,841 (control), and 31,763 (unstained). **(D)** Co-staining of polyP (magenta) with fibrillarin (green) in induced cells showing low and high polyP levels. Scale bar, 10 µm. (**E**) Correlation analysis between polyP and fibrillarin signal intensities (n = 128 cells). **(F)** Co-staining of polyP (magenta) with nucleolin (green) in induced cells showing low and high polyP levels. Scale bar, 10 µm. (**G**) Correlation analysis between polyP and nucleolin signal intensities (n = 129 cells).

Pretreatment of samples with alkaline phosphatase (ALP) before staining significantly reduced the fluorescence signal, indicating that the nuclear signals detected by both, DAPI and SiX-DPA-Zn, reflect intracellular polyP (**Figure 5A** and **S19A**). Quantitative analysis of individual nuclei staining confirmed this conclusion for both dyes (**Figure 5B** and **S19B**). SiX-DPA-Zn signal was photostable and allowed three-dimensional (3D) imaging of polyP distribution throughout the nucleus volume. We often observed brightly stained 3D polyP aggregates of granular structure, reminiscent of polyP-rich domains previously detected by SEM^32^ (**Video S1**). These results demonstrate that SiX-DPA-Zn is a highly effective polyP-staining probe with low background fluorescence and a high signal-to-noise ratio, enabling accurate detection of polyP in cells.

Next, we applied the probe to fluorescence-activated cell sorting (FACS) analysis to examine large cell populations. Consistent with the microscopy results, doxycycline-induced cells showed a strong increase in SiX-DPA-Zn fluorescence compared with control cells (**Figure 5C**), demonstrating selective detection of polyP at the single-cell level. In contrast, FACS analysis using DAPI showed only minimal differences between induced and control cells, likely because of background fluorescence from DAPI bound to DNA and RNA (**Figure S21**). These results highlight the advantage of the fluorogenic turn-on response of SiX-DPA-Zn for flow cytometric analysis of cellular polyP.

From a spectral perspective, DAPI emits across both blue and green channels, while SiX-DPA-Zn is restricted to a single far-red channel. This spectral separation is particularly advantageous for studying intracellular polyP in complex cellular contexts using multicolor imaging. Previous study characterizing TREx-PPK1 cell line described disorganization of nucleolus and upregulation of nucleolar components, including fibrillarin and nucleolin, upon induction of EcPPK1 expression^32^. The latter conclusion was based mainly on protein levels measured by Western blotting, which reflects population-level analysis. Taking advantage of multicolor imaging and heterogeneous polyP load in cell population, we were able to confirm this observation at single-cell level and directly correlate polyP and nucleolin and fibrillarin (**Figure 5D–G** and **Figure S22** and **S23**). To this end, we co-stained cells with antibodies against respective proteins and SiX-DPA-Zn and quantified both components in individual cells. Cells with higher polyP levels showed increased fibrillarin staining signal (**Figure 5D** and **Figure S22**) and pearson’s correlation analysis revealed a moderate positive correlation (r = 0.39, p < 0.001, n = 128) (**Figure 5E**). An even stronger positive correlation was observed between polyP and nucleolin levels (r = 0.88, p < 0.001, n = 129) (**Figure 5G**).

### Subcellular distribution of polyP revealed by super-resolution microscopy

SiX-DPA-Zn is compatible with high resolution STED imaging because its near-infrared fluorescence can be efficiently depleted using a 775 nm STED laser, which provides significantly higher spatial resolution than confocal microscopy (**Figure 6A,B, Figure S24**). STED images revealed finer details of polyP aggregates with features smaller than the light diffraction limit and beyond resolution of conventional confocal microscopy (**Figure 6C, Figure S24**). The average size (FWHM) of polyP granules measured as 69 ± 41 nm by STED microscopy, compared with 367 ± 166 nm determined by confocal microscopy (**Figure 6C**).

**Figure 6.**
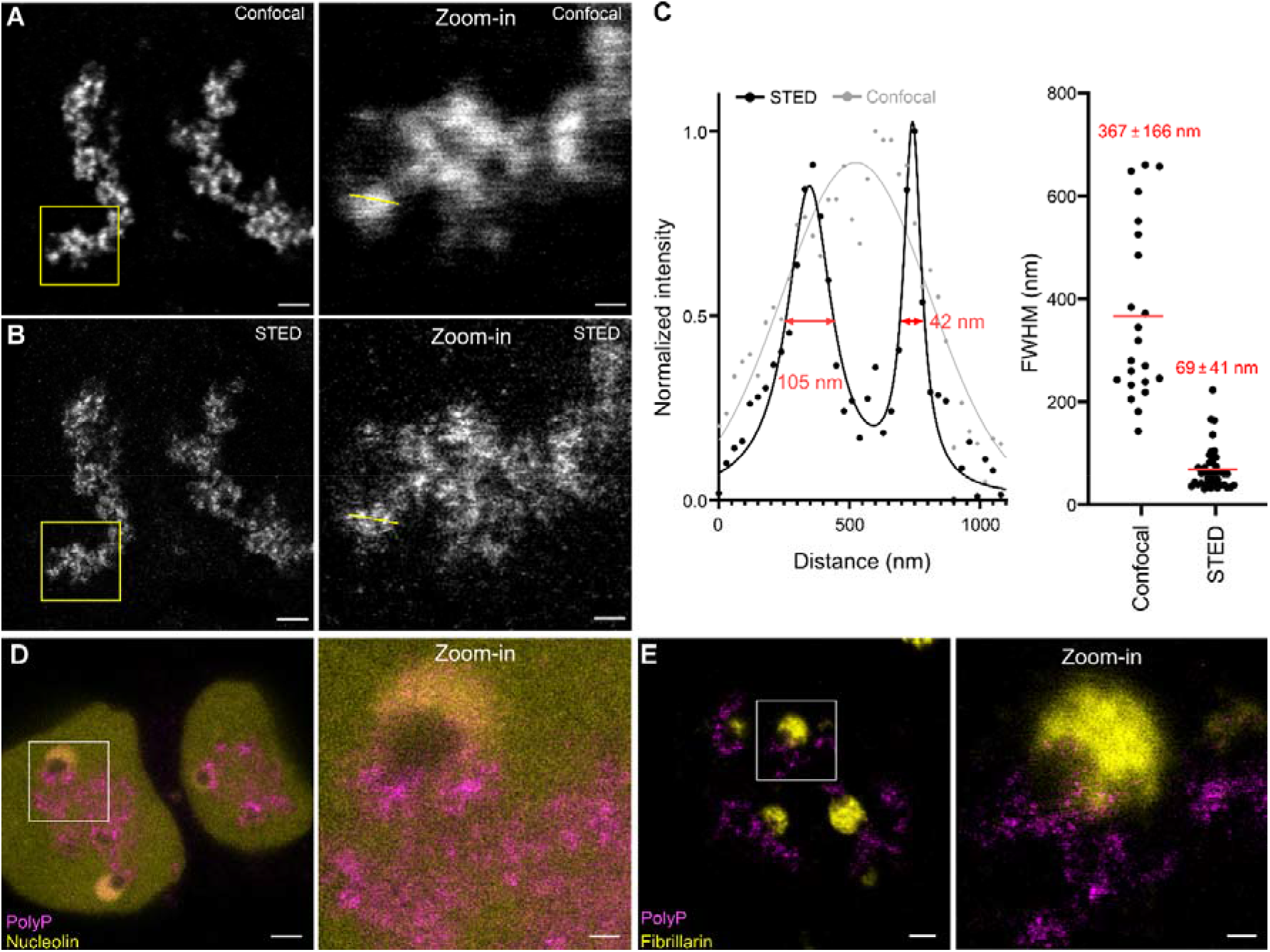
Subcellular distribution of polyP visualized using STED microscopy. (**A**) Representative confocal image of TREx-PPK cells induced with doxycycline for 24 h and stained with SiX-DPA-Zn (5.0 µM). Right panel: zoomed-in view of the region indicated by the white rectangle. (**B**) Corresponding STED images of the same nucleus. Right panel: zoomed-in view of the region indicated by the white rectangle. Scale bars, 2 µm and 500 nm (zoomed images). **(C)** Line profile plots of signal intensity along the yellow line in (**A**) and (**B**). Right graph: estimation of full width at half maximum (FWHM) of polyP granules from confocal (n = 22) and STED (n = 40) images from 20 nuclei. (**D, E**) Two-color STED imaging of polyP in TREx-PPK cells stained with SiX-DPA-Zn (5.0 µM, magenta), together with immunolabeled nucleolin (**D**) or fibrillarin (**E**) (yellow). Enlarged views on the right show the regions indicated by white squares. Scale bars, 2 µm and 500 nm (zoomed images).

Spatial distribution of the polyP relative to nucleolar proteins was highly heterogenous in the cell population (**Figure 6D, 6E, Figures S22** and **S23**), likely reflecting various degrees of nucleolar disorganization. Two distinct staining patterns were frequently observed. In cells with relatively weak SiX-DPA-Zn staining, polyP signals often colocalized with nucleolin. Both signals displayed a granular morphology characteristic of nucleolar substructures (**Figure S26**). In contrast, cells with strong polyP staining contained large and brightly labeled polyP aggregates with a similar granular appearance. In these cells, nucleolin was distributed diffusely throughout the nucleoplasm. In some cases, it accumulated in homogeneous spherical structures that did not overlap with polyP-positive regions and instead appeared spatially separated from them (**Figure 6D, Figure S27**). Fibrillarin was also detected in similar spherical compartments (**Figure 6E, Figure S25**). These observations are consistent with previous reports showing that elevated polyP levels induce collapse of the nucleolus into round homogeneous structures^32^. Notably, no internal organization could be resolved within these compartments, even by STED microscopy (**Figure 6D,E, Figures S25** and **S27**). The previous study also reported elevated phosphorus content in these rounded nucleolar structures, suggesting the presence of polyP^32^. However, we did not observe strong SiX-DPA-Zn staining within such compartments. A possible explanation is that SiX-DPA-Zn does not efficiently detect polyP when it is highly complexed with metal ions, as observed in our *in vitro* experiments (**Figure S10**). This suggests that SiX-DPA-Zn visualizes a specific, accessible fraction of intracellular polyP. Together, these results establish SiX-DPA-Zn as a useful probe for investigating the relationship between polyP accumulation and nucleolar organization.

## Conclusion

In this study, we developed SiX-DPA-Zn, the first near-infrared (NIR) fluorogenic chemosensor for selective detection of inorganic polyphosphate (polyP) *in vitro* and *in cellulo*. The probe exhibits high preference toward inorganic polyP over other abundant phosphate-containing molecules: ATP, ADP, DNA, and RNA. We demonstrated applicability of SiX-DPA-Zn for polyP detection in solutions, yeast cell extracts, polyacrylamide gels and in fixed cells, where for the first time we were able to image polyP by super-resolution fluorescence microscopy. In **Table S6**, we provide a detailed comparison of SiX-DPA-Zn and other dyes that are commonly used in polyP research.

In solution experiments, SiX-DPA-Zn signal is not affected by contaminating RNA, which is important when analyzing polyP from the natural sources. In polyacrylamide gels, SiX-DPA-Zn detects polyP chains as short as n=3, enabling analysis of short polyPs and unambiguous assignment of their chain length. In addition, it offers fast and simple staining procedure without extensive washing or prolonged and difficult to standardize UV irradiation. The most important, SiX-DPA-Zn can stain polyP in fixed cells thereby opening way for analysis of polyP levels and spacial distribution by fluorescent microscopy, including super-resolution STED imaging. Combined with immunostaining, it can become a valuable tool for analysis of processes involving polyP in cells. Moreover, SiX-DPA-Zn is compatible with fluorescence-activated cell sorting (FACS), enabling rapid and quantitative analysis of polyP levels in large cell populations.

Together, these features establish SiX-DPA-Zn as a broadly applicable tool for integrated quantitative and spatially resolved studies of polyP biology.

## Acknowledgments

The authors acknowledge the support of the Max Planck Society for this work. We are grateful to dr. Vladimir N. Belov for fruitful discussions and valuable suggestions. We are thankful for Jan Seikowski, Jens Schimpfhauser, and Jürgen Bienert from Facility for Synthetic Chemistry (MPI for Multidisciplinary Sciences) for the synthesis of X-DPA, as well as measuring NMR and mass-spectra of all synthesized componds. We thank the central analytics team at the Institute for Organic and Biomolecular Chemistry, Georg-August University of Göttingen, for acquiring the HRMS data of all synthesized compounds. We are thankful for Christine Quentin, Tanja Koenen, and Ellen Rothermel for their continuous care of the cell lines. We thank dr. Peter Lenart and dr. Antonio Politi from Facility for Light Microscopy (MPI for Multidisciplinary Sciences, Fassberg campus) for providing access to a spinning disk confocal microscope and FACS machine. We thank the RegeneTiss, Inc (Nagano, Japan) for kindly providing polyphosphate samples (polyP_14_, polyP_60_, and polyP_130_). We are grateful to dr. Adolfo Saiardi (Laboratory for Molecular Cell Biology, University College London) for a kind gift of PPK1-TREx cell line.

## References

(1) Kornberg, A. Inorganic Polyphosphate: Toward Making a Forgotten Polymer Unforgettable. J. Bacteriol. 1995, 177 (3), 491–496. 10.1128/jb.177.3.491-496.1995.

(2) Kornberg, A.; Rao, N. N.; Ault-Riché, D. Inorganic Polyphosphate: A Molecule of Many Functions. Annu. Rev. Biochem. 1999, 68 (Volume 68, 1999), 89–125. 10.1146/annurev.biochem.68.1.89.

(3) Brown, M. R. W.; Kornberg, A. Inorganic Polyphosphate in the Origin and Survival of Species. Proc. Natl. Acad. Sci. U. S. A. 2004, 101 (46), 16085–16087. 10.1073/pnas.0406909101.

(4) Rai, A.; Jakob, U. Polyphosphate: A Cellular Swiss Army Knife. Curr. Opin. Biotechnol. 2025, 93, 103303. 10.1016/j.copbio.2025.103303.

(5) Lynn, W. S.; Brown, R. H. Synthesis of Polyphosphate by Rat Liver Mitochondria. Biochem. Biophys. Res. Commun. 1963, 11 (5), 367–371. 10.1016/0006-291X(63)90124-4.

(6) Skorko, R. Polyphosphate as a Source of Phosphoryl Group in Protein Modification in the Archaebacterium Sulfolobus Acidocaldarius. Biochimie 1989, 71 (9), 1089–1093. 10.1016/0300-9084(89)90115-6.

(7) Baev, Y. A.; Angelova, R. P; Abramov, Y. A. Inorganic Polyphosphate Is Produced and Hydrolyzed in F0F1-ATP Synthase of Mammalian Mitochondria. Biochem. J. 2020, 477 (8), 1515–1524. 10.1042/BCJ20200042.

(8) Omelon, S.; Georgiou, J.; Henneman, Z. J.; Wise, L. M.; Sukhu, B.; Hunt, T.; Wynnyckyj, C.; Holmyard, D.; Bielecki, R.; Grynpas, M. D. Control of Vertebrate Skeletal Mineralization by Polyphosphates. PLOS ONE 2009, 4 (5), e5634. 10.1371/journal.pone.0005634.

(9) Andreeva, N.; Ryazanova, L.; Dmitriev, V.; Kulakovskaya, T.; Kulaev, I. Adaptation of Saccharomyces Cerevisiae to Toxic Manganese Concentration Triggers Changes in Inorganic Polyphosphates. FEMS Yeast Res. 2013, 13 (5), 463–470. 10.1111/1567-1364.12049.

(10) Pick, U.; Weiss, M. Polyphosphate Hydrolysis within Acidic Vacuoles in Response to Amine-Induced Alkaline Stress in the Halotolerant Alga Dunaliella Salina. Plant Physiol. 1991, 97 (3), 1234–1240. 10.1104/pp.97.3.1234.

(11) Gray, M. J.; Wholey, W.-Y.; Wagner, N. O.; Cremers, C. M.; Mueller-Schickert, A.; Hock, N. T.; Krieger, A. G.; Smith, E. M.; Bender, R. A.; Bardwell, J. C. A.; Jakob, U. Polyphosphate Is a Primordial Chaperone. Mol. Cell 2014, 53 (5), 689–699. 10.1016/j.molcel.2014.01.012.

(12) Yoo, N. G.; Dogra, S.; Meinen, B. A.; Tse, E.; Haefliger, J.; Southworth, D. R.; Gray, M. J.; Dahl, J.-U.; Jakob, U. Polyphosphate Stabilizes Protein Unfolding Intermediates as Soluble Amyloid-like Oligomers. J. Mol. Biol. 2018, 430 (21), 4195–4208. 10.1016/j.jmb.2018.08.016.

(13) Guan, J.; Jakob, U. The Protein Scaffolding Functions of Polyphosphate. J. Mol. Biol. 2024, 436 (14), 168504. 10.1016/j.jmb.2024.168504.

(14) Bondy-Chorney, E.; Abramchuk, I.; Nasser, R.; Holinier, C.; Denoncourt, A.; Baijal, K.; McCarthy, L.; Khacho, M.; Lavallée-Adam, M.; Downey, M. A Broad Response to Intracellular Long-Chain Polyphosphate in Human Cells. Cell Rep. 2020, 33 (4), 108318. 10.1016/j.celrep.2020.108318.

(15) Bru, S.; Samper-Martín, B.; Quandt, E.; Hernández-Ortega, S.; Martínez-Laínez, J. M.; Garí, E.; Rafel, M.; Torres-Torronteras, J.; Martí, R.; Ribeiro, M. P. C.; Jiménez, J.; Clotet, J. Polyphosphate Is a Key Factor for Cell Survival after DNA Damage in Eukaryotic Cells. DNA Repair 2017, 57, 171–178. 10.1016/j.dnarep.2017.08.001.

(16) Smith, S. A.; Mutch, N. J.; Baskar, D.; Rohloff, P.; Docampo, R.; Morrissey, J. H. Polyphosphate Modulates Blood Coagulation and Fibrinolysis. Proc. Natl. Acad. Sci. 2006, 103 (4), 903–908. 10.1073/pnas.0507195103.

(17) Müller, F.; Mutch, N. J.; Schenk, W. A.; Smith, S. A.; Esterl, L.; Spronk, H. M.; Schmidbauer, S.; Gahl, W. A.; Morrissey, J. H.; Renné, T. Platelet Polyphosphates Are Proinflammatory and Procoagulant Mediators In Vivo. Cell 2009, 139 (6), 1143–1156. 10.1016/j.cell.2009.11.001.

(18) Hassanian, S. M.; Dinarvand, P.; Smith, S. A.; Rezaie, A. R. Inorganic Polyphosphate Elicits Pro-inflammatory Responses through Activation of the Mammalian Target of Rapamycin Complexes 1 and 2 in Vascular Endothelial Cells. J. Thromb. Haemost. 2015, 13 (5), 860–871. 10.1111/jth.12899.

(19) Arredondo, C.; Cefaliello, C.; Dyrda, A.; Jury, N.; Martinez, P.; Díaz, I.; Amaro, A.; Tran, H.; Morales, D.; Pertusa, M.; Stoica, L.; Fritz, E.; Corvalán, D.; Abarzúa, S.; Méndez-Ruette, M.; Fernández, P.; Rojas, F.; Kumar, M. S.; Aguilar, R.; Almeida, S.; Weiss, A.; Bustos, F. J.; González-Nilo, F.; Otero, C.; Tevy, M. F.; Bosco, D. A.; Sáez, J. C.; Kähne, T.; Gao, F.-B.; Berry, J. D.; Nicholson, K.; Sena-Esteves, M.; Madrid, R.; Varela, D.; Montecino, M.; Brown, R. H.; van Zundert, B. Excessive Release of Inorganic Polyphosphate by ALS/FTD Astrocytes Causes Non-Cell-Autonomous Toxicity to Motoneurons. Neuron 2022, 110 (10), 1656–1670.e12. 10.1016/j.neuron.2022.02.010.

(20) Garcés, P.; Amaro, A.; Montecino, M.; van Zundert, B. Inorganic Polyphosphate: From Basic Research to Diagnostic and Therapeutic Opportunities in ALS/FTD. Biochem. Soc. Trans. 2024, 52 (1), 123–135. 10.1042/BST20230257.

(21) Christ, J. J.; Willbold, S.; Blank, L. M. Methods for the Analysis of Polyphosphate in the Life Sciences. Anal. Chem. 2020, 92 (6), 4167–4176. 10.1021/acs.analchem.9b05144.

(22) Gomes, F. M.; Ramos, I. B.; Wendt, C.; Girard-Dias, W.; De Souza, W.; Machado, E. A.; Miranda, K. New Insights into the in Situ Microscopic Visualization and Quantification of Inorganic Polyphosphate Stores by 4’,6-Diamidino-2-Phenylindole (DAPI)-Staining. Eur. J. Histochem. EJH 2013, 57 (4), e34. 10.4081/ejh.2013.e34.

(23) Aschar-Sobbi, R.; Abramov, A. Y.; Diao, C.; Kargacin, M. E.; Kargacin, G. J.; French, R. J.; Pavlov, E. High Sensitivity, Quantitative Measurements of Polyphosphate Using a New DAPI-Based Approach. J. Fluoresc. 2008, 18 (5), 859–866. 10.1007/s10895-008-0315-4.

(24) Omelon, S.; Georgiou, J.; Habraken, W. A Cautionary (Spectral) Tail: Red-Shifted Fluorescence by DAPI–DAPI Interactions. Biochem. Soc. Trans. 2016, 44 (1), 46–49. 10.1042/BST20150231.

(25) Kapuscinski, J. Interactions of Nucleic Acids with Fluorescent Dyes: Spectral Properties of Condensed Complexes. J. Histochem. Cytochem. 1990, 38 (9), 1323–1329. 10.1177/38.9.1696951.

(26) Kolozsvari, B.; Parisi, F.; Saiardi, A. Inositol Phosphates Induce DAPI Fluorescence Shift. Biochem. J. 2014, 460 (3), 377–385. 10.1042/BJ20140237.

(27) Angelova, P. R.; Agrawalla, B. K.; Elustondo, P. A.; Gordon, J.; Shiba, T.; Abramov, A. Y.; Chang, Y.-T.; Pavlov, E. V. In Situ Investigation of Mammalian Inorganic Polyphosphate Localization Using Novel Selective Fluorescent Probes JC-D7 and JC-D8. ACS Chem. Biol. 2014, 9 (9), 2101–2110. 10.1021/cb5000696.

(28) Roy, S. K.; Moser, S.; Dürr-Mayer, T.; Hinkelmann, R.; Jessen, H. J. ESIPT Fluorescence Turn-on Sensors for Detection of Short Chain Inorganic Polyphosphate in Water. Org. Biomol. Chem. 2025, 23 (6), 1373–1379. 10.1039/D4OB01926A.

(29) Saito, K.; Ohtomo, R.; Kuga-Uetake, Y.; Aono, T.; Saito, M. Direct Labeling of Polyphosphate at the Ultrastructural Level in Saccharomyces Cerevisiae by Using the Affinity of the Polyphosphate Binding Domain of Escherichia Coli Exopolyphosphatase. Appl. Environ. Microbiol. 2005, 71 (10), 5692–5701. 10.1128/AEM.71.10.5692-5701.2005.

(30) Samper-Martín, B.; Sarrias, A.; Lázaro, B.; Pérez-Montero, M.; Rodríguez-Rodríguez, R.; Ribeiro, M. P. C.; Bañón, A.; Wolfgeher, D.; Jessen, H. J.; Alsina, B.; Clotet, J.; Kron, S. J.; Saiardi, A.; Jiménez, J.; Bru, S. Polyphosphate Degradation by Nudt3-Zn2+ Mediates Oxidative Stress Response. Cell Rep. 2021, 37 (7), 110004. 10.1016/j.celrep.2021.110004.

(31) Jiménez, J.; Lázaro, B.; Sarrias, A.; Tadeo, F. J.; Pérez-Montero, M.; Clotet, J.; Bru, S. Protocol to Quantify Polyphosphate in Human Cell Lines Using a Tagged PPBD Peptide. STAR Protoc. 2022, 3 (2), 101363. 10.1016/j.xpro.2022.101363.

(32) Borghi, F.; Azevedo, C.; Johnson, E.; Burden, J. J.; Saiardi, A. A Mammalian Model Reveals Inorganic Polyphosphate Channeling into the Nucleolus and Induction of a Hyper-Condensate State. Cell Rep. Methods 2024, 4 (7), 100814. 10.1016/j.crmeth.2024.100814.

(33) Nag, A.; Das, S. Fluorescent Sensors of Phosphate Containing Biomolecules. Isr. J. Chem. 2021, 61 (3–4), 169–184. 10.1002/ijch.202000087.

(34) Wongkongkatep, J.; Ojida, A.; Hamachi, I. Fluorescence Sensing of Inorganic Phosphate and Pyrophosphate Using Small Molecular Sensors and Their Applications. In Phosphate Labeling and Sensing in Chemical Biology; Jessen, H. J., Ed.; Springer International Publishing: Cham, 2017; pp 1–33. 10.1007/978-3-319-60357-5_1.

(35) Bencini, A.; Bartoli, F.; Caltagirone, C.; Lippolis, V. Zinc(II)-Based Fluorescent Dyes: Luminescence Modulation by Phosphate Anion Binding. Dyes Pigments 2014, 110, 169–192. 10.1016/j.dyepig.2014.04.009.

(36) Sakamoto, T.; Ojida, A.; Hamachi, I. Molecular Recognition, Fluorescence Sensing, and Biological Assay of Phosphate Anion Derivatives Using Artificial Zn(Ii)–Dpa Complexes. Chem. Commun. 2009, 0 (2), 141–152. 10.1039/B812374H.

(37) Tien Ngo, H.; Liu, X.; A. Jolliffe K. Anion Recognition and Sensing with Zn(Ii)–Dipicolylamine Complexes. Chem. Soc. Rev. 2012, 41 (14), 4928–4965. 10.1039/C2CS35087D.

(38) Swamy, K. M. K.; Kwon, S. K.; Lee, H. N.; Shantha Kumar, S. M.; Kim, J. S.; Yoon, J. Fluorescent Sensing of Pyrophosphate and ATP in 100% Aqueous Solution Using a Fluorescein Derivative and Mn2+. Tetrahedron Lett. 2007, 48 (49), 8683–8686. 10.1016/j.tetlet.2007.10.022.

(39) Ojida, A.; Takashima, I.; Kohira, T.; Nonaka, H.; Hamachi, I. Turn-on Fluorescence Sensing of Nucleoside Polyphosphates Using a Xanthene-Based Zn(Ii) Complex Chemosensor. J. Am. Chem. Soc. 2008, 130 (36), 12095–12101. 10.1021/ja803262w.

(40) Kurishita, Y.; Kohira, T.; Ojida, A.; Hamachi, I. Organelle-Localizable Fluorescent Chemosensors for Site-Specific Multicolor Imaging of Nucleoside Polyphosphate Dynamics in Living Cells. J. Am. Chem. Soc. 2012, 134 (45), 18779–18789. 10.1021/ja308754g.

(41) Wang, J.; Liu, X.; Pang, Y. A Benzothiazole-Based Sensor for Pyrophosphate (PPi) and ATP: Mechanistic Insight for Anion-Induced ESIPT Turn-On. J. Mater. Chem. B 2014, 2 (38), 6634–6638. 10.1039/C4TB01109K.

(42) Jin, X.; Gao, J.; Xie, P.; Yu, M.; Wang, T.; Zhou, H.; Ma, A.; Wang, Q.; Leng, X.; Zhang, X. Dual-Functional Probe Based on Rhodamine for Sequential Cu2+ and ATP Detection in Vivo. Spectrochim. Acta. A. Mol. Biomol. Spectrosc. 2018, 204, 657–664. 10.1016/j.saa.2018.06.094.

(43) Gu, Q.-S.; Li, T.; Liu, T.; Yu, G.; Mao, G.-J.; Xu, F.; Li, C.-Y. Recent Advances in Design Strategies and Imaging Applications of Fluorescent Probes for ATP. Chemosensors 2023, 11 (7), 417. 10.3390/chemosensors11070417.

(44) Gomes, L. J.; Carrilho, J. P.; Pereira, P. M.; Moro, A. J. A Near InfraRed Emissive Chemosensor for Zn2+ and Phosphate Derivatives Based on a Di-(2-Picolyl)Amine-Styrylflavylium Push-Pull Fluorophore. Sensors 2023, 23 (1), 471. 10.3390/s23010471.

(45) Singh, H.; Sreedharan, S.; Tiwari, R.; Walther, C.; Smythe, C.; Pramanik, S. K.; Thomas, J. A.; Das, A. A Fluorescent Chemodosimeter for Organelle-Specific Imaging of Nucleoside Polyphosphate Dynamics in Living Cells. Cryst. Growth Des. 2018, 18 (11), 7199–7206. 10.1021/acs.cgd.8b01409.

(46) Jang, Y. J.; Jun, E. J.; Lee, Y. J.; Kim, Y. S.; Kim, J. S.; Yoon, J. Highly Effective Fluorescent and Colorimetric Sensors for Pyrophosphate over H2PO4-in 100% Aqueous Solution. J. Org. Chem. 2005, 70 (23), 9603–9606. 10.1021/jo0509657.

(47) Kim, M. J.; Swamy, K. M. K.; Lee, K. M.; Jagdale, A. R.; Kim, Y.; Kim, S.-J.; Yoo, K. H.; Yoon, J. Pyrophosphate Selective Fluorescent Chemosensors Based on Coumarin–DPA–Cu(II) Complexes. Chem. Commun. 2009, No. 46, 7215–7217. 10.1039/B913809A.

(48) Chen, W.-H.; Xing, Y.; Pang, Y. A Highly Selective Pyrophosphate Sensor Based on ESIPT Turn-On in Water. Org. Lett. 2011, 13 (6), 1362–1365. 10.1021/ol200054w.

(49) Zhang, J. F.; Kim, S.; Han, J. H.; Lee, S.-J.; Pradhan, T.; Cao, Q. Y.; Lee, S. J.; Kang, C.; Kim, J. S. Pyrophosphate-Selective Fluorescent Chemosensor Based on 1,8-Naphthalimide–DPA–Zn(II) Complex and Its Application for Cell Imaging. Org. Lett. 2011, 13 (19), 5294–5297. 10.1021/ol202159x.

(50) Romieu, A.; Dejouy, G.; Valverde, I. E. Quest for Novel Fluorogenic Xanthene Dyes: Synthesis, Spectral Properties and Stability of 3-Imino-3H-Xanthen-6-Amine (Pyronin) and Its Silicon Analog. Tetrahedron Lett. 2018, 59 (52), 4574–4581. 10.1016/j.tetlet.2018.11.031.

(51) Pastierik, T.; Šebej, P.; Medalová, J.; Štacko, P.; Klán, P. Near-Infrared Fluorescent 9-Phenylethynylpyronin Analogues for Bioimaging. J. Org. Chem. 2014, 79 (8), 3374–3382. 10.1021/jo500140y.

(52) Zastrow, M. L.; Radford, R. J.; Chyan, W.; Anderson, C. T.; Zhang, D. Y.; Loas, A.; Tzounopoulos, T.; Lippard, S. J. Reaction-Based Probes for Imaging Mobile Zinc in Live Cells and Tissues. ACS Sens. 2016, 1 (1), 32–39. 10.1021/acssensors.5b00022.

(53) Chyan, W.; Zhang, D. Y.; Lippard, S. J.; Radford, R. J. Reaction-Based Fluorescent Sensor for Investigating Mobile Zn2+ in Mitochondria of Healthy versus Cancerous Prostate Cells. Proc. Natl. Acad. Sci. U. S. A. 2014, 111 (1), 143–148. 10.1073/pnas.1310583110.

(54) Yang, X.; Gao, R.; Zhang, Q.; Yung, C. C. M.; Yin, H.; Li, J. Quantification of Polyphosphate in Environmental Planktonic Samples Using a Novel Fluorescence Dye JC-D7. Environ. Sci. Technol. 2024. 10.1021/acs.est.4c04545.

(55) Maki, H.; Tsujito, M.; Sakurai, M.; Yamada, T.; Nariai, H.; Mizuhata, M. Stabilities of the Divalent Metal Ion Complexes of a Short-Chain Polyphosphate Anion and Its Imino Derivative. J. Solut. Chem. 2013, 42 (11), 2104–2118. 10.1007/s10953-013-0099-2.

(56) Park, Y.; Malliakas, C. D.; Zhou, Q.; Gu, A. Z.; Aristilde, L. Molecular Coordination, Structure, and Stability of Metal-Polyphosphate Complexes Resolved by Molecular Modeling and X-Ray Scattering: Structural Insights on the Biological Fate of Polyphosphate. Environ. Sci. Technol. 2021, 55 (20), 14185–14193. 10.1021/acs.est.1c04782.

(57) Lorenz, B.; Schröder, H. C. Mammalian Intestinal Alkaline Phosphatase Acts as Highly Active Exopolyphosphatase. Biochim. Biophys. Acta BBA - Protein Struct. Mol. Enzymol. 2001, 1547 (2), 254–261. 10.1016/S0167-4838(01)00193-5.

(58) Geladopoulos, T. P.; Sotiroudis, T. G.; Evangelopoulos, A. E. A Malachite Green Colorimetric Assay for Protein Phosphatase Activity. Anal. Biochem. 1991, 192 (1), 112–116. 10.1016/0003-2697(91)90194-X.

(59) Hothorn, M.; Neumann, H.; Lenherr, E. D.; Wehner, M.; Rybin, V.; Hassa, P. O.; Uttenweiler, A.; Reinhardt, M.; Schmidt, A.; Seiler, J.; Ladurner, A. G.; Herrmann, C.; Scheffzek, K.; Mayer, A. Catalytic Core of a Membrane-Associated Eukaryotic Polyphosphate Polymerase. Science 2009, 324 (5926), 513–516. 10.1126/science.1168120.

(60) Christ, J. J.; Blank, L. M. Analytical Polyphosphate Extraction from Saccharomyces Cerevisiae. Anal. Biochem. 2018, 563, 71–78. 10.1016/j.ab.2018.09.021.

(61) Christ, J. J.; Blank, L. M. Saccharomyces Cerevisiae Containing 28% Polyphosphate and Production of a Polyphosphate-Rich Yeast Extract Thereof. FEMS Yeast Res. 2019, 19 (3), foz011. 10.1093/femsyr/foz011.

(62) Diaz, J. M.; Ingall, E. D. Fluorometric Quantification of Natural Inorganic Polyphosphate. Environ. Sci. Technol. 2010, 44 (12), 4665–4671. 10.1021/es100191h.

(63) Clark, J. E.; Wood, H. G. Preparation of Standards and Determination of Sizes of Long-Chain Polyphosphates by Gel Electrophoresis. Anal. Biochem. 1987, 161 (2), 280–290. 10.1016/0003-2697(87)90452-0.

(64) Manoukian, L.; Stein, R. S.; Correa, J. A.; Frigon, D.; Omelon, S. Short-Chain Polyphosphates: Extraction Effects on Migration and Size Estimation of Saccharomyces Cerevisiae Extracts with Polyacrylamide Gel Electrophoresis. ELECTROPHORESIS 2023, 44 (15–16), 1197–1205. 10.1002/elps.202300055.

(65) Chung, M.; Kim, D.; Herr, A. E. Polymer Sieving Matrices in Microanalytical Electrophoresis. Analyst 2014, 139 (22), 5635–5654. 10.1039/C4AN01179A.

(66) Sartori, A.; Barbier, V.; Viovy, J.-L. Sieving Mechanisms in Polymeric Matrices. ELECTROPHORESIS 2003, 24 (3), 421–440. 10.1002/elps.200390052.

(67) Tan, K. Y.; Herr, A. E. Ferguson Analysis of Protein Electromigration during Single-Cell Electrophoresis in an Open Microfluidic Device. The Analyst 2020, 145 (10), 3732–3741. 10.1039/c9an02553g.

(68) Pilatus, U.; Mayer, A.; Hildebrandt, A. Nuclear Polyphosphate as a Possible Source of Energy during the Sporulation of Physarum Polycephalum. Arch. Biochem. Biophys. 1989, 275 (1), 215–223. 10.1016/0003-9861(89)90366-4.

