## Supplementary information for "Near-Infrared Turn-On Fluorogenic Probe for Versatile Detection of Inorganic Polyphosphates"

#### Supplementary figures

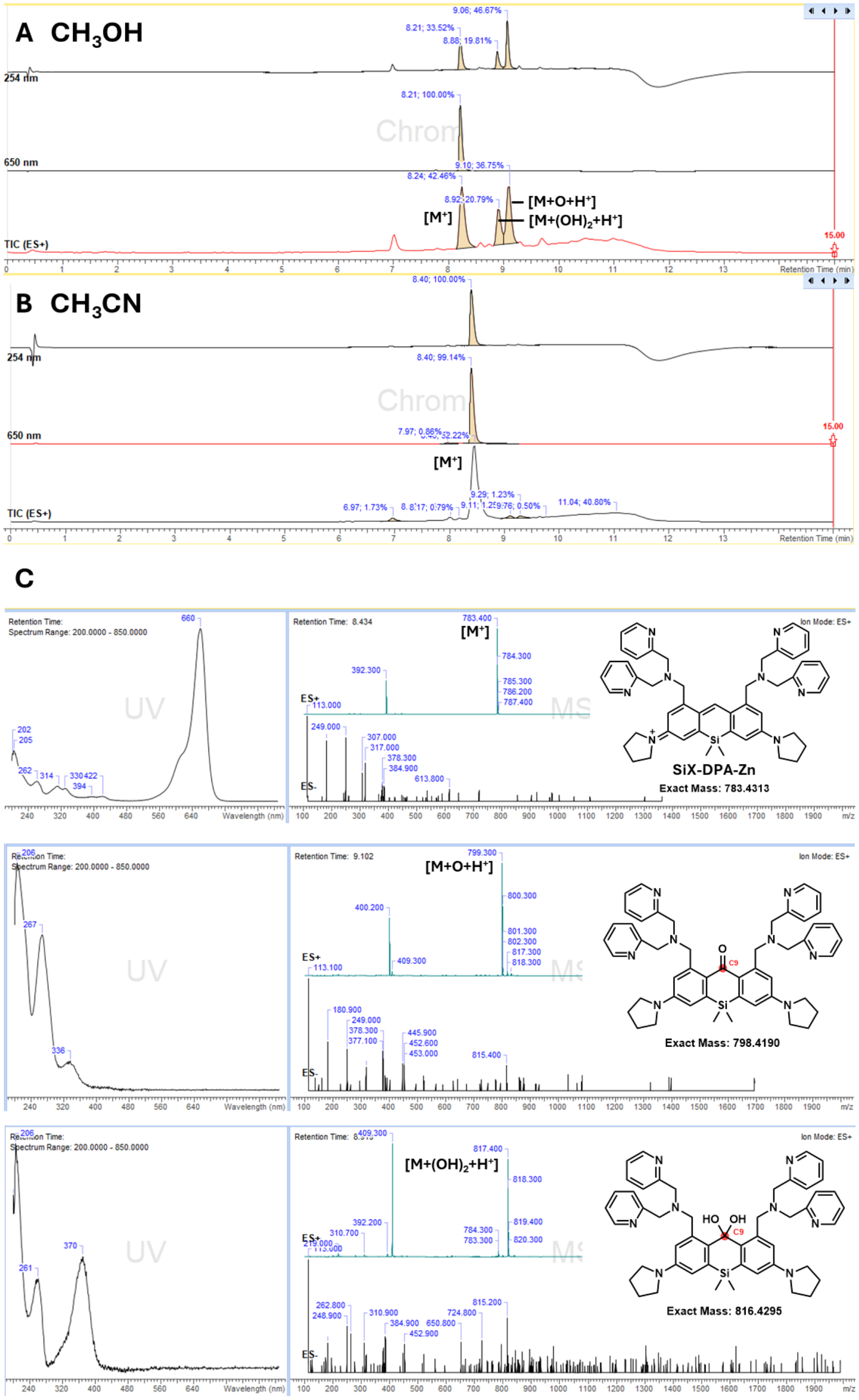

**Figure S1. SiX-DPA stability in methanol and acetonitrile.**

LC–MS spectra of purified SiX-DPA in (**A**) methanol (MeOH) and (**B**) acetonitrile (MeCN). (**C**) UV–vis absorbance and mass spectra of SiX-DPA and its oxidation byproducts in methanol. $[M]^+$ – molecular ion.

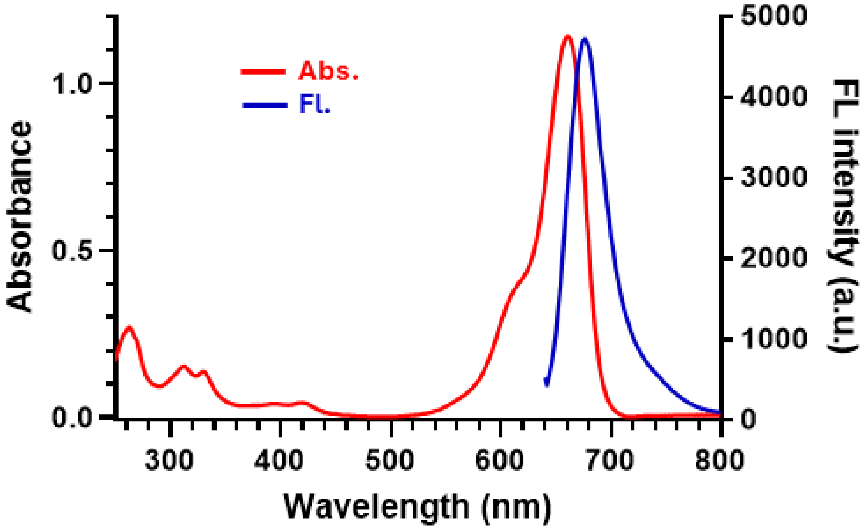

**Figure S2. Absorption and fluorescence spectra of SiX-DPA.**

Absorption spectra were recorded using 10 µM SiX-DPA, and fluorescence spectra were recorded using 1.0 µM SiX-DPA in 20 mM HEPES buffer (pH 7.5).

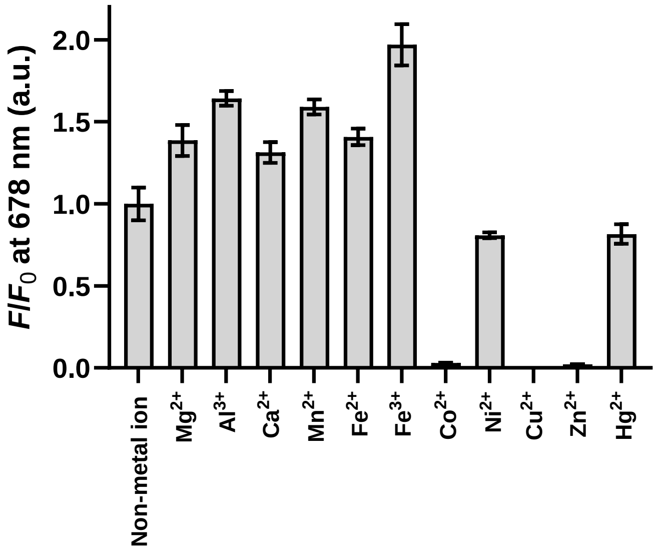

**Figure S3. Effect of various metal ions on SiX-DPA fluorescence.**

Fluorescence intensity of 1 µM SiX-DPA in 20 mM HEPES buffer (pH 7.5) supplemented with 5 µM of the indicated metal chloride. $\lambda_{ex/em}$ = 620/678 nm. $F/F_0$ represents fluorescence intensity normalized to that in the absence of any metal ions.

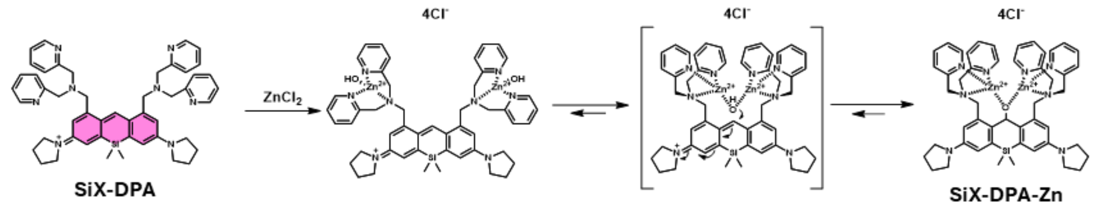

**Figure S4. Proposed quenching mechanism of SiX-DPA upon $Zn^{2+}$ complexation (SiX-DPA-Zn).**

The proposed mechanism was constructed based on the model reported in Ref 1.

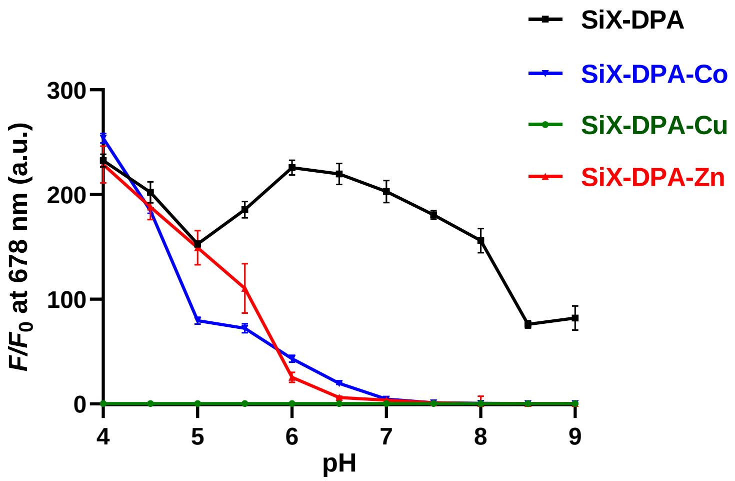

**Figure S5. pH dependency of SiX-DPA (-M) fluorescence.**

20 mM MES buffer was used for acidic conditions (pH 4.0–6.0), and 20 mM HEPES buffer was used for neutral to basic conditions (pH 6.5–9.0). $\lambda_{ex/em}$ = 620/678 nm. $F/F_0$ represents fluorescence intensity normalized to SiX-DPA-Zn at pH7.5.

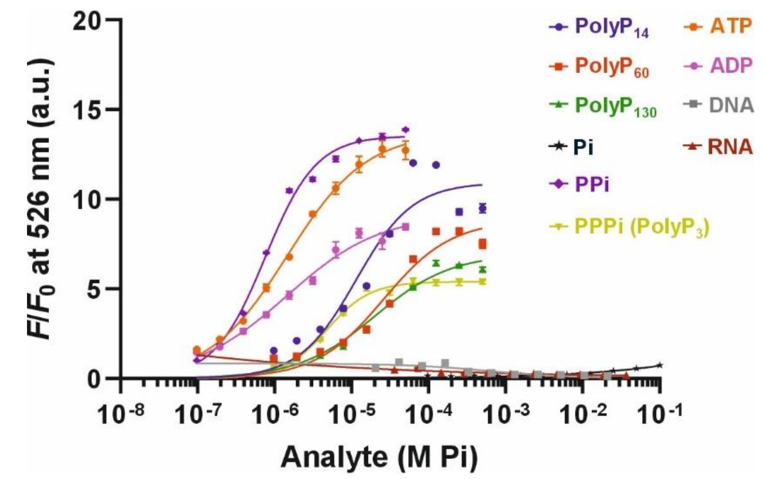

**Figure S6. Apparent affinity of X-DPA-Zn toward various phosphate-containing compounds.**

Fluorescence titration curves of 1 µM X-DPA-Zn with polyPs of different chain lengths (14, 60, and 130), PPi, PPPi (polyP$_3$), ATP, ADP, DNA, and RNA. $\lambda_{ex/em}$ = 470/526 nm. $F/F_0$ represents fluorescence intensity normalized to that in the absence of additives.

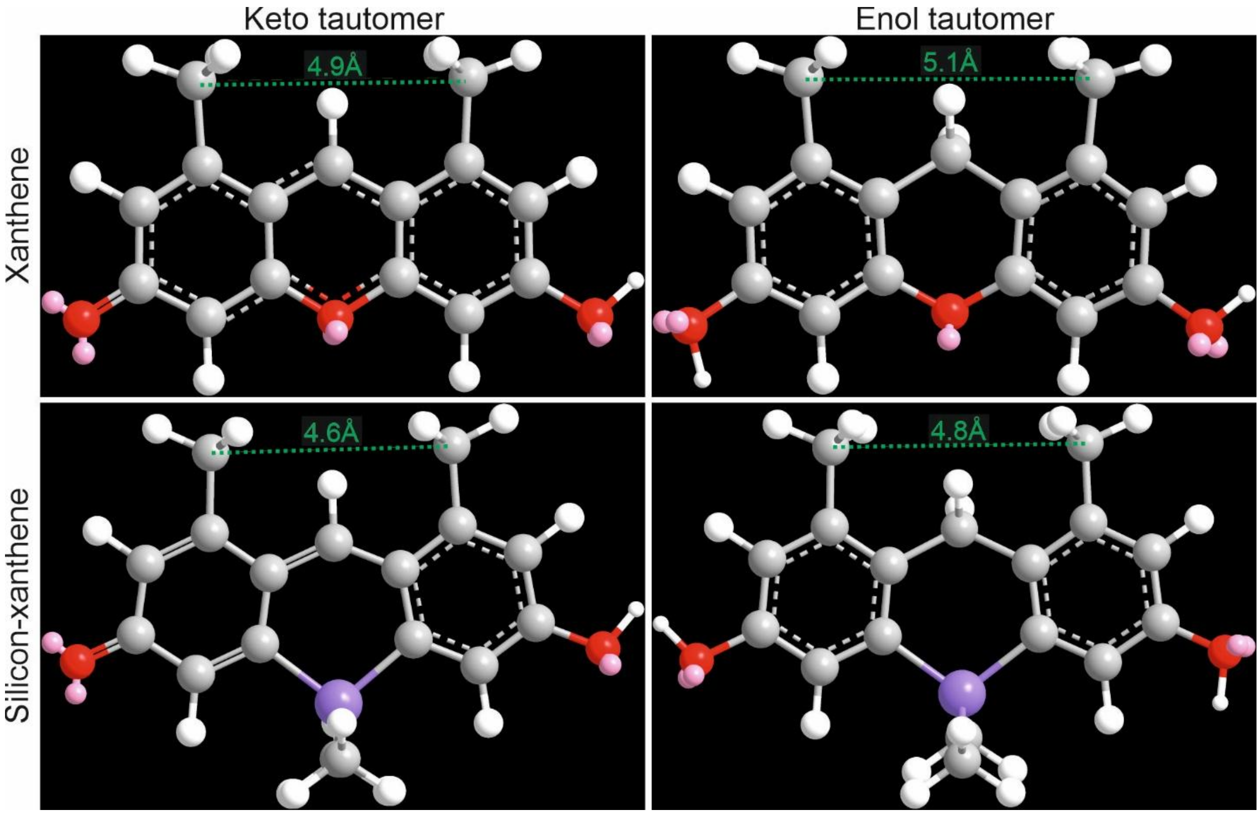

**Figure S7. Molecular models of xanthene and silicon–xanthene scaffolds.**

Molecular models were constructed in ChemDraw and geometry optimized using the Molecular Mechanics 2 (MM2) force field in Chem3D (v21.0.0.28, PerkinElmer Informatics). Interatomic distances between the substituents at 1- and 8-positions are indicated for each optimized structure.

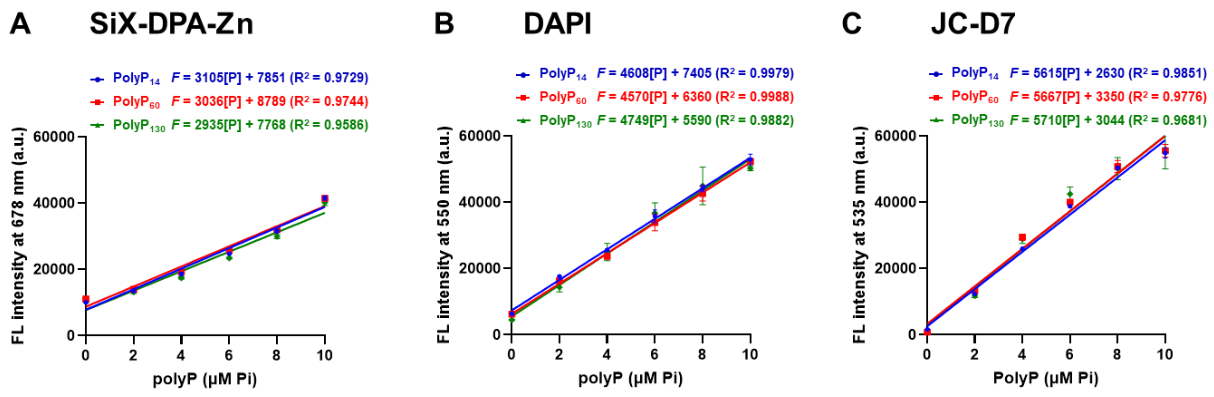

**Figure S8. Calibration curves for polyP quantification using (A) SiX-DPA-Zn, (B) DAPI, and (C) JC-D7.**

The polyP samples were diluted in 20 mM HEPES buffer (pH 7.5) supplemented with 20 µM of the indicated dye. Fluorescence was measured in a plate reader with the dye-specific $\lambda_{ex/em}$: **(A)** SiX-DPA-Zn, $\lambda_{ex/em}$ = 620/678 nm. **(B)** DAPi, $\lambda_{ex/em}$ = 415/550 nm. **(C)** JC-D7, $\lambda_{ex/em}$ = 405/535 nm. In case of SiX-DPA-Zn, the solution was additionally supplemented with 0.1% Triton X-100 in order to prevent probe aggregation. Mean ± s.d., n = 3.

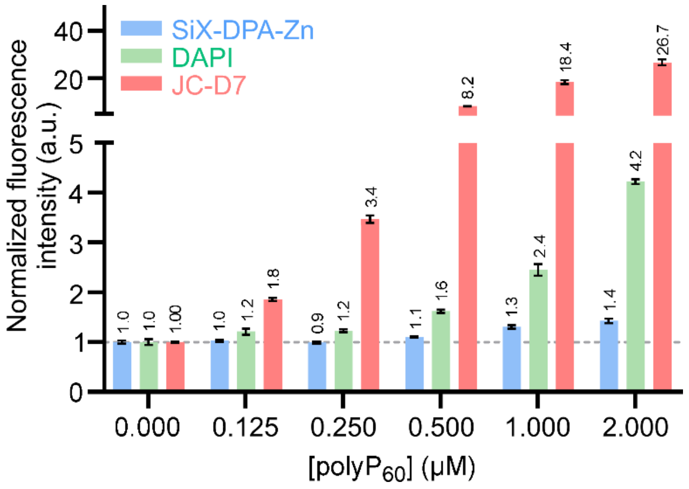

**Figure S9. Detection limits of SiX-DPA-Zn, DAPI, and JC-D7 for polyP.**

20 μM SiX-DPA-Zn, DAPI, or JC-D7 was added to polyP$_{60}$ solutions in 20 mM HEPES buffer (pH 7.5). Fluorescence was measured at $\lambda_{ex/em}$ = 620/678 nm for SiX-DPA-Zn, 415/550 nm for DAPI, and 405/535 nm for JC-D7. $F/F_0$ represents fluorescence intensity normalized to the signal in the absence of polyP.

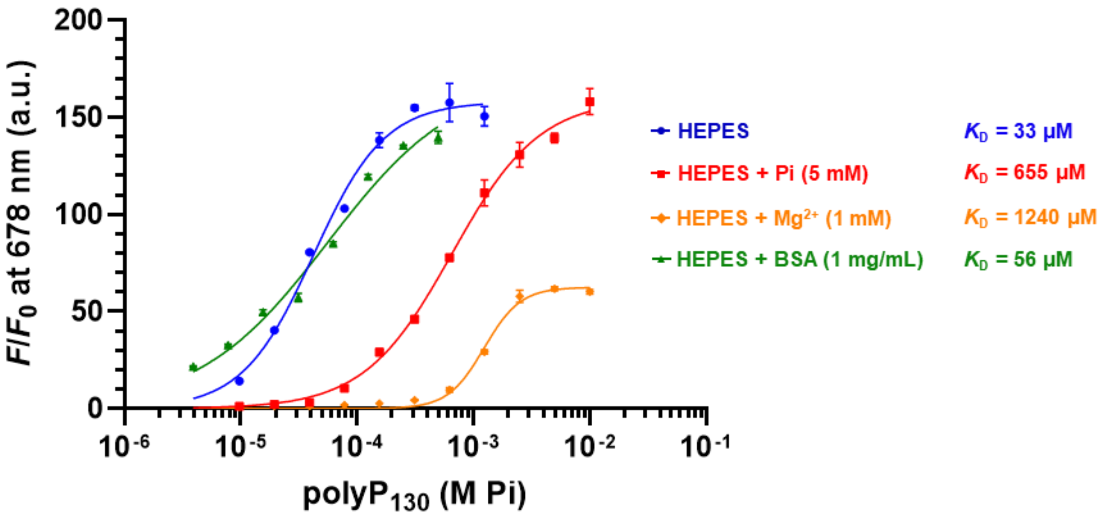

**Figure S10. Apparent affinity of SiX-DPA-Zn to polyP$_{130}$ in the presence of various competing molecules.**

Fluorescence titration curves of SiX-DPA-Zn (1.0 μM) with polyP$_{130}$ in 20 mM HEPES buffer (pH 7.5) in the absence (blue) and presence of 5 mM phosphate (red), 1 mM $Mg^{2+}$ (orange), or 1 mg/mL bovine serum albumin (green). $\lambda_{ex/em}$ = 620/678 nm. $F/F_0$ represents fluorescence intensity normalized to the signal without polyP$_{130}$.

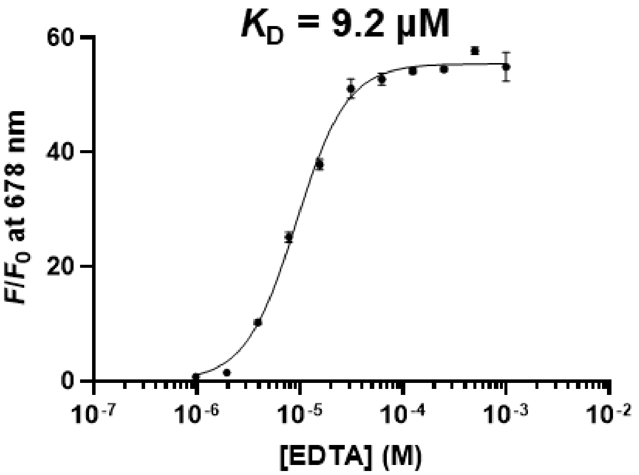

**Figure S11. Displacement of $Zn^{2+}$ from SiX-DPA-Zn by EDTA.**

Fluorescence titration curves of 1 µM SiX-DPA-Zn with different concentrations of EDTA. $\lambda_{ex/em}$ = 620/678 nm. Apparent dissociation constant ($K_D$) was determined to be 9.2 µM. $F/F_0$ represents fluorescence intensity normalized to that in the absence of EDTA.

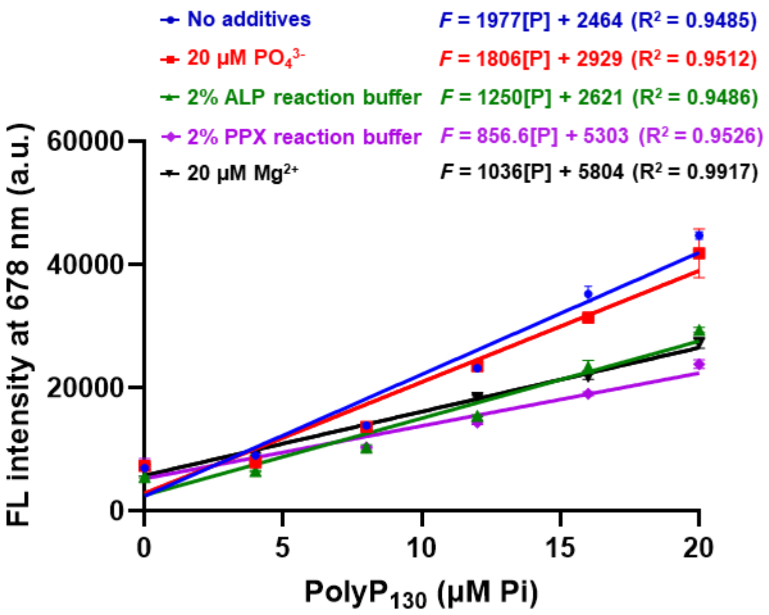

**Figure S12. Calibration curves under various conditions.**

20 μM of SiX-DPA-Zn was added to $polyP_{130}$ solutions in HEPES buffer (pH 7.5) with 0.1% TritonX-100 (blue), supplemented with the indicated additives: 20 μM $PO_4^{3-}$ (red), 2% ALP reaction buffer (20 μM $Mg^{2+}$) (green), PPX reaction buffer (20 μM $Mg^{2+}$) (purple), and 20 μM $Mg^{2+}$ (black). $\lambda_{ex/em}$ = 620/678 nm.

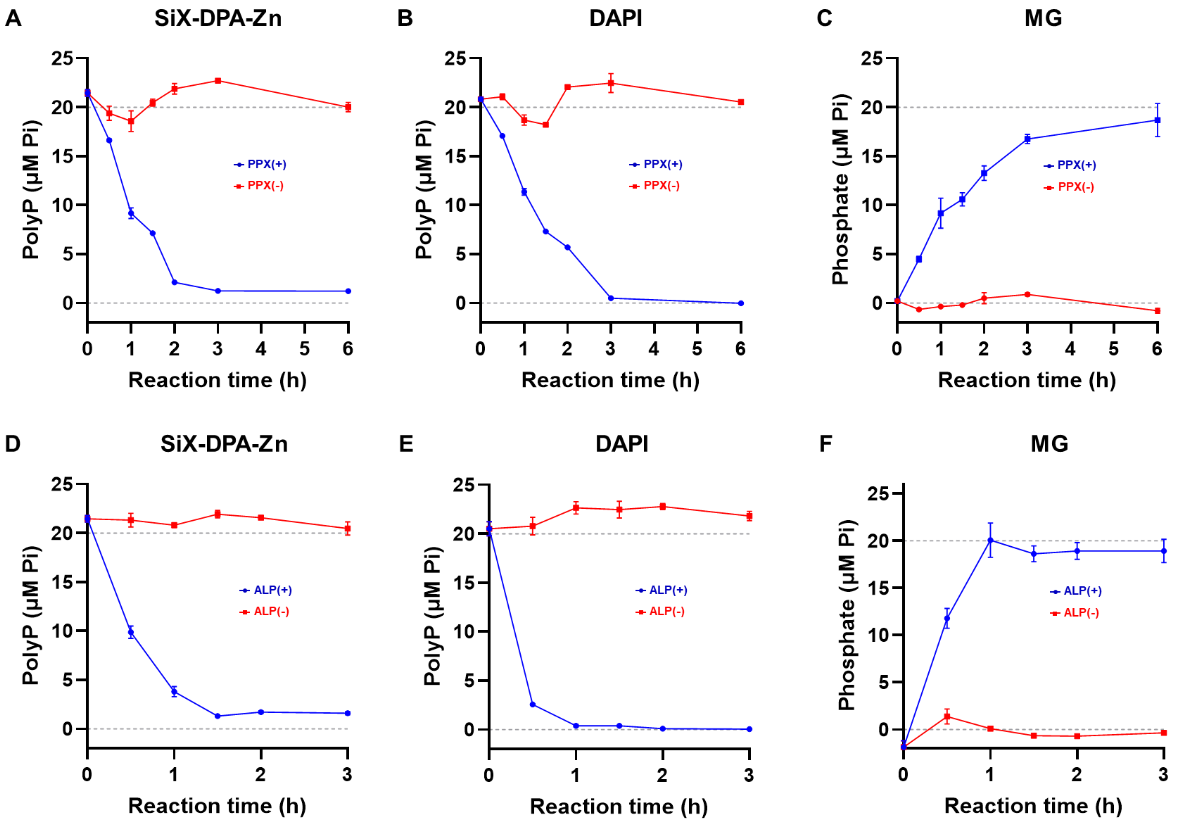

**Figure S13. Monitoring polyP digestion by *Ec*PPX and ALP using SiX-DPA-Zn, DAPI, and MG.**

Amount of $polyP_{130}$ (starting concentration 1 mM) was monitored over time during enzymatic degradation by **(A–C)** PPX (500 ng/mL) or **(D–F)** ALP (50 µg/mL). The reaction mixture was diluted 50-fold with 20 mM HEPES buffer prior to analysis. **(A, D)** 20 µM SiX-DPA-Zn or **(B, E)** 20 µM DAPI were added and fluorescence was measured immediately with the dye-specific $\lambda_{ex/em}$: SiX-DPA-Zn: $\lambda_{ex/em}$ = 620/678 nm. DAPI: $\lambda_{ex/em}$ = 415/550 nm. In case of SiX-DPA-Zn, the solution was additionally supplemented with 0.1% Triton X-100 in order to prevent probe aggregation. PolyP was quantified using calibration curves generated with 0–20 µM $polyP_{130}$ standards diluted in exactly the same buffer composition as the samples (e.g., respective enzymatic reaction buffer diluted 50-fold with 20 mM HEPES, pH 7.5) (see Figure S12 **(C, F)** The MG assay was applied to determine the concentration of released phosphate.

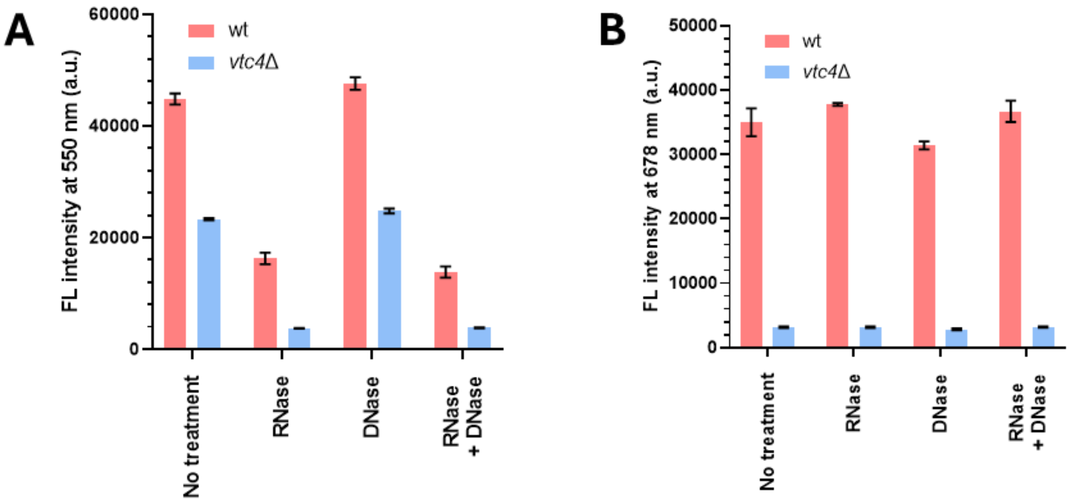

**Figure S14. Interference of nucleic acids with polyP quantification in cell extracts of *S. cerevisiae*.**

Fluorescence intensity of (**A**) 10 µM DAPI and (**B**) 10 µM SiX-DPA-Zn in cell lysates from wt and *vtc4*Δ cells. Significant DAPI fluorescence reduction upon RNase A treatment indicates potential to overestimate polyP due to co-isolated RNA. A similar observation has already been reported by other groups[2].

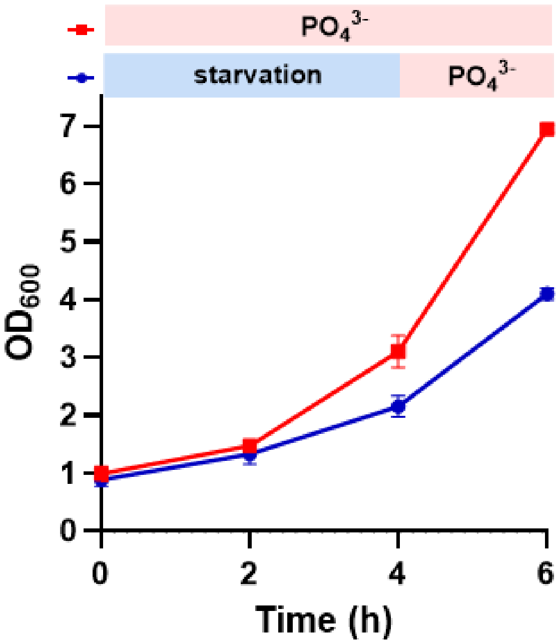

**Figure S15. Growth of wild-type *S. cerevisiae* under phosphate depletion.**

After 4 h phosphate starvation, 10 mM $KH_2PO_4$ was added to the depleted culture. Data represent three biological replicates and are shown as mean ± s.d.

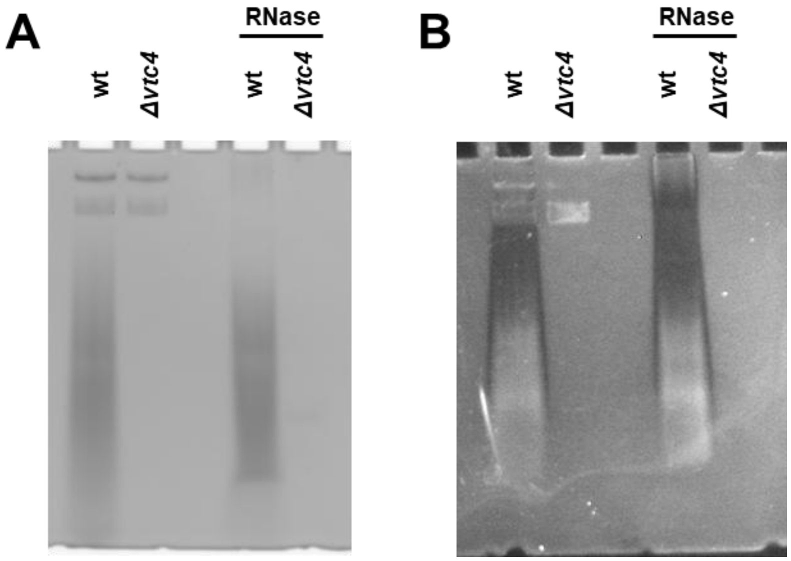

**Figure S16. Gel staining of RNA/polyP extracts from *S. cerevisiae*.**

RNA/polyP extracts were prepared from wild-type *S. cerevisiae* and the vacuolar transporter chaperone 4 (vtc4) knockout mutant. Samples were treated with 20 µg/mL RNase A for 1 h at 37 °C to remove RNA. Each sample was separated on a 30% PAGE gels and stained with (**A**) SiX-DPA-Zn and (**B**) DAPI (negative staining).

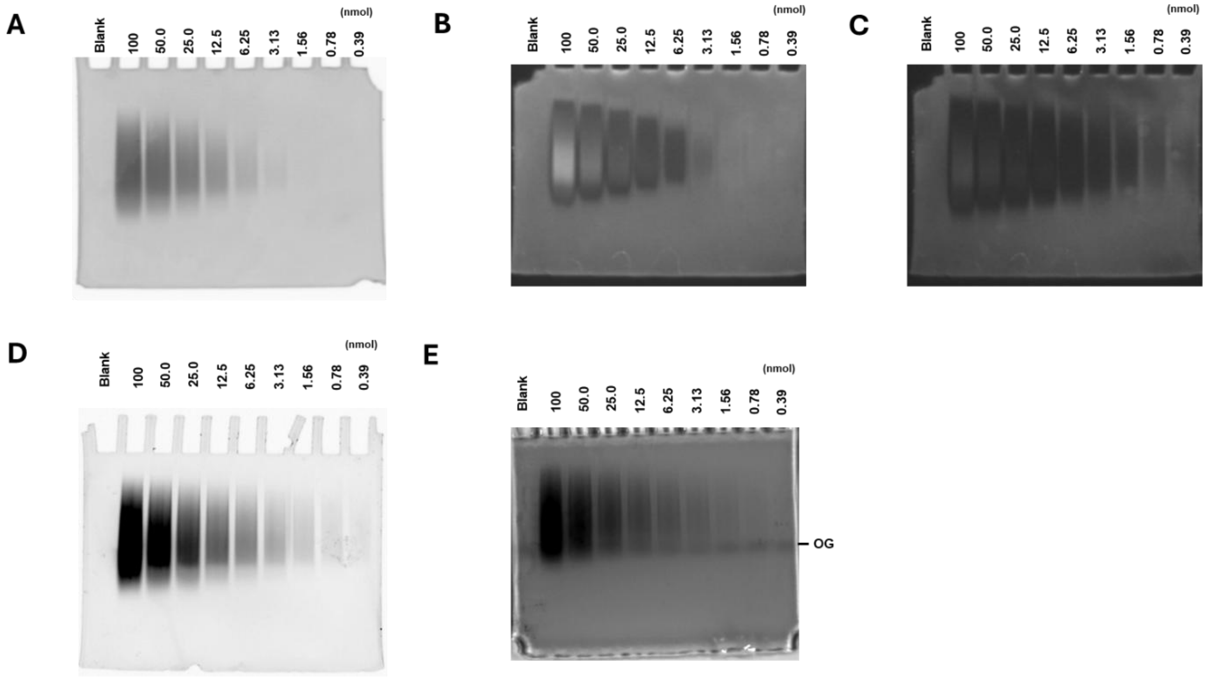

**Figure S17. Detection limits of SiX-DPA-Zn, DAPI, JC-D7, and TB-O in polyacrylamide gels.**

Different amounts of polyP$_{60}$ (0.39–100 nmol) were separated by 15% polyacrylamide gel electrophoresis and stained with (**A**) SiX-DPA-Zn, (**B**) DAPI before photobleaching, (**C**) DAPI after photobleaching, (**D**) JC-D7, and (**E**) TB-O. In Figure 4C these images were compressed vertically for visualization.

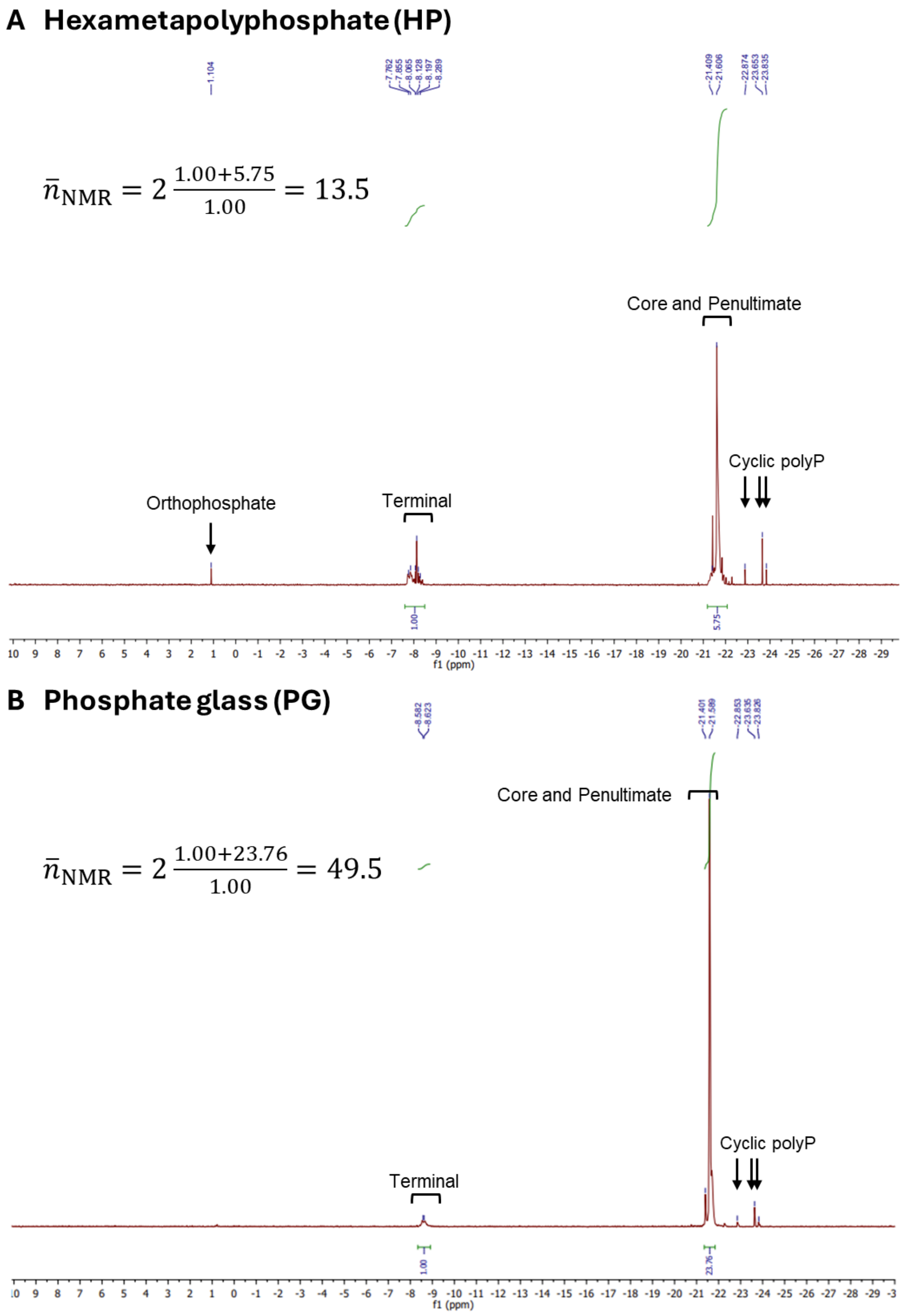

**Figure S18. 31P NMR spectra of (A) hexametaphosphate (HP) and (B) phosphate glass (PG).**

Spectral assignments were made based on Ref3. Detailed experimental conditions are provided in the Supplementary Methods section.

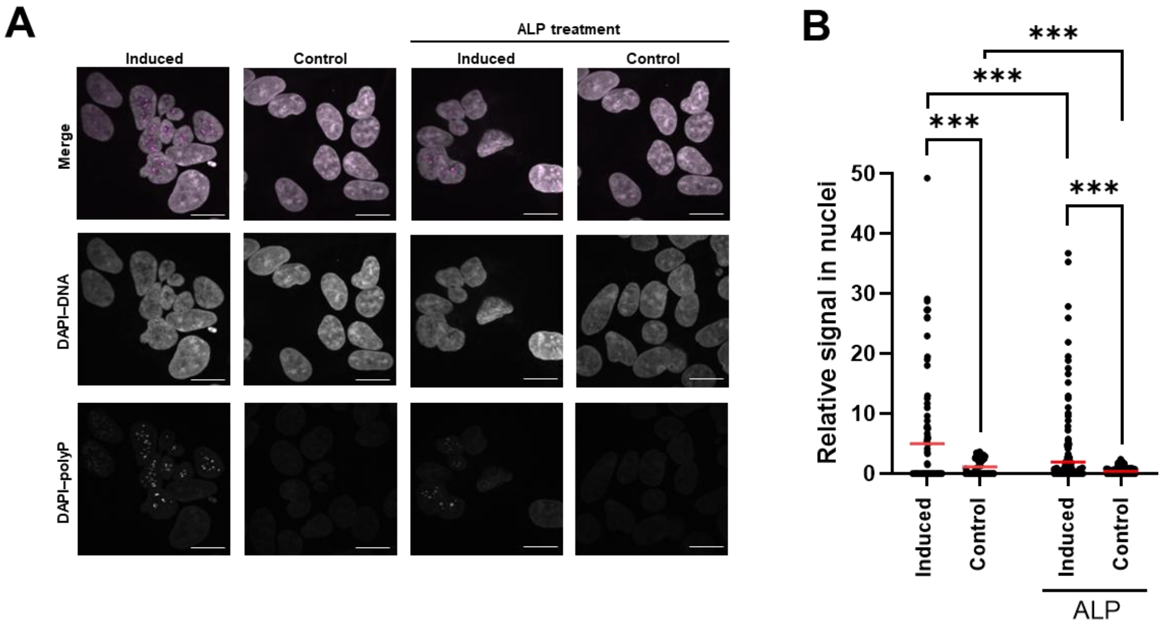

**Figure S19. Confocal fluorescence imaging of TREx-PPK cells stained with DAPI.**

**(A)** Confocal images of fixed cells induced with doxycycline for 24 h and non-induced controls. PolyP was visualized with 1.0 µM DAPI–polyP (magenta, detection at 500–550 nm), and DAPI–DNA signals (gray, detection at 435–485 nm) were recorded simultaneously. For polyP depletion, cells were treated with 1.0 mg/mL alkaline phosphatase (ALP) for 1 h before staining. Scale bar: 20 µm. **(B)** Quantification of nuclear DAPI–polyP signals. Nuclear regions were defined from DAPI–DNA staining. Signal intensities were measured per nucleus after background subtraction. Nuclei without detectable signal were plotted as 0. The numbers of analyzed nuclei were 107 (induced, no ALP), 78 (control, no ALP), 263 (induced, +ALP), and 293 (control, +ALP). Red lines indicate mean values. Statistical significance: ***$p < 0.0001$, **$p < 0.01$, *$p < 0.1$.

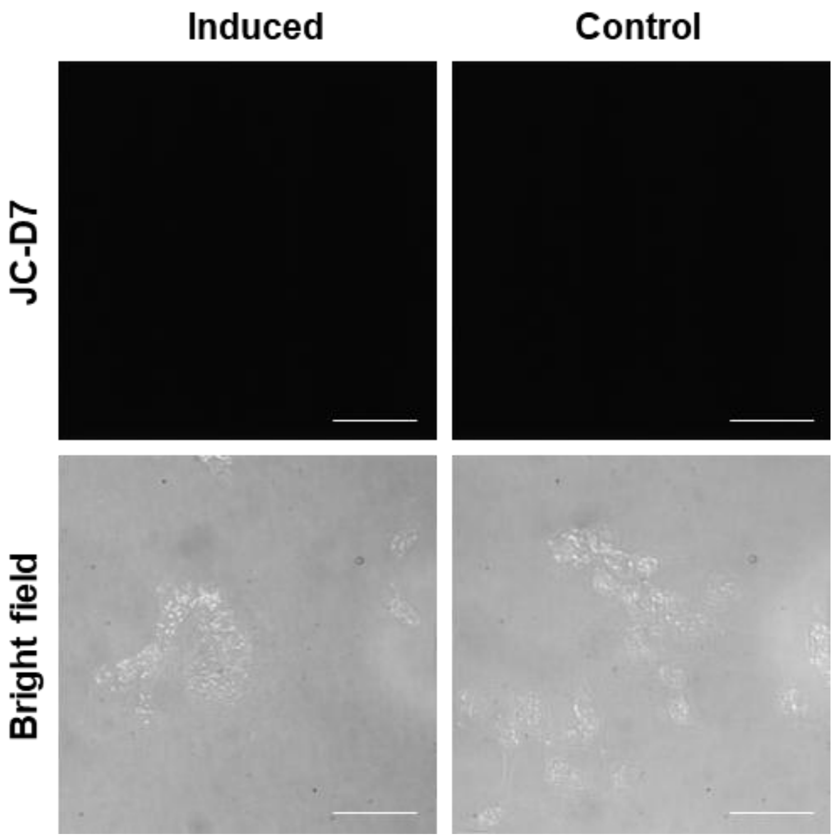

**Figure S20. Confocal fluorescence microscopy images of TREx-PPK cells stained with JC-D7.**

*Ec*PPK1 expression was induced by adding doxycycline and incubating the cells for an additional 24 hours. Control cells were non-induced cells incubated for the same period. PolyP was stained with 5 µM JC-D7 and imaged using 405 nm excitation laser and emission detection window 500–550 nm. No detectable signal was observed in the imaging field, which is consistent with the previous report[4]. Scale bar: 50 µm.

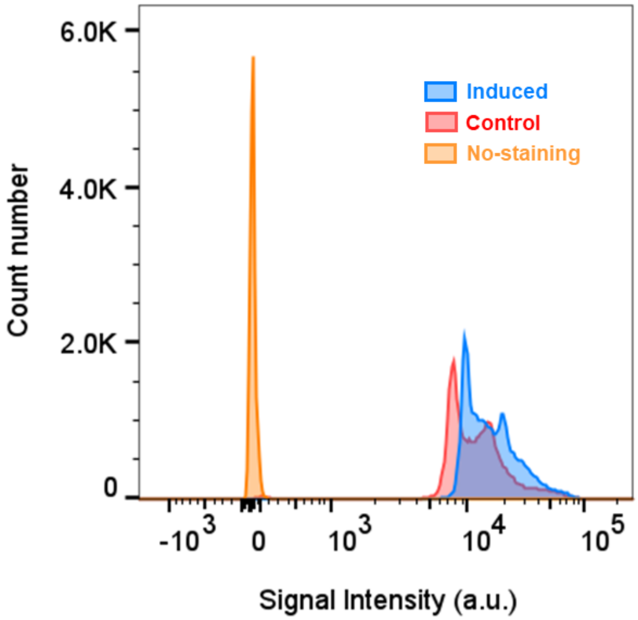

**Figure S21. Single-cell analysis of polyP using DAPI staining by flow cytometry.**

Cells were analyzed by flow cytometry using 405 nm excitation laser combined with a 502 nm LP mirror and a 510/50 nm emission filter to detect DAPI-PolyP complex fluorescence. The number of analyzed cells was 42274 (induced), 38230(control), and 18938 (no-staining).

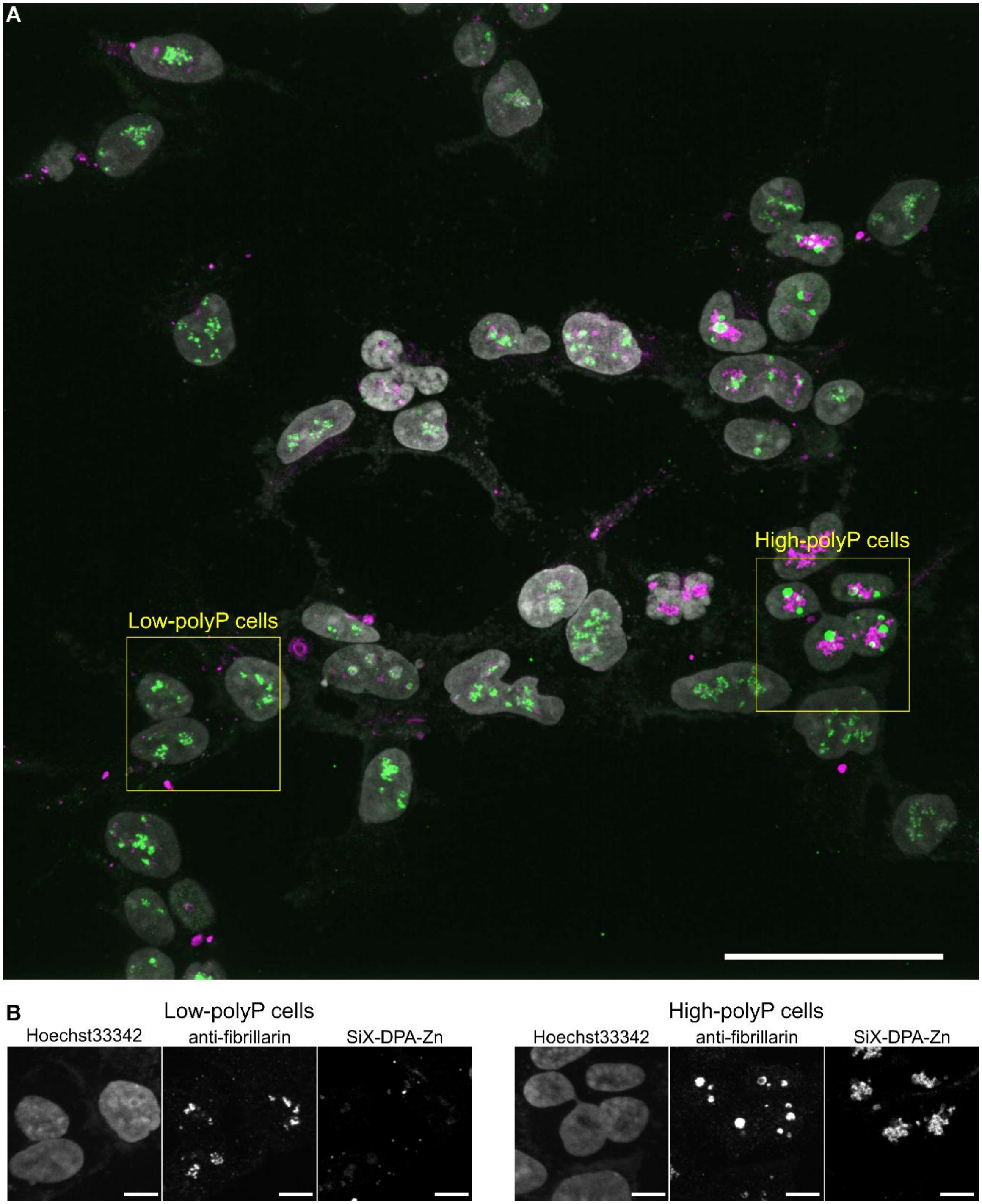

**Figure S22. Representative image of induced TREx-PPK cells co-stained for polyP and fibrillarin.**

**(A)** Large area overview, maximum intensity projection of confocal z-stack spanning the whole cell thickness. 5 µM SiX-DPA-Zn (magenta), fibrillarin immunostaining (green), Hoechst 33342 (gray). Scale bar: 50 µm. **(B)** Separated-channel images from representative low-and high-polyP cells outlined in **A**. Scale bar: 10 µm. The same insets are shown as two-channel overlays in **Figure 5D**.

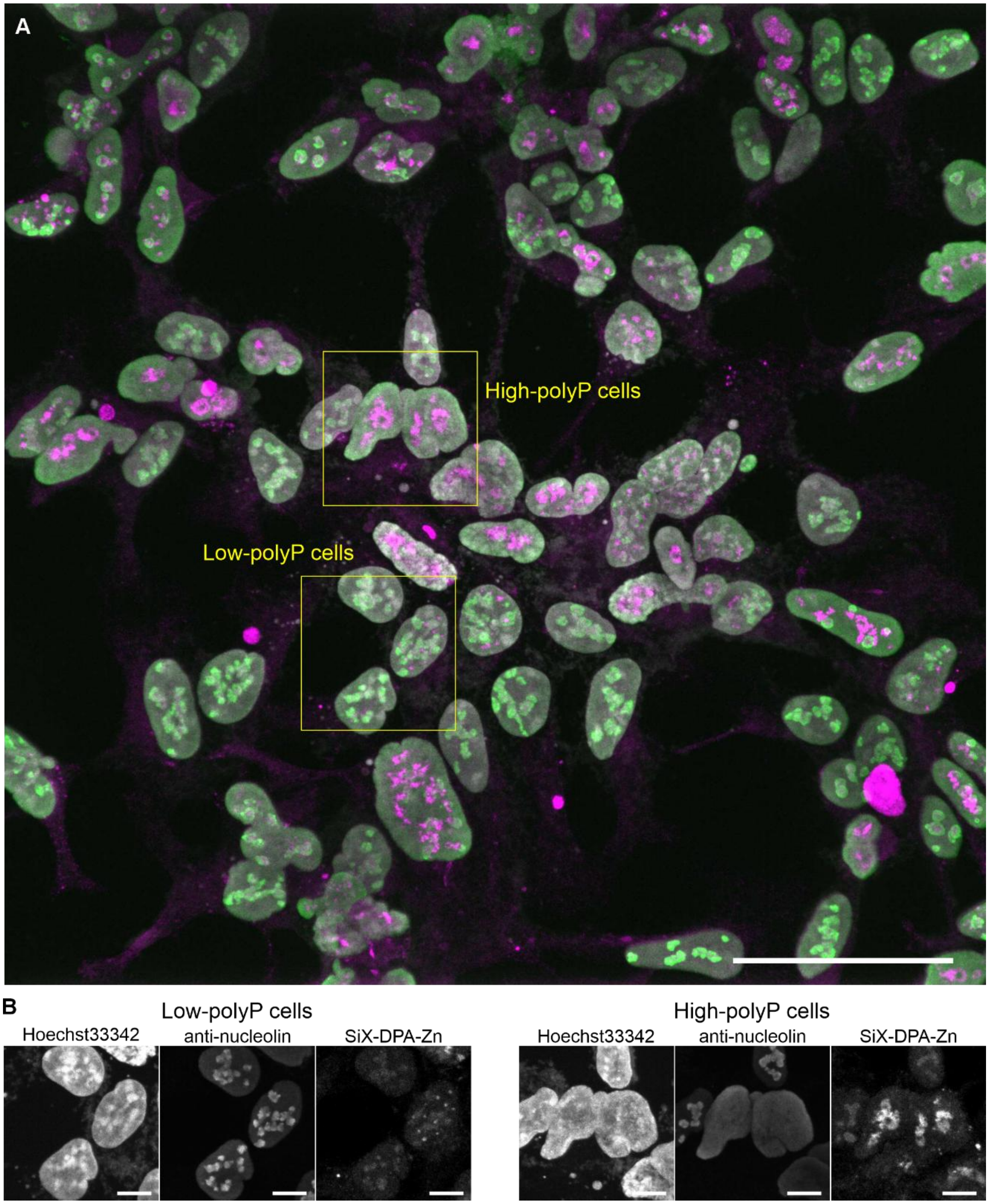

**Figure S23. Representative image of induced TREx-PPK cells co-stained for polyP and nucleolin.**

**(A)** Large area overview, maximum intensity projection of confocal z-stack spanning the whole cell thickness. 5 µM SiX-DPA-Zn (magenta), nucleolin immunostaining (green), Hoechst 33342 (gray). Scale bar: 50 µm. **(B)** Separated-channel images from representative low-and high-polyP cells outlined in **A**. Scale bar: 10 µm. The same insets are shown as two-channel overlays in **Figure 5F**.

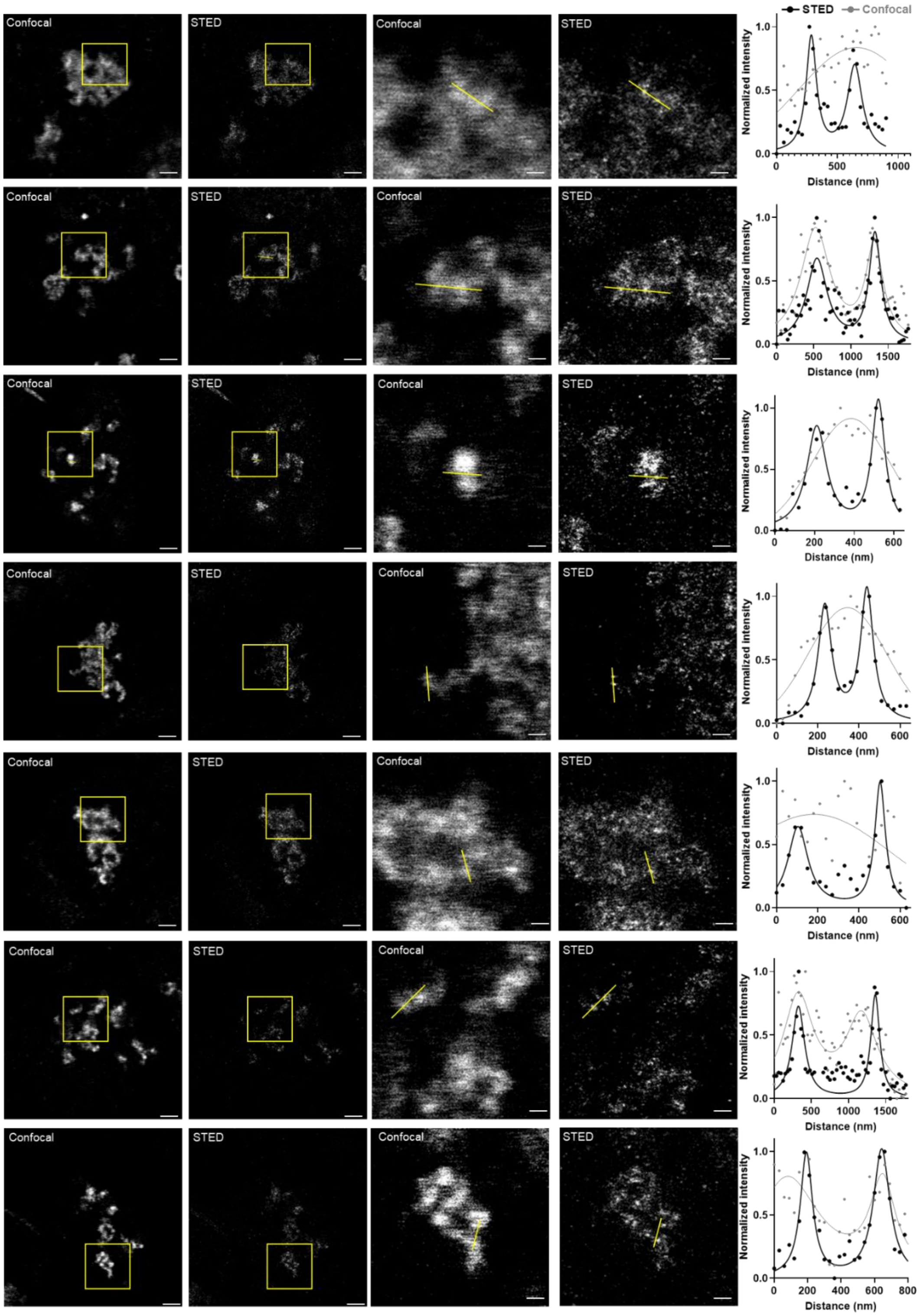

**Figure S24. Confocal and STED images of induced TREx-PPK cells stained with 5 µM SiX-DPA-Zn.**

These images correspond to **Figure 6F**. Enlarged views of the regions indicated by yellow squares. Line profiles show fluorescence intensity along the yellow line. Scale bar: 2 µm (original), 500 nm (zoom).

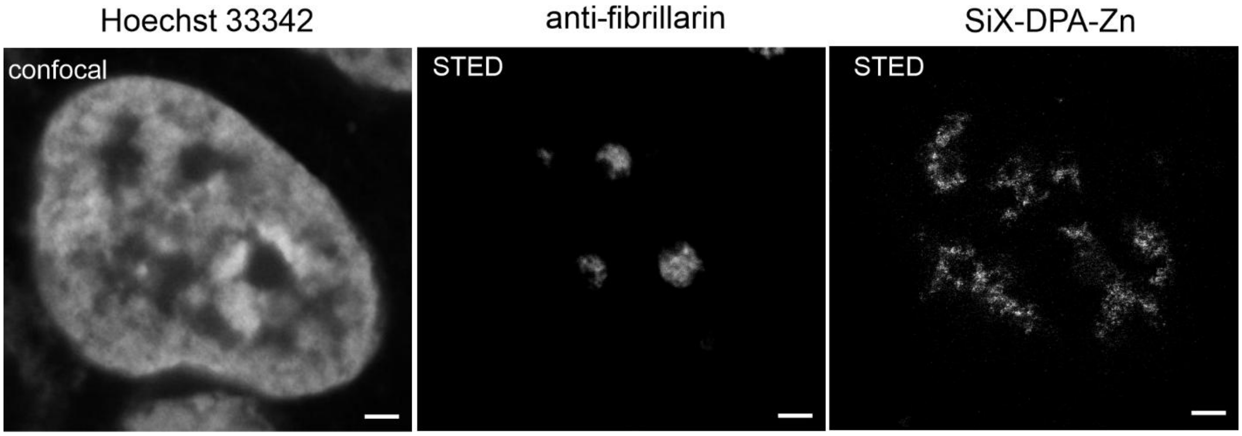

**Figure S25. Two-color STED image of doxycycline-induced TREx-PPK cells showing the distribution of polyP relative to fibrillarin.**

Intracellular polyP and nucleus were stained with 5 µM SiX-DPA-Zn and 1 µM Hoechst 33342, respectively. Fibrillarin was labeled using mouse anti-fibrillarin primary antibody and an anti-mouse nanobody-Halo-tag conjugated with 580CP-Halo. Images correspond to Figure 6G. Scale bars: 2 µm.

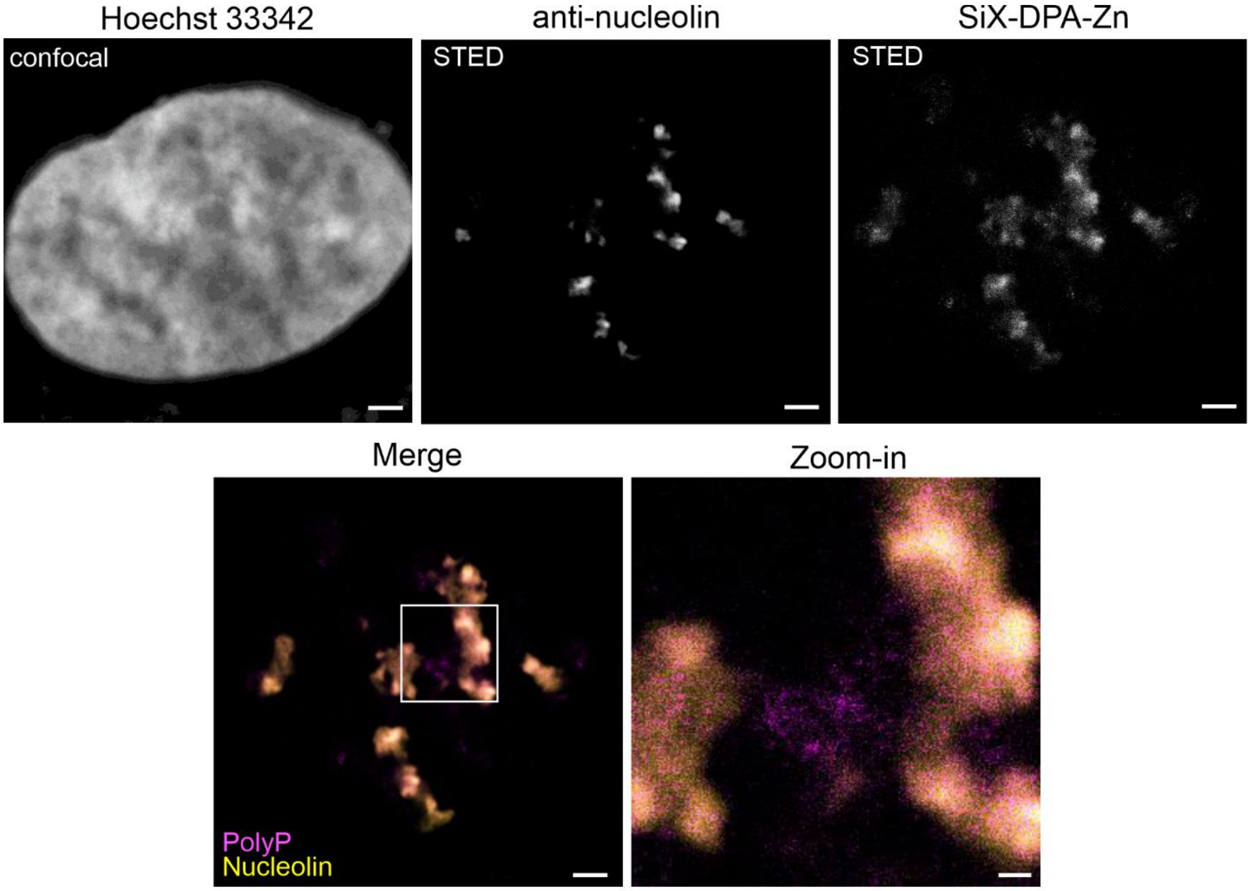

**Figure S26. Two-color STED image of doxycycline-induced TREx-PPK cells (low-polyP cell) showing the distribution of polyP relative to nucleolin.**

Cells were stained with 5 µM SiX-DPA-Zn (magenta) and 1 µM Hoechst 33342. Nucleolin was labeled using a rabbit anti-nucleolin antibody and an anti-rabbit nanobody–Halo tag conjugated with 580CP-Halo (yellow). Enlarged views of the regions indicated by white squares. Scale bars: 2 µm (original), 500 nm (zoom-in).

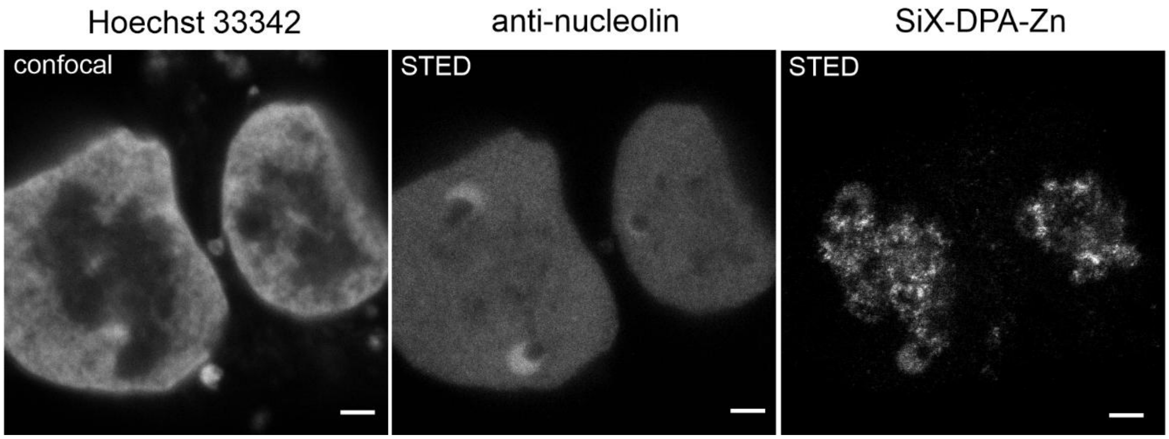

**Figure S27. Two-color STED image of doxycycline-induced TREx-PPK cells (high-polyP cell) showing the distribution of polyP relative to nucleolin.**

Cells were stained with 5 µM SiX-DPA-Zn and 1 µM Hoechst 33342. Nucleolin was labeled using a rabbit anti-nucleolin antibody and an anti-rabbit nanobody–Halo tag conjugated with 580CP-Halo. Representative raw images corresponding to Figure 6I. Scale bars: 2 µm.

### Supplementary tables

##### Table S1. Photophysical properties of SiX-DPA and SiX-DPA-Zn with $polyP_{60}$

| Sample | $\lambda_{ex,max}$ (nm) | $\varepsilon$ ($M^{-1}cm^{-1}$) | $\lambda_{em,max}$ (nm) | QY | Lifetime (ns) |
|---|---|---|---|---|---|
| SiX-DPA | 666 | 11.1 × $10^4$ | 678 | 0.235 ± 0.001 | 2.90 ($a_1$ = 3912, 34%)<br>0.27 ($a_2$ = 7722, 66%) |
| SiX-DPA-Zn with $polyP_{60}$ | 666 | 8.7 × $10^4$ | 679 | 0.083 ± 0.002 | 2.52 ($a_1$ = 3491, 22%)<br>0.23 ($a_2$ = 12130, 78%) |

SiX-DPA (1.0 µM) or SiX-DPA-Zn (1.0 µM) in 20 mM HEPES buffer (pH7.5) including 0.1% DMSO. [$PolyP_{60}$] = 1.0 mM. For fluorescence quantum yield (QY) measurement, photon was counted at $\lambda_{ex/em}$= 620 / 640–800 nm. For each lifetime measurement, 10,000 photons were collected at $\lambda_{ex/em}$= 630/680 nm. $a_i$ (i = 1,2) represents the relative contribution of each fluorescence lifetime component (see Equation S1).

##### Table S2. Apparent dissociation constants ($K_D$), Hill coefficient ($n$) and fluorescence enhancement ($F_{max}/F_0$) of SiX-DPA-Zn toward various phosphate-containing compounds

| Sample | $K_D$ (M) | | $n$ | | $F_{max}/F_0$ | | $R^2$ |
|---|---|---|---|---|---|---|---|
| | Best-fit values | 95% CI | Best-fit values | 95% CI | Best-fit values | 95% CI | |
| $polyP_{14}$ | 1.6 ×$10^{-5}$ | (1.4–1.8) ×$10^{-5}$ | 1.92 | 1.52–2.48 | 265 | 252–279 | 0.9946 |
| $polyP_{60}$ | 2.8 ×$10^{-5}$ | (2.6–3.1) ×$10^{-5}$ | 1.89 | 1.63–2.21 | 272 | 262–283 | 0.9977 |
| $polyP_{130}$ | 3.3 ×$10^{-5}$ | (3.0–3.7) ×$10^{-5}$ | 1.48 | 1.30–1.69 | 232 | 223–242 | 0.9981 |
| Pi | n.d. | n.d. | n.d. | n.d. | n.d. | n.d. | n.d. |
| PPi | 2.5 ×$10^{-6}$<br>*1.3 ×$10^{-6}$ | (2.1–2.9) ×$10^{-6}$<br>* (1.1–1.5) ×$10^{-6}$ | 1.45 | 1.23–1.98 | 222 | 209–238 | 0.9950 |
| PPPi ($polyP_3$) | 2.1 ×$10^{-5}$<br>*7.0 ×$10^{-6}$ | (1.7–2.6) ×$10^{-5}$<br>* (5.6–8.7) ×$10^{-6}$ | 2.28 | 1.52-3.58 | 203 | 184–224 | 0.9867 |
| ATP | 3.6×$10^{-3}$<br>*1.2×$10^{-3}$ | (2.3–9.5) ×$10^{-3}$<br>* (0.8–3.1) ×$10^{-3}$ | 1.26 | 0.74–2.16 | 186 | 149–248 | 0.9790 |
| ADP | 5.7×$10^{-3}$<br>*2.9×$10^{-3}$ | (4.5–7.3) ×$10^{-3}$<br>* (2.2–3.7) ×$10^{-3}$ | 2.21 | 1.43–3.64 | 264 | 236–285 | 0.9849 |
| DNA | n.d. | n.d. | n.d. | n.d. | n.d. | n.d. | n.d. |
| RNA | 6.9×$10^{-3}$ | n.d. | n.d. | n.d. | 185 | 168–202 | 0.9908 |

*$K_D$ value determined based on molar concentration. 95% CI: 95% confidence interval. n.d.: not determined due to negligible fluorescence changes.

**Table S3. Apparent dissociation constants ($K_D$), Hill coefficient ($n$) and fluorescence enhancement ($F_{max}/F_0$) of X-DPA-Zn toward various phosphate-containing compounds**

| Sample | $K_D$ (M) | | $n$ | | $F_{max}/F_0$ | | $R^2$ |
|---|---|---|---|---|---|---|---|
| | Best-fit values | 95% CI | Best-fit values | 95% CI | Best-fit values | 95% CI | |
| polyP$_{14}$ | $1.1\times10^{-5}$ | $(0.5–3.7)\times10^{-5}$ | 1.18 | 0.50–2.90 | 13 | 11–18 | 0.8923 |
| polyP$_{60}$ | $2.6\times10^{-5}$ | $(1.4–10.5)\times10^{-5}$ | 0.99 | 0.53–1.85 | 11 | 8.7–15 | 0.9552 |
| polyP$_{130}$ | $2.0\times10^{-5}$ | $(1.2–6.1)\times10^{-5}$ | 0.88 | 0.53–1.40 | 8.4 | 7.1–11 | 0.9684 |
| Pi | n.d. | n.d. | n.d. | n.d. | n.d. | n.d. | n.d. |
| PPi | $7.6\times10^{-7}$<br>*$3.8\times10^{-7}$ | $(6.3–9.2)\times10^{-7}$<br>* $(3.2–4.6)\times10^{-7}$ | 1.32 | 1.06–1.66 | 16 | 15–17 | 0.9936 |
| PPi** | $2.5\times10^{-8}$ | --- | --- | --- | 55 | --- | --- |
| PPPi (polyP$_3$) | $4.7\times10^{-6}$<br>*$1.6\times10^{-6}$ | $(3.9–5.8)\times10^{-6}$<br>* $(1.3–1.9)\times10^{-6}$ | 1.53 | 1.18–2.03 | 6.5 | 6.3–6.8 | 0.9884 |
| ATP | $1.5\times10^{-6}$<br>*$4.9\times10^{-7}$ | $(1.2–1.8)\times10^{-6}$<br>* $(4.1–6.0)\times10^{-7}$ | 0.85 | 0.75–0.96 | 16 | 15–17 | 0.9974 |
| ATP** | $7.7\times10^{-7}$ | --- | --- | --- | 33 | --- | --- |
| ADP | $1.4\times10^{-6}$<br>*$7.1\times10^{-7}$ | $(0.9–3.0)\times10^{-6}$<br>* $(4.5–10.1)\times10^{-7}$ | 0.71 | 0.52–0.94 | 11 | 9.7–13 | 0.9862 |
| ADP** | $5.9\times10^{-7}$ | --- | --- | --- | 18 | --- | --- |
| DNA | n.d. | n.d. | n.d. | n.d. | n.d. | n.d. | n.d. |
| RNA | n.d. | n.d. | n.d. | n.d. | n.d. | n.d. | n.d. |

*$K_D$ value determined based on molar concentration. **The value provided by previous study[1].

95% CI: 95% confidence interval. n.d.: not determined due to negligible fluorescence changes.

**Table S4. Electrophoretic mobility ($R_f$) and chain length of polyPs ($n$) in each gel**

| $n$ | $R_f$ | | | |
|---|---|---|---|---|
| | gel1 | gel2 | gel3 | average |
| **3** | 2.81 | 2.76 | 2.48 | 2.68 |
| **4** | 2.60 | 2.40 | 2.27 | 2.42 |
| **5** | 2.31 | 2.16 | 2.05 | 2.18 |
| **6** | 2.15 | 2.01 | 1.91 | 2.02 |
| **7** | 2.02 | 1.89 | 1.78 | 1.90 |
| **14** | 1.37 | 1.21 | 1.09 | 1.22 |
| **60** | 0.63 | 0.65 | 0.58 | 0.62 |
| **130** | 0.33 | 0.35 | 0.30 | 0.33 |

#### Table S5. Calibration curves obtained from each gel

| | Calibration curve | Range | $R^2$ |
|---|---|---|---|
| gel1 | $\log R_f = -0.3949 \log n + 0.6417$ | $3 \leq n \leq 7$ | 0.9945 |
| | $R_f = -1.086 \log n + 2.601$ | $14 \leq n \leq 130$ | 0.9949 |
| gel2 | $\log R_f = -0.4494 \log n + 0.6527$ | $3 \leq n \leq 7$ | 0.9976 |
| | $R_f = -0.8871 \log n + 2.226$ | $14 \leq n \leq 130$ | 0.9999 |
| gel3 | $\log R_f = -0.3893 \log n + 0.5837$ | $3 \leq n \leq 7$ | 0.9954 |
| | $R_f = -0.8201 \log n + 2.033$ | $14 \leq n \leq 130$ | 0.9999 |

#### Table S6. Comparison of fluorescent chemosensors for polyPs

| Chemical structure | | 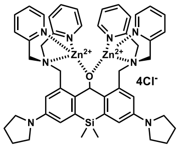 | 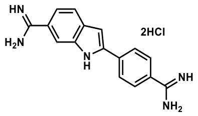 | 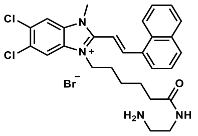 |
|---|---|---|---|---|
| Name | | SiX-DPA-Zn | DAPI | JC-D7 |
| Photophysical properties | $\lambda_{ex,\ max}$ (nm)* | 666 | 380 | 366 |
| | $\lambda_{em,\ max}$ (nm)* | 679 | 536 | 504 |
| Quantitative analysis | Working polyP concentration (μM) | 2–20 | 2–20 (0.75–3.0)[ref5] | 2–20 (0.5–36)[ref6] |
| | Detection limit (μM) | 0.5 | <0.125 | <0.125 |
| | Cell lysate from *S.cerevisiae* | ✓ | Background signal from RNA | × |
| Gel staining | Detection of short-chain polyP (<15P unit) | ✓ | × | × |
| | **Detection limit (nmol) | 1.6 | <0.4*** (0.004)***[ref7] | <0.4 |
| Intracellular detection | Confocal | ✓ | ✓ | × |
| | Flow cytometry | ✓ | × | – |
| | STED | ✓ | – | – |

* Data were collected using chemosensors at 10 μM in 20 mM HEPES (pH 7.4) with 1 μM $polyP_{60}$. **Detection limit for gel staining was determined using 15% polyacrylamide loaded with various concentrations of $polyP_{60}$ gel and stained with 10 μM chemosensors. ***Data were collected using negative DAPI staining method. ✓: applicable, ×: not applicable, –: not examined.

#### Supplementary video

**Video S1. Rotating maximum-intensity projection of a 3D confocal image stack of doxycycline-induced TREx-PPK cells.**

Intracellular polyP was stained with SiX-DPA-Zn (5.0 µM, red), and nuclei were counterstained with Hoechst 33342 (1.0 µM, blue).

#### Supplementary methods

##### Materials and instruments for synthesis

All chemical reagents used for the synthesis were purchased from Merck (Darmstadt, Germany; formerly Sigma-Aldrich, St. Louis, MO, USA), BLD Pharmatech GmbH (Reinbek, Germany), Carbolution Chemicals GmbH (St. Ingbert, Germany), and Tokyo Chemical Industry Co., Ltd. (Tokyo, Japan), and were used without further purification. Reaction progress was monitored by thin-layer chromatography (TLC) on silica gel 60 $F_{254}$ aluminum sheets (Merck). Purification was performed by normal-phase flash column chromatography (Isolera™ One; Biotage, Uppsala, Sweden) with Sfär Silica HC Columns (Biotage). SiX-DPA was purified by preparative reverse-phase high-pressure liquid chromatography (HPLC) using Agilent1260/1290 Infinity II system (Agilent Technologies, CA, USA) composed of an autosampler (Cat. No. G7158B), binary pump (Cat. No. G7161B), multiple wavelength detector (Cat. No. G7165A). Sample separation was performed using a Pursuit 10 C18 preparative column (10 µm, 250 × 50 mm; Agilent Technologies) with mobile phase A consisting of 0.1% trifluoroacetic acid in water and mobile phase B consisting of acetonitrile. A linear gradient was applied from 70:30 (A/B) to 0:100 over 25 min at a flow rate of 150 mL/min.

The purity of the synthesized compounds was confirmed by both thin-layer chromatography (TLC) and liquid chromatography–mass spectrometry (LC–MS) using an Agilent 1260 Infinity II system (Agilent Technologies). The system was composed of a binary pump (G7112B), an autosampler (G7129A), temperature-controlled column compartment (G7116A), diode array detector WR (G7115A) and 6100 series quadruple LC/MSD XT (G6135B) with API electrospray. Sample separation was performed using an Ascentis® Express AQ–C18 UHPLC column (2 µm, 5 cm × 2.1 mm; Merck) with mobile phase A consisting of 25 mM ammonium formate (pH 3.5) in water and mobile phase B consisting of methanol. A linear gradient was applied from 60:40 (A/B) to 0:100 over 6 min at a flow rate of 0.4 mL/min, with the column maintained at 40 °C.

High resolution mass spectra (HRMS) were recorded on a micrOTOF spectrometer (Bruker Daltonics, MA, USA) equipped with electrospray ionization (ESI) source (Apollo; Bruker Daltonics) and direct injector with LC autosampler (RR 1200; Agilent Technologies).

$^{1}$H NMR (400 MHz) and $^{13}$C NMR (101 MHz) spectra were recorded on an Agilent 400–MR spectrometer. Chemical shifts ($\delta$) were referenced to residual non-deuterated solvent signals as internal standards: $CDCl_3$ ($^{1}$H NMR: $\delta$ = 7.26 ppm; $^{13}$C NMR: $\delta$ = 77.16 ppm)[8]. Signal multiplicities are reported as follows: s = singlet, d = doublet, t = triplet, q = quartet, m = multiplet or overlap of non-equivalent resonances, br = broad signal. Coupling constants (*J*) are given in Hz.

**Measurement of absolute quantum yield of SiX-DPA**

The absolute fluorescence quantum yield (QY) of SiX-DPA was measured using Quantaurus-QY spectrometer (Cat. No. C11347; Hamamatsu Photonics, Shizuoka, Japan). Measurements were performed on solutions of SiX-DPA (1 µM) in 20 mM HEPES buffer (pH 7.5) using 3 mL quartz cuvettes (Cat. No. A10095-02). Photons were collected in the 640–800 nm range following excitation at 619 nm.

**Measurement of fluorescence lifetime of SiX-DPA**

The fluorescence lifetime ($\tau$) of SiX-DPA was measured using a Quantaurus-Tau fluorescence lifetime spectrometer (Cat. No. C11367; Hamamatsu Photonics). The fluorescence decay profile was recorded over a 53 ns interval following excitation. Measurements were performed on 1 µM solutions of SiX-DPA in 20 mM HEPES buffer (pH 7.5) using 3 mL quartz cuvettes (Cat. No. A10095-02). Each measurement was terminated when 10,000 peak photons were collected at an excitation/emission wavelength of 630/680 nm. The fluorescence decay data were processed and fitted using a sum of exponential functions via nonlinear least-squares deconvolution (Equation S1). $a_i$ denotes the amplitude (or pre-exponential factor) corresponding to each fluorescence lifetime component.

$$I(t) = \sum a_i \exp \frac{t}{\tau_i} \qquad \text{Equation S1}$$

**Measurement of fluorescence intensity of SiX-DPA and SiX-DPA-M (M = $Zn^{2+}$, $Cu^{2+}$, and $Co^{2+}$)**

The fluorescence intensity of SiX-DPA was measured using a microplate reader (Spark 20M; TECAN Group Ltd., Zürich, Switzerland) in a 96-well half-area microplate (Cat. No. 4580, Corning, NY, USA). The sample volume in each well was fixed at 100 µL. Prior to measurement, the plate was shaken for 5 seconds to homogenize the solution. Fluorescence was recorded at an excitation wavelength of 620 nm and an emission wavelength of 678 nm at 25 °C.

**Preparation of SiX-DPA-metal complexes (SiX-DPA-Co, SiX-DPA-Cu, and SiX-DPA-Zn)**

SiX-DPA-metal complexes (SiX-DPA-Co, SiX-DPA-Cu, and SiX-DPA-Zn) were prepared by adding 2.5 equivalents of 200 mM $CoCl_2$, $CuCl_2$, and $ZnCl_2$ aqueous solution to 1- or 10-mM SiX-DPA in DMSO (Cat. No. D2653, Merck), respectively. The samples were stored at –80 °C in the dark until use.

**Preparation of phosphate-containing samples (polyPs, PPi, ATP, ADP, DNA, and RNA)**

PolyPs ($polyP_{14}$, $polyP_{60}$, and $polyP_{130}$) were kindly provided by RegeneTiss, Inc (Nagano, Japan). Each sample was dissolved in $dH_2O$ as a sodium polyphosphate. The concentration of polyP was determined by using the Malachite Green (MG) phosphate assay after complete digestion into phosphate by incubation with 1 M $HClO_4$ for 30 min at 100 °C. After digestion, the sample solution was diluted 200–2,000-fold with $dH_2O$ to adjust the final concentration to 10–40 µM, Malachite Green was added, and the absorbance at 620 nm was recorded according to the manufacturer's specifications. PPi (Cat. No. P9146, Sigma-Aldrich) was used as sodium pyrophosphate decahydrate. The concentration of PPi was calculated from the molecular weight ($Na_4P_2O_7 \cdot 10H_2O$ = 446.05). ATP (Cat. No. A2383; Merck) and ADP (Cat. No. A2754; Merck) were purchased as sodium adenosine triphosphate and sodium diphosphate, respectively. The concentration of ATP and ADP were determined from the absorbance at 260 nm, using a molar extinction coefficient of 15,400 $M^{-1}$ $cm^{-1}$. The concentrations of DNA from calf thymus (Cat. No. D8515; Merck) and RNA from yeast (Cat. No. 10109223001; Roche, Basel, Switerland) were determined using extinction coefficients of 0.020 $(\mu g/mL)^{-1}$ $cm^{-1}$ for DNA and 0.025 $(\mu g/mL)^{-1}$ $cm^{-1}$ for RNA at 260 nm, respectively. The concentrations of all phosphate-containing samples are expressed as concentration of phosphate units ([Pi]). The [Pi] values for DNA and RNA were calculated by approximating the average molecular weight of nucleotides as 330 Da. All samples were dissolved in $dH_2O$ and stored at –80 °C until use.

**Determination of $K_D$, $n$, and $F_{max}/F_0$ for phosphate-containing compounds**

The fluorescence intensity of SiX-DPA-Zn and X-DPA-Zn was recorded using multimode microplate reader (Spark 20M; TECAN Group Ltd., Zürich, Switzerland) in a 96-well half-area microplate (Cat. No. 4580, Corning, NY, USA). The concentrations of SiX-DPA-Zn and X-DPA-Zn were adjusted to 1.0 µM in 20 mM HEPES buffer (pH 7.5) containing 1% DMSO. Analytical parameters indicating the affinity for phosphate-containing samples, including $K_{\mathrm{D}}$ (apparent dissociation constant), $n$ (Hill coefficient), and $F_{\mathrm{max}}$ (saturated fluorescence intensity) were determined from the fluorescence intensity at 678 nm (excitation at 600 nm) for SiX-DPA-Zn and 524 nm (excitation at 470 nm) for X-DPA-Zn in response to phosphate concentration ([Pi]) by fitting the data to the Hill equation (Equation S2).

$$F = F_{\mathrm{max}} \frac{[\mathrm{P}]^n}{K_{\mathrm{D}}^n + [\mathrm{P}]^n} \qquad \text{Equation S2}$$

$F_0$ was determined by measuring the fluorescence intensity of the probe without containing any phosphates in 20 mM HEPES buffer.

**Fluorescence titration for the quantification of polyP concentration**

To quantify the concentration of polyPs, fluorescence titration was performed using SiX-DPA-Zn as a polyP-sensitive probe. A final concentration of 20 μM SiX-DPA-Zn, 20 μM DAPI (Cat. No. D9542, Merck), or 20 μM JC-D7 (Cat. No. V22869, InvivoChem, IL, USA) was added to polyP solutions of varying concentrations (0, 2, 4, 6, 8, and 10 μM) prepared in 20 mM HEPES buffer (pH 7.5). For experiments requiring calibration curves generated using SiX-DPA-Zn (e.g. Figure 3B, 3C, 3E, 3F, S8, S12, and S13), the buffer was supplemented with 0.1% Triton X-100 (Cat. No. 1.08603.1000, Merck) to prevent intermolecular aggregation of the probe.

**Monitoring polyphosphatase reactions**

Alkaline phosphatase (ALP) from calf intestinal mucosa (Cat. No. P7640, Merck) was dissolved in buffer composed of 10 mM Tris-HCl (pH 8.0), 1.0 mM $MgCl_2$, 0.1 mM $ZnCl_2$, 50 mM KCl and 50% glycerol, and stored at –20°C. Exopolyphosphatase (PPX) from *E. coli* (HY-P71502, Hycultec GmbH, Beutelsbach, Germany) was dissolved in PBS (pH 7.4) (Cat. No. 9143.1, Carl Roth GmbH + Co. KG, Karlsruhe, Germany) supplemented with 6% trehalose and 0.1% BSA and stored at –20°C. The concentrations of ALP and PPX in stock solutions were 10 mg/μL and 100 μg/μL, respectively. For the ALP reaction, 2 μL of ALP (f.c. 50 μg/μL) stock solution was added to 398 μL of 1 mM $polyP_{130}$ in reaction buffer (50 mM Tris-HCl, pH 9.0; 1 mM $MgCl_2$; 0.2 mM $ZnCl_2$) and incubated at 37°C with shaking. For the PPX reaction, 2 μL of PPX (f.c. 0.5 μg/μL) stock solution was added to 398 μL of 1 mM $polyP_{130}$ in reaction buffer (50 mM Tris-HCl, pH 8.0; 150 mM KCl).

To monitor the reaction using SiX-DPA-Zn or DAPI, 4 μL of the reaction mixture was added to 396 μL of 20 μM SiX-DPA-Zn or 20 μM DAPI in 20 mM HEPES buffer (pH 7.5), and the fluorescence intensity was rapidly recorded at 678 nm for SiX-DPA-Zn (620 nm excitation) or 550 nm for DAPI (415 nm excitation) using a multimode microplate reader (Spark M20), with appropriate standard samples used for the calibration curves (0–20 μM). To measure inorganic phosphate released during the reaction, the reaction mixture was 50-fold diluted in 20 mM HEPES buffer (pH 7.5) and MG assay was performed according to the manufacturer's specifications (MAK307; Sigma-Aldrich). Measurements were taken at 0.5, 1, 1.5, 2, 3, and 6 h after the start of the reaction. The conversion rate was calculated as the ratio of the estimated polyP concentration in the presence of enzyme to that in the absence of enzyme at each time point (Equation S3). The conversion rate was also calculated directly from the phosphate concentration determined by the MG assay (Equation S4). $[\mathrm{polyP}]_0$represents the initial polyP concentration, which was fixed at 20 μM in this experiment.

$$\text{Conversion rate (\%)} = 100 \times \left(1 - \frac{[\text{polyP}]_{with\ enzyme}}{[\text{polyP}]_{without\ enzyme}}\right) \qquad \text{Equation S3}$$

$$\text{Conversion rate (\%)} = 100 \times \left(\frac{[\text{Pi}]_{measured}}{[\text{polyP}]_0}\right) \qquad \text{Equation S4}$$

**Maintenance of *Saccharomyces cerevisiae***

*Saccharomyces cerevisiae* (BY4741 wt and *vtc4*Δ) was cultured on yeast extract-peptone-dextrose (YPD) agar plates at 30 °C for about 3 days and were stored at 4 °C. A single colony was picked and precultured in 10 mL of YPD medium at 30 °C with shaking at 150 rpm for 18 h. Subsequently, 15 µL of the precultured medium was inoculated into 25 mL of fresh YPD medium and cultured under the same conditions (30 °C, 150 rpm) for 18 h. The $OD_{600}$ of cultured medium was measured by Biophotometer 6131 (Eppendorf, Hamburg, Germany). BY4741 *vtc4*Δ strain was from *Saccharomyces* Genome Deletion Project, obtained from Horizon discovery as a part of collection #YSC1053.

**PolyP quantification in *Saccharomyces cerevisiae* lysates**

Cultures ($OD_{600}$ ≈ 5, 25 mL) were transferred to 50 mL Falcon tubes and centrifuged at 1,000 × g for 2 min at 4 °C using a Sorvall RC 6 Plus (Thermo Fisher Scientific, MA, USA) equipped with an HS-4 rotor (Thermo Fisher Scientific). The supernatant was discarded, and the resulting cell pellet was resuspended in 2 mL of 20 mM HEPES buffer (pH 7.5) including 150 mM NaCl and 0.1% Triton X-100. Glass beads (425–600 µm diameter; Cat. No. G8772; Sigma-Aldrich) were added, and cells were lysed by vortexing for 5 min using cycles of 30 s vortexing followed by 30 s on ice (10 cycles total) using a vibra cell VCX 750 (Sonics, Newtown, CT, USA). After vortexing, the cell lysate was transferred to 2 mL tubes, heated to 95 °C for 5 min and centrifuged at 21,000 × g for 10 min at 4 °C in a Centrifuge 5424 R (Eppendorf) equipped with an FA-45-24-11 rotor (Eppendorf). The supernatant was collected and divided into two 2-mL tubes (1,000 µL each). The supernatant was used for quantitative polyP analysis after dilution (usually up to 2,000-fold) with 20 mM HEPES buffer (pH 7.5), adjusting the polyP concentration to approximately 10 µM, which falls within the linear range of the assay. The enzymatic polyP quantification was performed according to the manufacturer's instructions (Phosfinity Quant; Aminoverse, Nuth, Nederland). The polyP concentration was arbitrarily expressed as the value normalized to $OD_{600}$ (µM/$OD_{600}$), according to Equation S5. In this equation, $[\text{Pi}]$ denotes the polyP concentration in the cell lysate, $a$ denotes the dilution factor of the original cell culture (for example when 25 mL of cell culture is concentrated to 2 mL of cell lysate, $a$ is calculated as 2/25).

$$\text{Arbitrary polyP concentration} = a\frac{[\text{Pi}]}{\text{OD}_{600}}\left(\frac{\mu\text{M}}{\text{OD}_{600}}\right) \qquad \text{Equation S5}$$

**Phosphate depletion experiment in *Saccharomyces cerevisiae***

Cultures ($OD_{600}$ ≈ 5, 25 mL) were divided into two 15-mL Falcon tubes and centrifuged at 1,000 × g for 2 min at 4 °C using a Sorvall RC 6 Plus equipped with an HS-4 rotor. The cell pellets were washed with SC complete (-) phosphate medium (5.9 g/L yeast nitrogen base without amino acids and phosphate (Cat. No. CYN0801, Formedium, Norfolk, UK), 2.0 g/L Synthetic complete drop-out (Cat. No. DSCK012, Formedium), 2% glucose, and 10 mM KCl). One pellet was resuspended in an appropriate volume (25 or 50 mL) of SC complete (-) phosphate medium to $OD_{600}$ ≈ 1, whereas the other pellet was resuspended in SC complete (+) phosphate medium (5.9 g/L yeast nitrogen base without amino acids and phosphate, 2.0 g/L Synthetic complete drop-out, 2% glucose, and 10 mM $KH_2PO_4$) to $OD_{600}$ ≈ 1. The cultures were incubated at 30 °C with shaking 120 rpm. After 4 hours, $KH_2PO_4$ was added to the phosphate-depleted culture to a final concentration of 10 mM. Samples (5 mL) were collected every 2 hours (at 0, 2, 4, and 6 hour) to monitor $OD_{600}$ and polyP concentration. polyP concentrations were determined as described in the previous section.

**RNA/polyP extraction from Saccharomyces cerevisiae**

Yeast cultures ($OD_{600}$ ≈ 5, 25 mL) were transferred to 50 mL Falcon tubes and centrifuged at 1,000 × g for 2 min at 4 °C using a Sorvall RC 6 Plus equipped with an HS-4 rotor. The supernatant was discarded, and the resulting cell pellet was resuspended in 1 mL LETS buffer (0.1 M LiCl, 10 mM EDTA, 10 mM Tris-HCl, pH 8.0, 0.5% SDS) and 1 mL acid phenol (pH 4.5). Glass beads (425–600 µm diameter; Cat. No. G8772; Sigma-Aldrich) were added, and cells were lysed by vortexing for 5 min using cycles of 30 s vortexing followed by 30 s on ice (10 cycles total) using a vibra cell VCX 750 (Sonics, Newtown, CT, USA). The lysate was transferred to 2 mL tubes and centrifuged at 21,000 × g for 10 min at 4 °C using a Centrifuge 5424 R (Eppendorf) equipped with an FA-45-24-11 rotor (Eppendorf). The supernatant was collected and divided into two 2 mL tubes (500 µL each). RNA was precipitated by adding 1.5 mL ethanol to each tube, followed by incubation at –20°C for 5 h. Samples were then centrifuged at 9,400 × g for 10 min at 4 °C, and the supernatant was discarded. The resulting pellet was dried under reduced pressure and resuspended in buffer containing 10 mM Tris-HCl (pH 7.4), 0.1% SDS, and 1 mM EDTA. RNA concentration was determined based on $A_{260}$ measurements (1 $A_{260}$ unit = 40 µg/mL RNA). The RNA/polyP extracts from *S. cerevisiae* were further treated with 20 µg/mL RNase A (Cat. No. T3018L; New England BioLabs, MA, USA) for 1 hour at 37 °C to remove RNA in the polyP samples.

**PAGE of polyP and gel staining**

Polyacrylamide gels for electrophoresis were prepared using TBE buffer. The 10× TBE (pH 8.3) stock solution was prepared by dissolving 106 g Tris base, 55 g boric acid, and 4 mL of 500 mM EDTA (pH 8.0) in

1 L of deionized water. 15% polyacrylamide gels were polymerized by mixing 2 mL of 10× TBE stock solution and 7.5 mL of 40% acrylamide/bisacrylamide (19:1) (Cat. No. 3030.1; Carl Roth) with 10.5 mL of deionized water, followed by the addition of 20 µL TEMED and 200 µL freshly prepared 10% APS. The gels were cast at dimensions of 8.4 × 6.0 × 0.1 cm. Similarly, 30% gels were polymerized by mixing 8 mL of 10× TBE stock solution and 60 mL of 40% acrylamide/bisacrylamide (19:1) with 12 mL of deionized water, followed by 40 µL TEMED and 400 µL freshly prepared 10% APS, and cast at dimensions 13.4 × 12.8 × 0.1 cm. Before loading onto the gels, 10 µL of 6× loading solution (1 mM Orange G, 30 mM glycerol, 1 mM EDTA) was added to each 50 µL sample. 15% gels were run at 150 V for 45 min until the Orange G dye reached about 3 cm from the gels bottom. 30% gels were run at 100 V for 24 hours at 4 °C until Orange G dye reached about 7 cm from the gels bottom. For SiX-DPA-Zn staining, the gels were stained for 45 min in 10 µM SiX-DPA-Zn solution (10 mM HEPES buffer (pH 7.5), 20% ethanol, 2% glycerol) and then immediately gel images were captured under 630 nm excitation using Amersham Imager 600 (GE Healthcare Europe, GmbH, Freiburg, Germany). For DAPI staining, the gels were stained for 45 min in 10 µM DAPI solution (1.5 g/L Tris base, 2% glycerol) and then destained in 1.5 g/L Tris base with 2% glycerol for 30 min. Destained gels were irradiated with UV light (313 nm) and images were recorded after complete photobleaching of DAPI from the polyP bands, which typically required more than 10 min. UV irradiation and image acquisition was performed using Gel-Imager (Intas Science Imaging Instruments GmbH, Göttingen, Germany). Since prolonged UV exposure heats and dries the gels, they were cooled in water every 5 min during photobleaching. For JC-D7 staining, the gel was stained with 10 µM JC-D7 aqueous solution containing 20 % ethanol and 2% glycerol for 30 min. The image was captured under 460 nm excitation using Amersham Imager 600. For Toluidine Blue O (TB-O) staining, the gel was stained with 0.2% TB-O aqueous solution containing 20% ethanol and 2% glycerol for 60 min, and subsequently destained in 20% ethanol and 2% glycerol for 90 min. The image was captured under white light illumination.

**Quantification of polyP distribution using PAGE gel stained with SiX-DPA-Zn**

The $R_\mathrm{f}$ values of polyPs were determined following the Equation S6.

$$R_\mathrm{f} = \frac{polyP\ migration\ distance\ (\mathrm{cm})}{Orange\ G\ migration\ distance\ (\mathrm{cm})} \qquad \text{Equation S6}$$

Due to the large band distribution of polyPs, we estimated the polyP migration distance based on intensity-weighted centroid ($x_\mathrm{centroid}$) using the pixel position $x_i$ and signal intensities $I_i$ of each pixel within the band region (Equation S7).

$$x_\mathrm{centroid} = \frac{\sum x_i I_i}{\sum I_i} \qquad \text{Equation S7}$$

Empirically $R_f$ values were plotted versus the logarithm of polyphosphate chain lengths, followed by linear regression analysis (Equation S8)[9].

$$R_{\mathrm{f}} = a \log_{10} n + b \ (n \geq n^*) \qquad \text{Equation S8}$$

The fluorescence bands of the unknown polyP (hexametapolyphosphate (HP) (Cat. No. 305553; Sigma-Aldrich), and phosphate glass (Type 45) (Cat. No. S4379; Sigma-Aldrich)) were used with the calibration line to estimate the chain length distribution via the inverse function (Equation S9).

$$n = 10^{\frac{R_{\mathrm{f}}-b}{a}} \ (n \geq n^*) \qquad \text{Equation S9}$$

For the band regime below $n^*$, we adjusted the chain length distribution using free-solution model described in the next section.

The band peaks were determined based on their centroid positions ($x_{\mathrm{centroid}}$). However, since the fluorescence signal intensity ($I_i$) is expected to be proportional to the product of chain number ($n_i$) and number of molecules ($N_i$) (Equation S10), the $x_{\mathrm{centroid}}$ correspond to the weight-average chain length ($\bar{n}_{\mathrm{PAGE}} = \bar{n}_{\mathrm{w}}$) rather than the number-average chain length ($\bar{n}_{\mathrm{n}}$) which is calculated from $^{31}$P NMR spectra (Equation S11 and S12). Although many studies on polyphosphate gel analysis report results based on the number-average chain length, such analyses using similar calibration lines have not introduced significant errors[9-11]. Therefore, the present analysis also adopts this approach.

$$I_i \propto n_i N_i \qquad \text{Equation S10}$$

$$\bar{n}_{\mathrm{PAGE}} = \bar{n}_{\mathrm{w}} = \frac{\sum n_i^2 N_i}{\sum n_i N_i} \propto \frac{\sum n_i I_i}{\sum I_i} \qquad \text{Equation S11}$$

$$\bar{n}_{\mathrm{NMR}} = \bar{n}_{\mathrm{n}} = \frac{\sum n_i N_i}{\sum N_i} \propto \frac{\sum I_i}{\sum \frac{I_i}{n_i}} \qquad \text{Equation S12}$$

**Free-solution model vs Ogston sieving model**

The migration behavior of polyPs in gel electrophoresis strongly depends on the relationship between the polymer's radius of gyration ($R_{\mathrm{g}}$) and the characteristic mesh size of the gel ($\xi$). Two distinct regimes can be identified depending on whether the polymer is significantly smaller or comparable to the pore size of the gel matrix.

**Free-solution model (short chains ($R_{\mathrm{g}} \ll \xi$)):** When the polymer's radius of gyration $R_{\mathrm{g}}$ are much smaller than the characteristic pore size ($R_{\mathrm{g}} \ll \xi$), the mobility of polyP is estimated as a free-solution model without considering the gel sparse network. In this regime, the mobility ($\mu$) can be expressed as a function of the chain length number ($n$):

$$\mu = \frac{q}{f} \propto \frac{1}{R_{\mathrm{g}}} \propto n^{-\nu} \qquad \text{Equation S13}$$

where $q$ is the effective charge, $f$ is the friction coefficient, and $\nu$ is the scaling exponent which typically takes a value of around 0.4–0.7 for a flexible coil, including biological polymers such as unfolded proteins[12] and single-stranded DNA[13]. The $\log R_{\mathrm{f}}$ ($R_{\mathrm{f}} \propto \mu$) versus $\log n$ gives linear regression with the slope $\nu$:

$$\log R_{\mathrm{f}} = a - \nu \log n \qquad \text{Equation S14}$$

In this study, the linear regression was performed for the range corresponding to polyP$_{3-7}$. The regression yields $\nu = 0.4122$, which is a reasonable value when compared with previous studies on other biological polymers.

**Ogston sieving model (long chains ($R_{\mathrm{g}} \sim \xi$)):** When the polymer's size becomes comparable to or larger than the mesh size ($R_{\mathrm{g}} \sim \xi$), the molecule experiences significant sieving effects due to steric hindrance by the gel network. In this case, migration requires the polymer to deform and thread through the gel pores, and the mobility shows a strong dependence on the polymer size (and thus on its molecular weight). Empirically, this behavior is commonly described by Ferguson equation which was originally derived from the Ogston model (Equation S15)[14,15].

$$\log \mu = \log \mu_0 - K_{\mathrm{r}} C_{\mathrm{g}} \qquad \text{Equation S15}$$

where $\mu_0$ is the mobility in the free-solution, and $C_{\mathrm{g}}$ is the gel concentration determined by the gel conditions. $K_{\mathrm{r}}$ (the retardation coefficient) depends on the molecular size. For polymers, $K_{\mathrm{r}}$ scales with the chain length $n$ ($K_{\mathrm{r}} \propto n^{\nu}$). Consequently, the Ferguson model effectively captures the size-dependent sieving regime, in which empirically the mobility ($\mu$) decreases with increasing $\log n$ as described in the previous section[9,10,16,17] (Equation S8).

In this study, we applied the free-solution model to polyP$_{3–7}$ and the Ogston sieving model to polyP$_{14–130}$. The transition zone (polyP$_{7–14}$) was represented by a line connecting the $R_{\mathrm{f}}$ values of polyP$_7$ and polyP$_{14}$ plotted against log *n*, to complete the calibration curve shown in Figure 4D. The chain-length distributions of HP, PG, and polyp extract from *S. cerevisiae* were then determined using these calibration curves, as shown in Figure 4E and 4F.

**Measurement of number-average chain length of HP and PG by $^{31}$P NMR**

PolyP samples (HP and PG) for $^{31}$P NMR measurements were prepared by dissolving in $D_2O/H_2O$ (8:2), and the final concentration was adjusted to 10 mg/mL. $^{31}$P NMR (162 MHz) spectra were recorded on an Agilent 400–MR spectrometer. The number-average chain length ($\bar{n}_{\mathrm{NMR}}$) of samples was determined by $^{31}$P NMR spectra following the reference[3,18]. Briefly, it was calculated from the ratio of the integral of the terminal phosphate signal ( $\mathrm{PP1}$ ) to that of the penultimate and core phosphate signals ($\mathrm{PP2}, \mathrm{PP3}, \text{and } \mathrm{PP4}$) (Equation S16).

$$\bar{n}_{\mathrm{NMR}} = 2\frac{\sum \mathrm{PP}i}{\mathrm{PP1}}\ (i = 1, 2, 3, \text{and } 4) \qquad \text{Equation S16}$$

**Maintenance and preparation of the PPK1-TRex cells**

PPK1-TREx cells[4] (a kind gift from dr. A. Saiardi) were cultured in high-glucose DMEM supplemented with 10% FBS in a humidified incubator containing 5% $CO_2$ at 37 °C. The cells were passaged every 3 – 4 days upon reaching full confluence. For confocal imaging, cells were seeded onto 8-well high glass-bottom plate (Cat. No. 80807; ibidi GmbH, Gräfelfing, Germany) and cultured for 2 days in a humidified incubator. Doxycycline (1µM) was added 24 h prior to imaging to induce PPK1 expression via the Tet-regulated expression system. For the FACS study, cells were seeded onto 6-well plates (Cat. No. 83.3920; Sarstedt AG & Co. KG, Nümbrecht, Germany) and prepared in the same manner as described above.

**Fixed cell staining**

Cells were washed twice with PBS (Cat. No. 9143.1; Carl Roth) and fixed in 4% paraformaldehyde (prepared from 16% Formaldehyde solution; Cat. No. 28908, Thermo Fisher Scientific) in PBS for 10 min. After fixation, cells were incubated in 30 mM glycine in PBS for 5 min and washed twice with PBS. Cells were then permeabilized with 0.1% Triton X-100 in PBS for 10 min, followed by two PBS washes and blocking in 3% BSA (Cat. No. A7030; Merck) in PBS for 60 min. After two additional PBS washes, cells were stained with 5.0 µM SiX-DPA-Zn and 1.0 µM Hoechst 33342 in PBS for 20 min. For DAPI staining, cells were incubated with 1.0 µM DAPI in PBS. ALP treatment was carried out prior to blocking by incubating the cells in 1.0 mg/mL ALP prepared in 50 mM Tris-HCl buffer (pH 9.3) containing 1 mM $MgCl_2$, 0.1 mM $ZnCl_2$, and 1 mM spermidine for 1 hour, followed by two washes with PBS.

**Immunostaining of fibrillarin and nucleolin**

After fixation, permeabilization, and blocking as described above, cells were incubated for 1 h at room temperature with either an anti-fibrillarin mouse monoclonal antibody (Cat. No. Ab4566, Abcam, Cambridge, UK) diluted 1:100, or an anti-nucleolin rabbit polyclonal antibody (Cat. No. ab22758, Abcam) diluted 1:250 in PBS containing 1% BSA. Cells were then washed three times with PBS containing 1% BSA (10 min each). Subsequently, cells were incubated for 1 h at room temperature with either a donkey anti-mouse IgG (H+L), highly cross-adsorbed, secondary antibody conjugated to Alexa Fluor Plus 488 (Cat. No. A21202, Invitrogen, MS, USA) for fibrillarin, or a donkey anti-rabbit IgG (H+L), highly cross-adsorbed, secondary antibody conjugated to Alexa Fluor Plus 488 (Cat. No. A32790, Invitrogen, MS, USA) for nucleolin, diluted 1:1000 in PBS containing 1% BSA. For multi-color STED imaging, we used an sdAb anti-mouse IgG1 Halo fusion (Cat. No. N2041, NanoTag Biotechnologies GmbH, Göttingen, Germany) for fibrillarin or an sdAb anti-rabbit IgG1 Halo fusion (Cat. No. N2441, NanoTag Biotechnologies GmbH) for nucleolin. These HaloTag fusion nanobodies were labeled by preincubating with 6-580CP-Halo ligand for 20 min at room temperature to a final concentration of 1 µM. The mixture (5 µL) was then directly added to each well containing 230 µL of PBS (1% BSA) and incubated for 1 h at room temperature. After incubation, cells were washed three times with PBS containing 1% BSA (10 min each). Finally, cells were stained with the appropriate dyes, including SiX-DPA-Zn and Hoechst 33342.

**Confocal imaging**

Confocal imaging was performed using Visitron Spinning disk/TIRF/SMLM system (Visitron Systems GmbH, Germany) equipped with 50 µm pinhole disk. Images were acquired with a Visitron imaging system (VisiView software, version 6.0.0.16) based on Nikon optics (Nikon, Tokyo, Japan). The Ti2-W1 was equipped with a CFI Plan Apo Lambda 60× oil-immersion objective lens (NA 1.4; Cat. No. MRD73250, Nikon). Excitation/emission settings were as follows: 640 nm laser / ET665lp for SiX-DPA-Zn detection; 405 nm laser / ET460/50 (435–485 nm) for Hoechst 33342 and DAPI–DNA detection; and 405 nm laser / ET525/50 (500–550 nm) for DAPI–polyP detection. Z-stack images were collected from –5.00 to +5.00 µm at 0.20 µm intervals (51 steps).

**Flow cytometry**

Cells were washed twice with PBS, detached from the culture dish with 600 µL StemRro™Accutase™ (Cat. No. A1110501; Thermo Fisher Scientific), and collected in 2 mL Eppendorf tubes. Pellets were sequentially centrifuged at 94 × g at 4 °C (2 min before fixation, 5 min after fixation), with the supernatant removed and pellets resuspended in PBS as needed before each step. Cells were fixed with 1 mL of 4% paraformaldehyde (first, resuspended in 500 µL PBS, then 500 µL of 8% paraformaldehyde added) for 10 min, quenched with 1 mL of 30 mM glycine in PBS for 5 min, permeabilized with 1 mL of 0.1% Triton X-100 in PBS for 10 min, and blocked with 1 mL of 3% BSA (Cat. No. A7030; Merck) in PBS for 60 min. Finally, the pellets were stained with 5.0 µM SiX-DPA-Zn in PBS for 20 min and washed with PBS prior to further analysis.

Flow cytometry was performed using FACS ARIA™ III Sorter (BD Biosciences, NJ, USA). For each experiment, 50,000 or 100,000 cells were analyzed. The fluorescence signal from SiX-DPA-Zn was recorded using a 670/14 nm filter, a 630 LP mirror and 633 nm excitation laser. The fluorescence signal from DAPI was recorded using a 510/50 nm emission filter, a 502 LP mirror and 405 nm excitation laser. The collected data were processed using software FlowJo (BD Biosciences).

**STED imaging and 3D confocal imaging**

STED imaging was performed using an Abberior Mirava STED microscope (Abberior Instruments GmbH, Göttingen, Germany) equipped with a UPLSAPO 60×/NA 1.4 oil-immersion objective lens (Evident Corporation, Tokyo, Japan, formerly Olympus) and a pulsed 775 nm 40 MHz 3 W STED laser or Abberior Expert STED microscope equipped with UPlanSApo 100×/NA 1.4 oil-immersion objective lens (Evident Corporation) and a pulsed 775 nm 40 MHz 3W STED laser. Microscope has two APD and two MATRIX detectors which can be tuned to any detection window in the range 400 – 800 nm. The pixel size was set to 20 (for Abberior Mirava STED microscope) or 30 nm (for Abberior Expert STED microscope) in the X-Y plane for STED imaging and 60 nm in the X-Y plane for 3D confocal imaging. Z-stack images were acquired with a step size of 0.20 µm (Abberior Mirava STED microscope). Excitation/emission settings were as follows: 640 nm/680–705 nm for SiX-DPA-Zn; 561 nm/605–625 for 580CP-Halo detection.

**Data processing**

Data analyses, including both linear and non-linear regression (Hill plot analysis and correlation), were performed using GraphPad Prism 10.4 (GraphPad Software, Inc., San Diego, CA, USA). Statistical comparisons between groups were performed using unpaired two-tailed t-tests in Microsoft Excel (Microsoft Corporation, Redmond, WA, USA).

**Image processing and analysis (confocal images)**

All acquired images were processed using ImageJ (Fiji[19]) and CellProfiler (version 4.2.8, Broad Institute). Image contrast was adjusted as needed, and multicolor images were generated using Fiji (ImageJ). Z-stack images were first converted into single-slice maximum-intensity projection images using Fiji, and the resulting images were segmented using CellProfiler. Nuclear regions were identified from Hoechst 33342 or DAPI-stained DNA images using the Otsu thresholding method (typical object diameter: 50–150 pixels; threshold smoothing scale: 1; threshold correction factor: 0.9; lower and upper bounds on threshold:

0.002–1.0). For quantification of DAPI–polyP signals, the signals were first enhanced and segmented (feature type: speckles; feature size: 10). PolyP areas were then extracted using the Otsu thresholding method (typical object diameter: 1–50 pixels; threshold smoothing scale: 1.35; threshold correction factor: 1.0; lower and upper bounds on threshold: 0.0–1.0). The mean signal intensity of polyP labeled with SiX-DPA-Zn or DAPI was measured for each nucleus.

For colocalization analysis the integrated pixel intensity of fibrillarin, nucleolin, and polyp (Figure 5E and 5G), representing the total signal intensity divided by each area of the nucleus, was measured using Fiji. Nuclear regions were defined based on Hoechst 33342 staining by applying a thresholding method, followed by particle analysis with the following parameters: size = 50–infinity and circularity = 0.10–1.00.

**Image processing (3D images)**

Three-dimensional (3D) rendered images and rotation movies were generated using LIGHTBOX software (Lightbox 2025.10.22339-g33d8d658c0, Abberior Instruments GmbH). X–Y images were obtained by maximum intensity projection in Fiji. X–Z and Y–Z views were generated using the orthogonal view function in Fiji.

**Image processing and analysis (STED images)**

Signal intensity profile was obtained using the plot profile function in Fiji. The full width at half maximum (FWHM) values were determined by Lorentzian fitting using GraphPad Prism 10.4, according to Equation S17. In the fitting equation, $I(x)$, $A_i$, $x_i$, and $w_i$ represent signal intensity at position $x$, amplitude, the peak position, width (corresponding to the FWHM), respectively. The number of peaks ($n$) was set to 2 for STED and 1 or 2 for confocal profile.

$$I(x) = \sum_{i=1}^{n} \frac{A_i}{\left(\frac{x-x_i}{w_i}\right)^2+1} \qquad \text{Equation S17}$$

**Synthetic procedures**

**Compound 1**

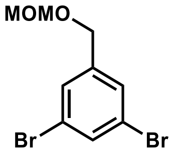

**1,3-dibromo-5-((methoxymethoxy)methyl)benzene**
**Chemical Formula:** $C_9H_{10}Br_2O_2$
**Exact Mass: 307.9048**

(3,5-Dibromophenyl)methanol (14.0 g, 52.7 mmol, 1.0 eq.) and *N*,*N*-diisopropylethylamine (11.0 mL, 63.3 mmol, 1.2 eq.) were dissolved in 100 mL dichloromethane (DCM) and cooled to 0 °C. Chloromethyl methyl ether (MOM-Cl) (4.79 mL, 63.3 mmol, 1.2 eq.) was then added dropwise to the solution at 0 °C with stirring. After the addition of MOM-Cl, the reaction mixture was allowed to warm to room temperature and stirred for 16 hours. The reaction was quenched with saturated aqueous $NaHCO_3$. and the organic layer was extracted with ethyl acetate three times. The combined organic layer was further washed with saturated aqueous $NaHCO_3$. and brine and dried over anhydrous $Na_2SO_4$. The solvent was removed under reduced pressure and the product was completely dried in vacuo. The compound 1 (15.1 g, 48.8 mmol, 93%) was obtained as colorless oil and used for next reaction without further purification.

**$^1$H NMR (400 MHz, $CDCl_3$):** $\delta$ (ppm) = 7.60 (m, 1H), 7.45 (m, 2H), 4.71 (s, 2H), 4.54 (m, 2H), 3.41 (s, 3H)
**$^{13}$C NMR (101 MHz, $CDCl_3$):** $\delta$ (ppm) = 142.00, 133.16, 129.20, 122.93, 95.92, 67.55, 55.55
**MS (ESI+)** For unknown reasons, the signal could not be detected.

**Compound 2**

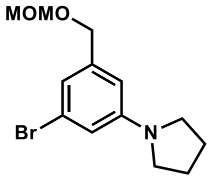

**1-(3-bromo-5-((methoxymethoxy)methyl)phenyl)pyrrolidine**
**Chemical Formula:** $C_{13}H_{18}BrNO_2$
**Exact Mass: 299.0521**

Compound 1 (7.57 g, 24.4 mmol, 1.0 eq.), pyrrolidine (2.0 mL, 24.4 mmol, 1.0 eq.), $Pd_2dba_3$ (560 mg, 610 µmol, 0.025 eq.), rac-BINAP (1.14 g, 183 µmol, 0.075 eq.), and Sodium *tert*-butoxide (2.8 g, 29.3 mmol, 1.2 eq.) were dissolved in anhydrous toluene (30 mL) in an argon-purged sealed tube. The reaction mixture was stirred at 80 °C for 4 hours and then cooled down to room temperature. The mixture was filtered through Celite to remove palladium catalyst, followed by washing with methanol. The filtrate was evaporated and the crude product was purified by column chromatography (ethyl acetate/hexane = 0–50%) to afford compound 2 (4.53 g, 15.1 mmol, 62%) as pale-yellow oil.

**$^1$H NMR (400 MHz, CDCl$_3$):** δ (ppm) = 6.79 (m, 1H), 6.62 (m, 1H), 6.46 (m, 1H), 4.70 (s, 2H), 4.51 (m, 2H), 3.42 (s, 3H), 3.27(m, 4H), 2.01 (m, 4H)

**$^{13}$C NMR (101 MHz, CDCl$_3$):** δ (ppm) = 148.91, 140.43, 123.32, 117.49, 113.65, 109.62, 95.63, 68.85, 55.41, 47.71, 25.41

**HRMS (ESI+)** $[M(^{79}Br)+Na]^+$ found 322.0414, calculated 322.0413, $[M(^{81}Br)+Na]^+$ found 324.0394, calculated 324.0393.

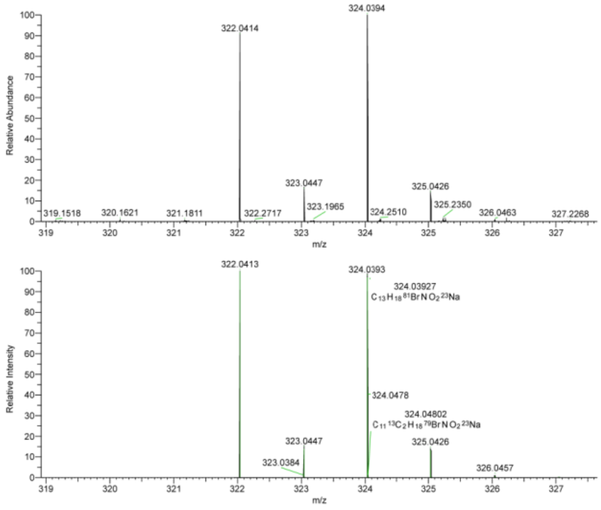

HRMS (ESI) spectra of Compound 2. Top: observed spectrum; Bottom: calculated spectrum.

#### Compound 3

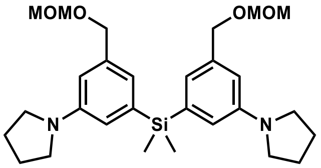

**bis(3-((methoxymethoxy)methyl)-5-(pyrrolidin-1-yl)phenyl)dimethylsilane**
**Chemical Formula:** $C_{28}H_{42}N_2O_4Si$
**Exact Mass: 498.2914**

Compound 2 (2.50 g, 8.33 mmol, 1.0 eq.) was dissolved in 40 mL of anhydrous THF under an argon atmosphere at –78 °C. A solution of 2.5 M *n*-butyllithium (3.3 mL, 8.33 mmol, 1.0 eq.) was added dropwise over 5 min to the solution at –78 °C and the mixture was stirred for an additional 40 min at the same temperature. A solution of Dichlorodimethylsilane (452 µL, 3.75 mmol, 0.45 eq.) in anhydrous THF (10 mL) was added to the reaction mixture at –78 °C and the mixture was subsequently stirred while warming to room temperature for 3 h. Upon completion of the reaction, as confirmed by TLC, the mixture was quenched with saturated aqueous $NaHCO_3$ and extracted with ethyl acetate. The combined organic layers were washed with brine, dried over $Na_2SO_4$, filtered, and concentrated under reduced pressure. The crude product was purified by column chromatography (ethyl acetate/hexane = 0–30%) to afford compound 3 (1.17 g, 2.35 mmol, 63%) as colorless oil.

**$^1$H NMR (400 MHz, $CDCl_3$):** δ (ppm) = 6.85 (s, 2H), 6.72 (s, 2H), 6.61 (s, 2H), 4.72 (s, 4H), 4.57 (s, 4H), 3.43 (s, 6H), 3.30 (m, 8H), 2.00 (m, 8H), 0.54 (s, 6H)

**$^{13}$C NMR (101 MHz, $CDCl_3$):** δ (ppm) = 147.34, 139.23, 137.96, 121.05, 116.96, 112.17, 95.58, 69.73, 55.31, 47.76, 25.40, –2.06

**HRMS (ESI+)** $[M+H]^+$ found 499.2983, calculated 499.2987

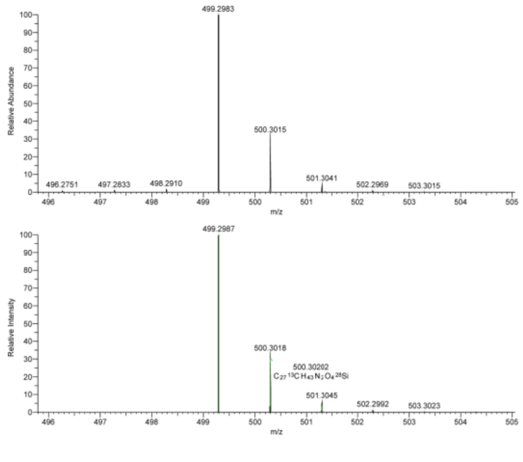

HRMS (ESI) spectra of Compound 3. Top: observed spectrum; Bottom: calculated spectrum.

**Compound 4**

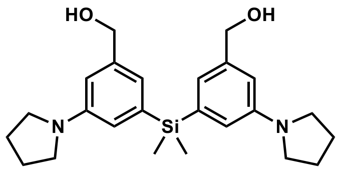

**((dimethylsilanediyl)bis(5-(pyrrolidin-1-yl)-3,1-phenylene))dimethanol**
**Chemical Formula:** $C_{24}H_{34}N_2O_2Si$
**Exact Mass: 410.2390**

20 mL of 6M hydrochloric acid was added to a solution of compound 3 (1.17 g, 2.35 mmol) in methanol (20 mL) and stirred at 50 °C for 30 min. The reaction mixture was cooled to room temperature and then quenched with saturated aqueous $NaHCO_3$. The product was extracted with ethyl acetate three times, washed with brine, dried over by anhydrous $Na_2SO_4$. After removal of the solvent under reduced pressure, the residue was thoroughly dried in vacuum. The compound 4 (1.00 g, 2.02 mmol) was used for next reaction without further purification.

**$^1$H NMR (400 MHz, CDCl$_3$):** $\delta$ (ppm) = 6.87 (s, 2H), 6.76 (s, 2H), 6.66 (s, 2H), 4.63 (s, 4H), 3.31 (m, 8H), 2.01 (m, 8H), 0.54 (s, 6H)

**$^{13}$C NMR (101 MHz, CDCl$_3$):** $\delta$ (ppm) = 147.18, 141.34, 139.56, 120.47, 117.07, 111.60, 65.94, 48.26, 25.37, –2.15

**HRMS (ESI+)** $[M+Na]^+$ found 433.2280, calculated 433.2282

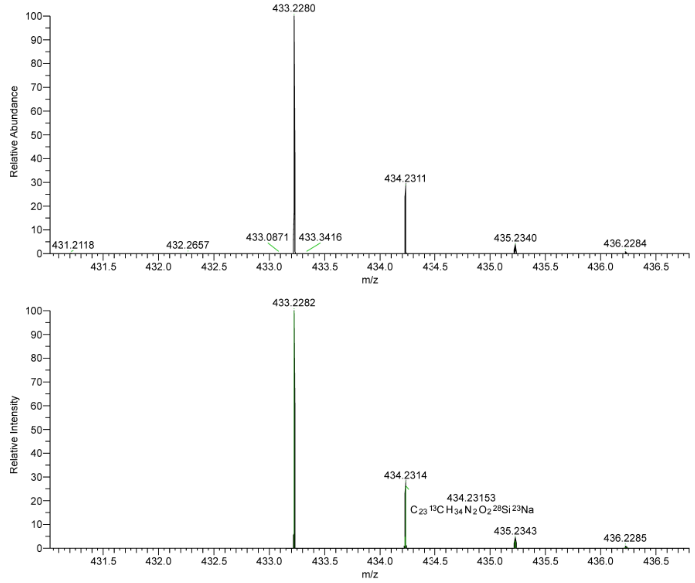

HRMS (ESI) spectra of Compound 4. Top: observed spectrum; Bottom: calculated spectrum.

**Compound 5**

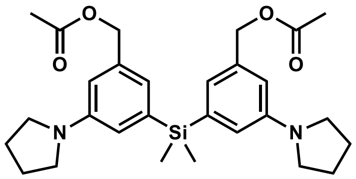

**((dimethylsilanediyl)bis(5-(pyrrolidin-1-yl)-3,1-phenylene))bis(methylene) diacetate**
**Chemical Formula:** $C_{28}H_{38}N_2O_4Si$
**Exact Mass: 494.2601**

The crude compound 4 (1.00 g, 2.02 mmol, 1.0 eq.) was dissolved in DCM (20 mL). A solution of triethylamine (1.2 mL, 8.10 mmol, 4.0 eq.), acetic anhydride (567 µL, 6.00 mmol, 3.0 eq.), and DMAP (73 mg, 598 µmol, 0.3 eq.) in DCM (10 mL) was then added to this solution and stirred at room temperature for 2 h. Upon completion of the reaction, as confirmed by TLC, the solvent was removed under reduced pressure. The concentrate was purified by column chromatography (ethyl acetate/hexane = 0–20%) to afford compound 5 (775 mg, 1.57 mmol, 67% (from compound 3)) as colorless oil.

**$^1$H NMR (400 MHz, CDCl$_3$):** δ (ppm) = 6.83 (s, 2H), 6.73 (s, 2H), 6.58 (s, 2H), 5.05 (s, 4H), 3.29 (m, 8H), 2.09 (s, 6H), 2.00 (m, 8H), 0.53 (s, 6H)

**$^{13}$C NMR (101 MHz, CDCl$_3$):** δ (ppm) = 170.98, 147.28, 139.41, 136.02, 121.34, 117.48, 112.52, 67.09, 47.69, 25.41, 21.10, –2.14

**HRMS (ESI+)** [M+H]$^+$ found 495.2671, calculated 495.2674

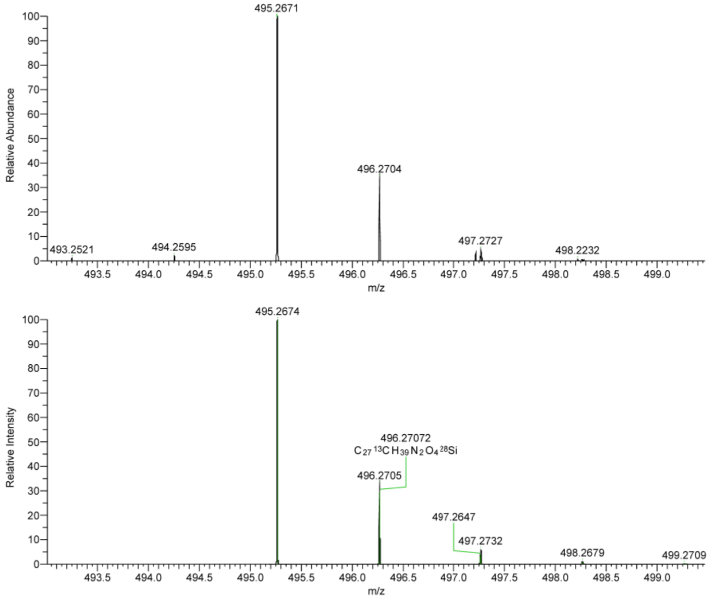

HRMS (ESI) spectra of Compound 5. Top: observed spectrum; Bottom: calculated spectrum.

**Compound 6**

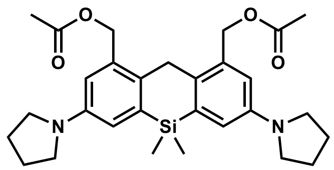

**(5,5-dimethyl-3,7-di(pyrrolidin-1-yl)-5,10-dihydrodibenzo[*b*,*e*]siline-1,9-diyl)bis(methylene) diacetate**
**Chemical Formula:** $C_{29}H_{38}N_2O_4Si$
**Exact Mass: 506.2601**

An aqueous solution of 37% paraformaldehyde (520 µL, 7.00 mmol, 5.0 eq.) was added to a solution of compound 5 (700 mg, 1.38 mmol, 1.0 eq.) in acetic acid (10 mL), and the mixture was gradually warmed to 55 °C with stirring. Upon warming to 55 °C, the solution gradually turned deep blue, and the mixture was subsequently stirred at the same temperature for 1 h. Completion of the reaction was confirmed by TLC monitoring (Rf = 0.5, ethyl acetate/hexane = 33%). The mixture was then cooled to room temperature and quenched with saturated aqueous $NaHCO_3$. The organic layer was extracted with chloroform three times, washed with brine, and dried over anhydrous $Na_2SO_4$. The combined organic layers were concentrated under reduced pressure and dried thoroughly in vacuo. Compound 6 was used for next reaction without further purification.

**$^1$H NMR (400 MHz, $CDCl_3$):** $\delta$ (ppm) = 6.79 (s, 2H), 6.60 (s, 2H), 5.27 (s, 4H), 4.02 (s, 2H), 3.32 (m, 8H), 2.12 (s, 6H), 2.02 (m, 8H), 0.48 (s, 6H)

**$^{13}$C NMR (400 MHz, $CDCl_3$):** The obtained compound 6 was found to be unstable and readily underwent oxidation, which prevented acquisition of a clear $^{13}$C NMR spectrum.

**HRMS (ESI+)** $[M+Na]^+$ found 529.2493, calculated 529.2493

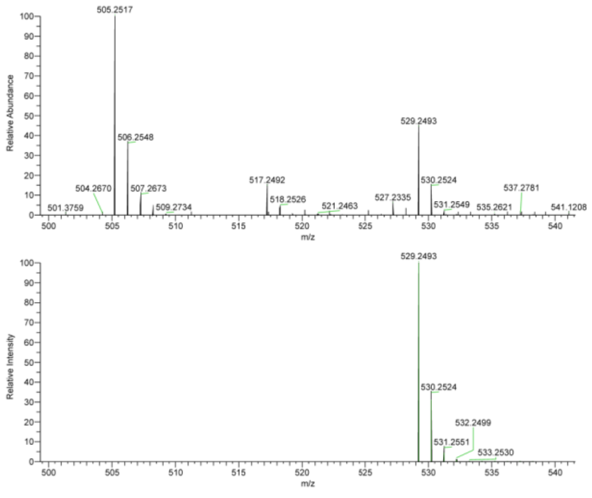

HRMS (ESI) spectra of Compound 6. Top: observed spectrum; Bottom: calculated spectrum.

**Compound 7**

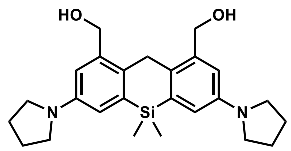

**(5,5-dimethyl-3,7-di(pyrrolidin-1-yl)-5,10-dihydrodibenzo[*b*,*e*]siline-1,9-diyl)dimethanol**
**Chemical Formula:** $C_{25}H_{34}N_2O_2Si$
**Exact Mass: 422.2390**

2 M $LiBH_4$ in THF (1.4 mL, 2.80 mmol, 2.0 eq.) was added to the solution of compound 6 (708 mg, 1.40 mmol, 1.0 eq.) in anhydrous THF at 0 °C and subsequently stirred while warming to room temperature for 0.5 h. Afterwards, the reaction mixture was cooled to 0 °C again, the mixture of 1 M aqueous NaOH and methanol was added to this solution. The reaction mixture was stirred for an additional 20 min, quenched with saturated aqueous $NaHCO_3$, and extracted three times with ethyl acetate. The combined organic extracts were washed with brine, dried over anhydrous $Na_2SO_4$, and concentrated under reduced pressure. The crude product was purified by column chromatography (ethyl acetate/hexane = 12–100%) to yield compound 7 (295 mg, 700 µmol, 50%) as a colorless to light greenish-blue powder. The product is prone to oxidation, turning deep blue upon exposure to air, and was therefore stored under an inert atmosphere at –20 °C until use for next reaction.

**$^1$H NMR (400 MHz, $CDCl_3$):** $\delta$ (ppm) = 6.82 (s, 2H), 6.59 (s, 2H), 4.77 (s, 4H), 4.08 (s, 2H), 3.31 (s (br), 8H), 2.00 (s(br), 8H), 0.48 (s, 6H)

**$^{13}$C NMR (101 MHz, $CDCl_3$):** $\delta$ (ppm) = 145.56, 138.53, 137.64, 132.90, 116.50, 114.56, 64.78, 48.40, 28.87, 25.34, –3.14

**HRMS (ESI+)** $[M+Na]^+$ found 445.2281, calculated 445.2282

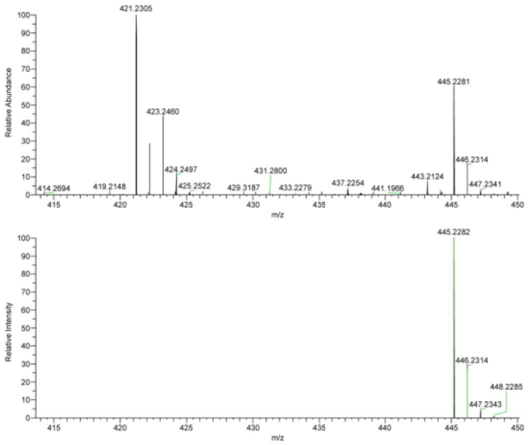

HRMS (ESI) spectra of compound 7. Top: observed spectrum; Bottom: calculated spectrum.

#### Compound 8

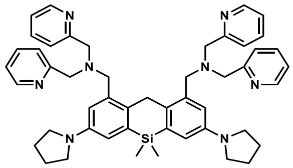

1,1'-(5,5-dimethyl-3,7-di(pyrrolidin-1-yl)-5,10-dihydrodibenzo[*b*,*e*]siline-1,9-diyl)bis(*N*,*N*-bis(pyridin-2-ylmethyl)methanamine)
Chemical Formula: $C_{49}H_{56}N_8Si$
Exact Mass: 784.4397

The solution of triphenylphosphine (463 mg, 1.77 mmol, 3.0 eq.) in DCM (5 mL) was added dropwise to the solution of compound 7 (250 mg, 591 µmol, 1.0 eq.) and tetrabromoethane (471 mg, 1.42 mmol, 2.4 eq.) in DCM (10 mL) under argon atmosphere at 0 °C and allowed to warm to room temperature for 2 hours. The reaction mixture was concentrated under reduced pressure and subsequently dried in vacuo for 1 hour. The crude dibrominated product was dissolved in anhydrous DMF with bis(2-pyridylmethyl)amine (236 mg, 1.18 mmol, 2.0 eq. ) and $K_2CO_3$ (327 mg, 2.36 mmol, 4.0 eq.) and stirred for 3 hours at room temperature. The reaction mixture was quenched with water and organic layer was extracted with ethyl acetate three times, washed with brine, and dried over anhydrous $Na_2SO_4$. After removal of solvent under the reduced pressure, the residue was purified by column chromatography (methanol/DCM with 1% TEA = 0–10%) to yield compound 8 (141 mg, 179 µmol, 30%) as a brawn sticky oil.

**$^{1}$H NMR (400 MHz, $CDCl_3$):** δ (ppm) = 8.49 (d, *J* = 4.8 Hz, 4H), 7.55 (m, 8H), 7.09 (d, *J* = 4.8 Hz, 4H), 6.94 (d, *J* = 2.6 Hz, 2H), 6.67 (d, *J* = 2.6 Hz, 2H), 4.11 (s, 4H), 3.29 (m, 8H), 1.99 (m, 8H), 0.42 (s, 6H)

**$^{13}$C NMR (101 MHz, $CDCl_3$):** δ (ppm) = 159.67, 148.87, 145.57, 136.53, 136.33, 136.27, 131.91, 122.49, 121.75, 114.51, 114.31, 60.00, 57.46, 47.75, 29.03, 25.43, –2.13

**HRMS (ESI+)** $[M+H]^+$ found 785.4469, calculated 785.4470

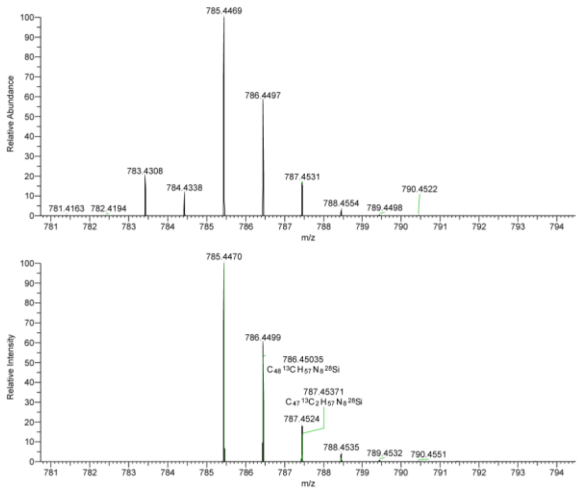

HRMS (ESI) spectra of Compound 8. Top: observed spectrum; Bottom: calculated spectrum.

**SiX-DPA**

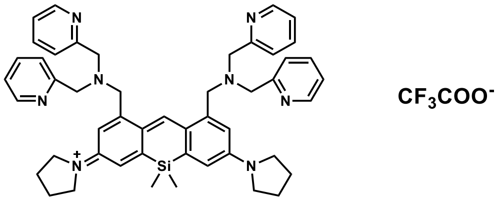

1-(1,9-bis((bis(pyridin-2-ylmethyl)amino)methyl)-5,5-dimethyl-7-(pyrrolidin-1-yl)dibenzo[*b*,*e*]silin-3(5*H*)-ylidene)pyrrolidin-1-ium
Chemical Formula: $C_{49}H_{55}N_8Si^+$
Exact Mass: 783.4313

Chloranil (60 mg, 244 µmol, 1.7 eq.) was added to the solution of compound 8 (112 mg, 143 µmol, 1.0 eq.) in anhydrous DCM (10 mL). The solution was stirred at room temperature for 30 min. The reaction mixture was evaporated under reduced pressure to remove the solvent, while keeping the temperature below 20 °C. The crude product was purified by reverse-phase column chromatography using a linear gradient of $CH_3CN$/water containing 0.1% trifluoroacetic acid (30:70 → 100:0). The collected fractions were combined and lyophilized to afford SiX-DPA as a purple powder (41 mg, 52.4 µmol, 37%). The purity of SiX-DPA was confirmed by LC–MS spectroscopy.

**$^1$H NMR (400 MHz, $CDCl_3$):** δ (ppm) = 8.83 (m, 4H), 8.52 (s, 1H), 8.32–8.21 (m, 8H), 7.69 (m, 4H), 7.07 (d, 2H), 6.68 (d, 2H), 4.51 (s, 8H), 4.34 (s, 4H), 3.67 (s(br), 8H), 2.13 (m, 8H), 0.35 (s, 6H)

**$^{13}$C NMR (101 MHz, $CDCl_3$):** δ (ppm) = 160.86, 153.58, 152.00, 148.47, 147.82, 144.91, 142.62, 127.91, 125.35, 120.41, 117.58, 116.12, 56.43, 56.01, 49.22, 25.00, –1.05

**HRMS (ESI+)** $[M+H]^+$ found 783.4308, calculated 783.4313

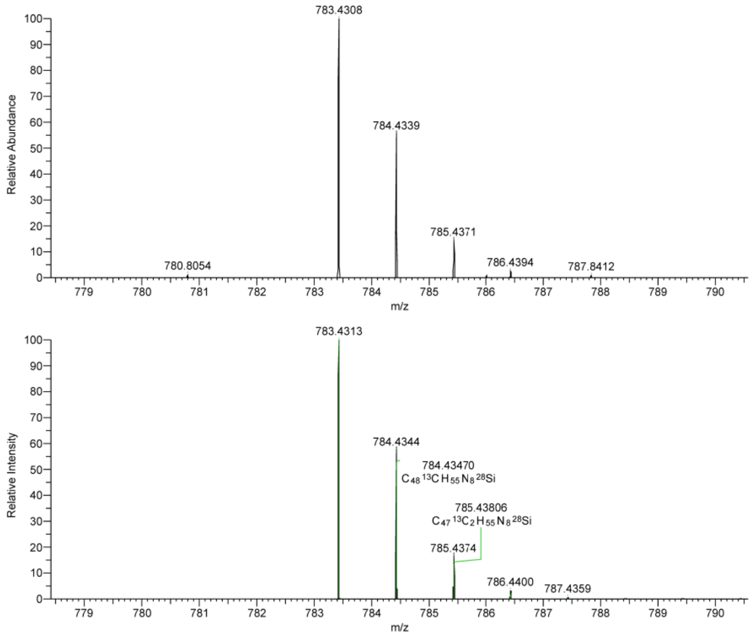

HRMS (ESI) spectra of SiX-DPA. Top: observed spectrum; Bottom: calculated spectrum.

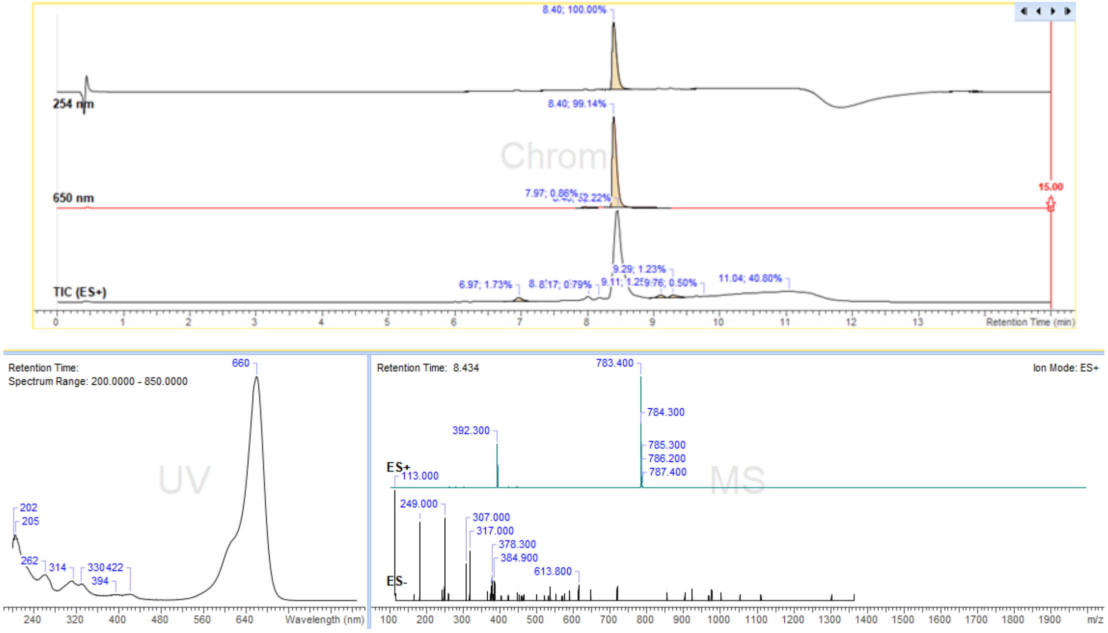

LC–MS spectra of SiX-DPA (after purification). The data are the same as those shown in Figure S1.

#### NMR stectra

Solvent peaks and trace impurities (if observed) are characterized using ref8.

**$^{1}$H NMR of compound 1**

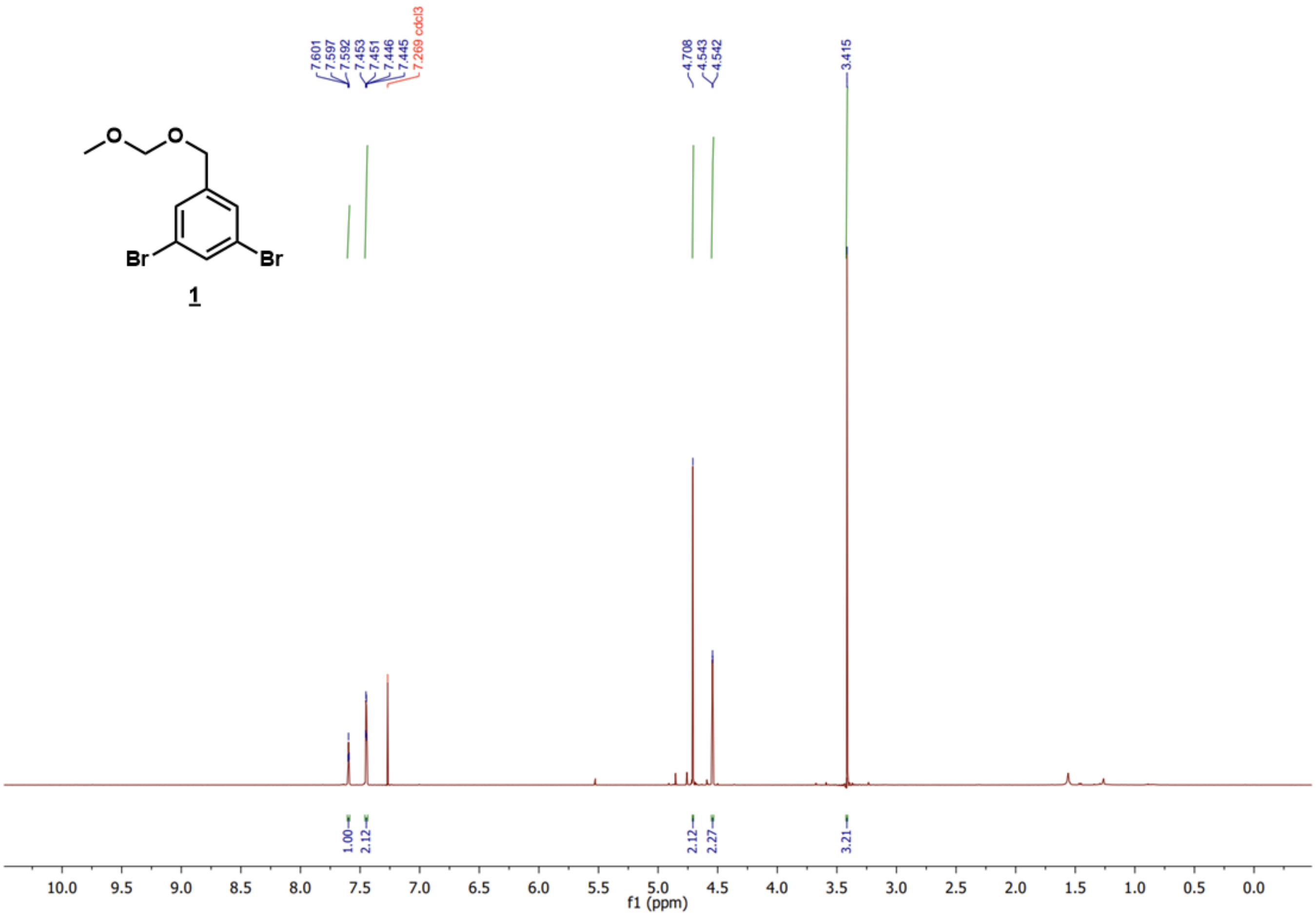

$^{13}C$ NMR of compound 1

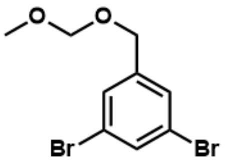

1

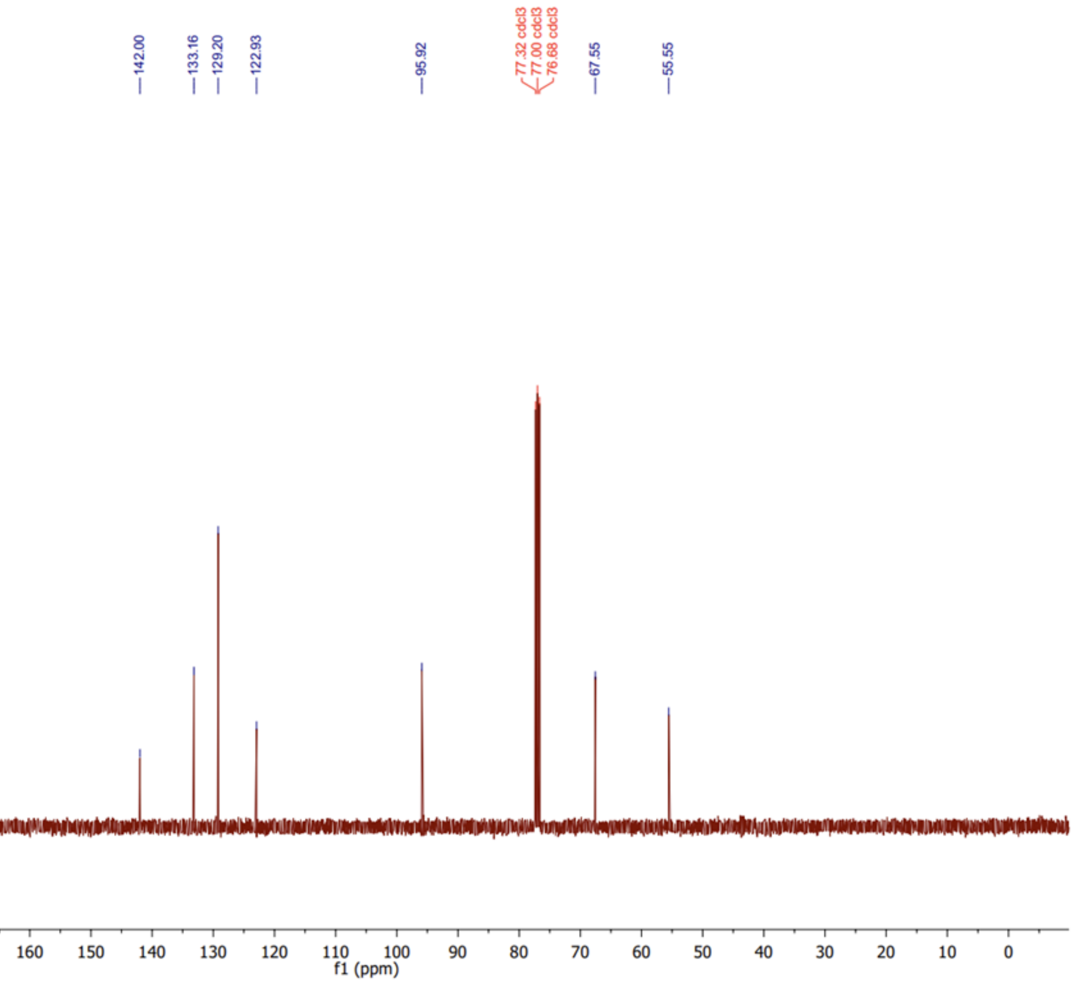

#### [1]H NMR of compound 2

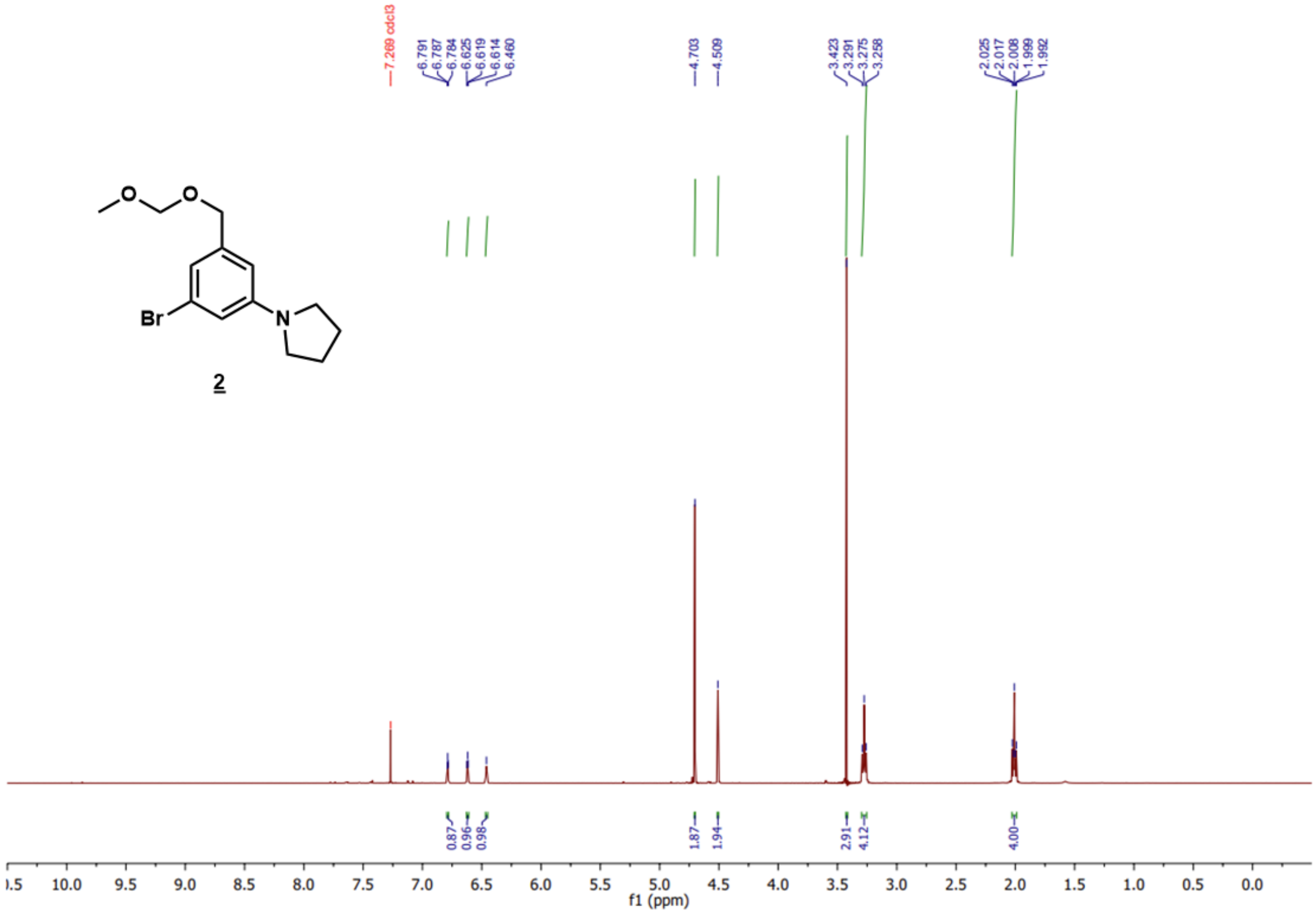

2

$^{13}$C NMR of compound 2

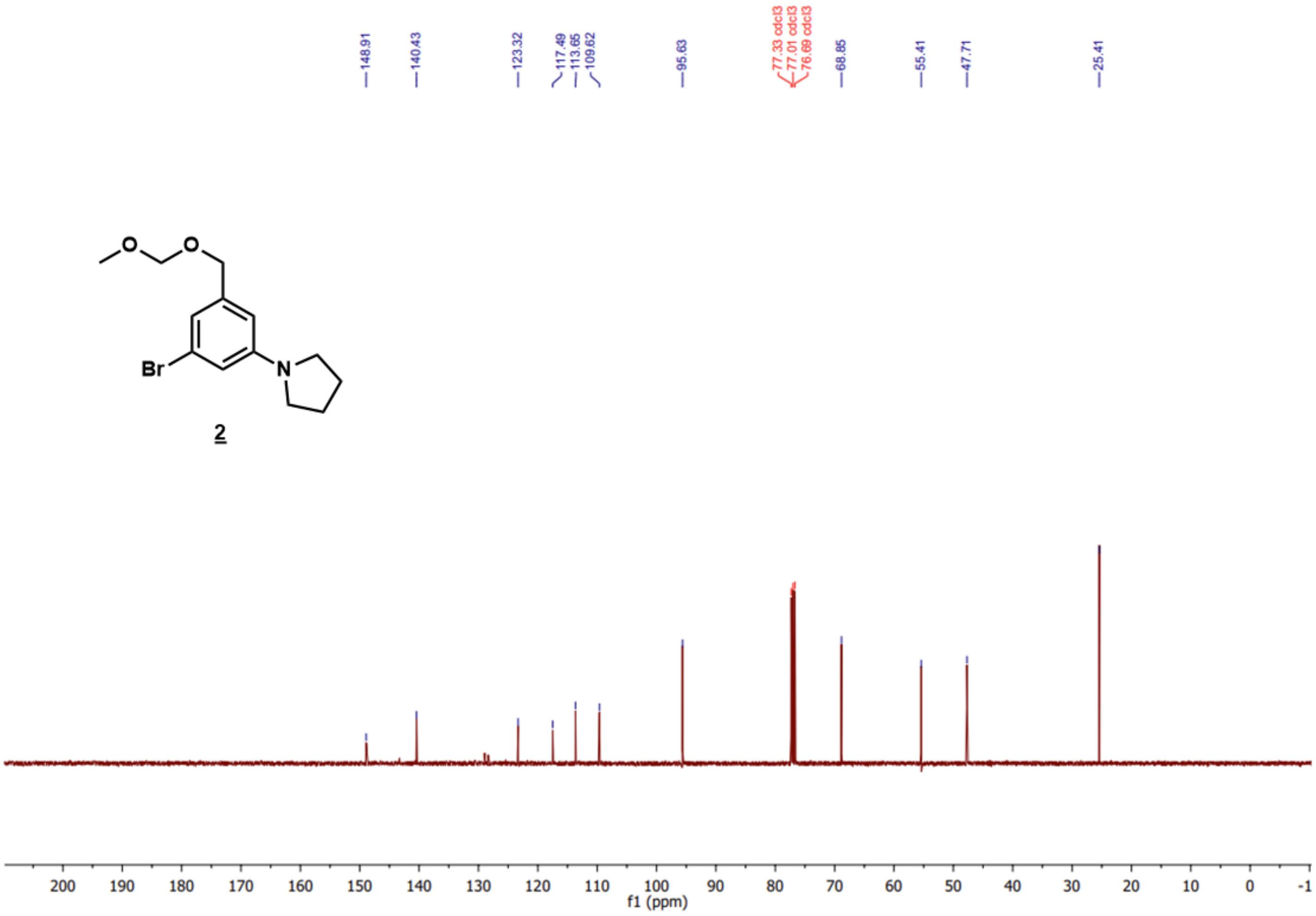

[1]H NMR of compound 3

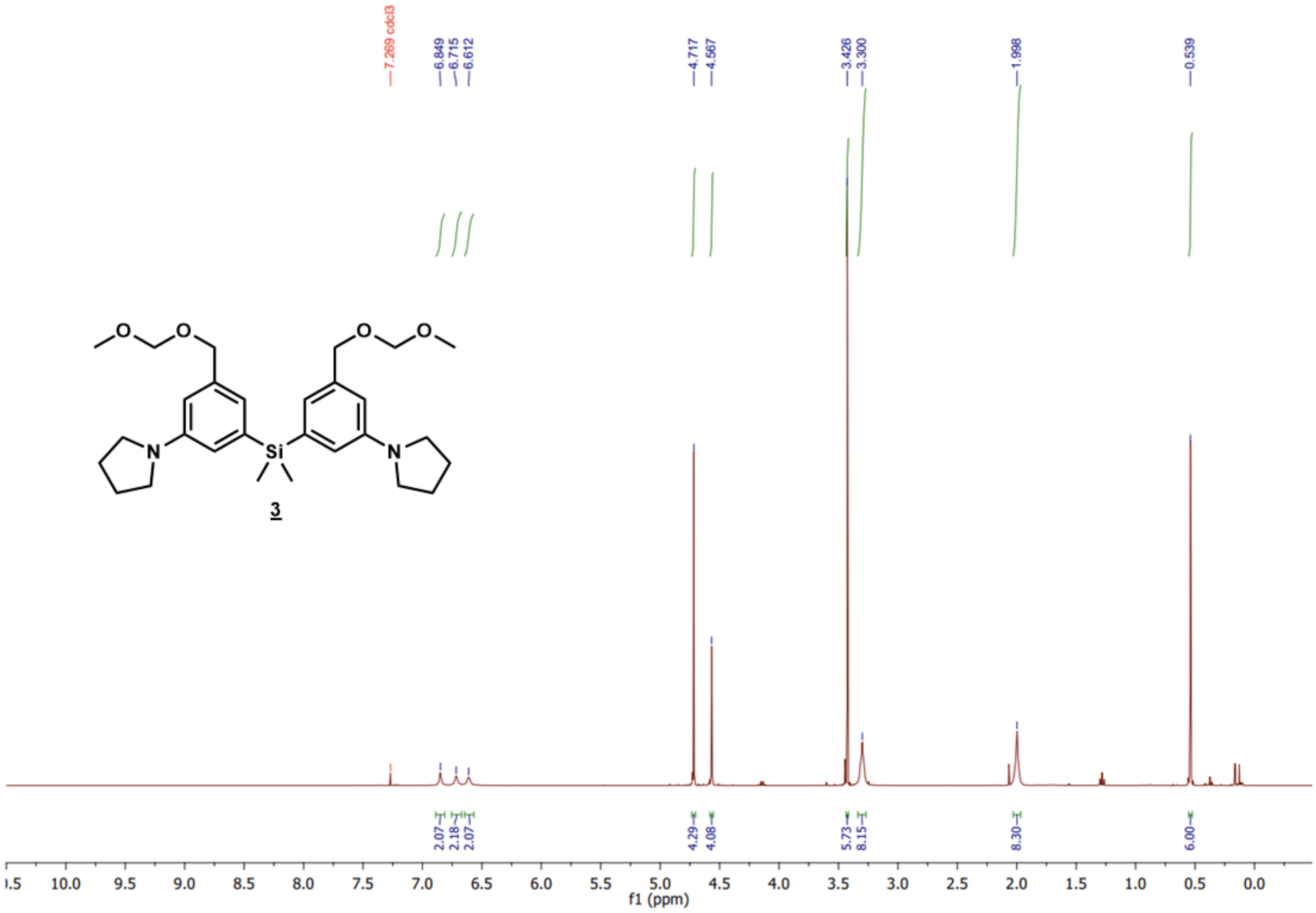

$^{13}C$ NMR of compound 3

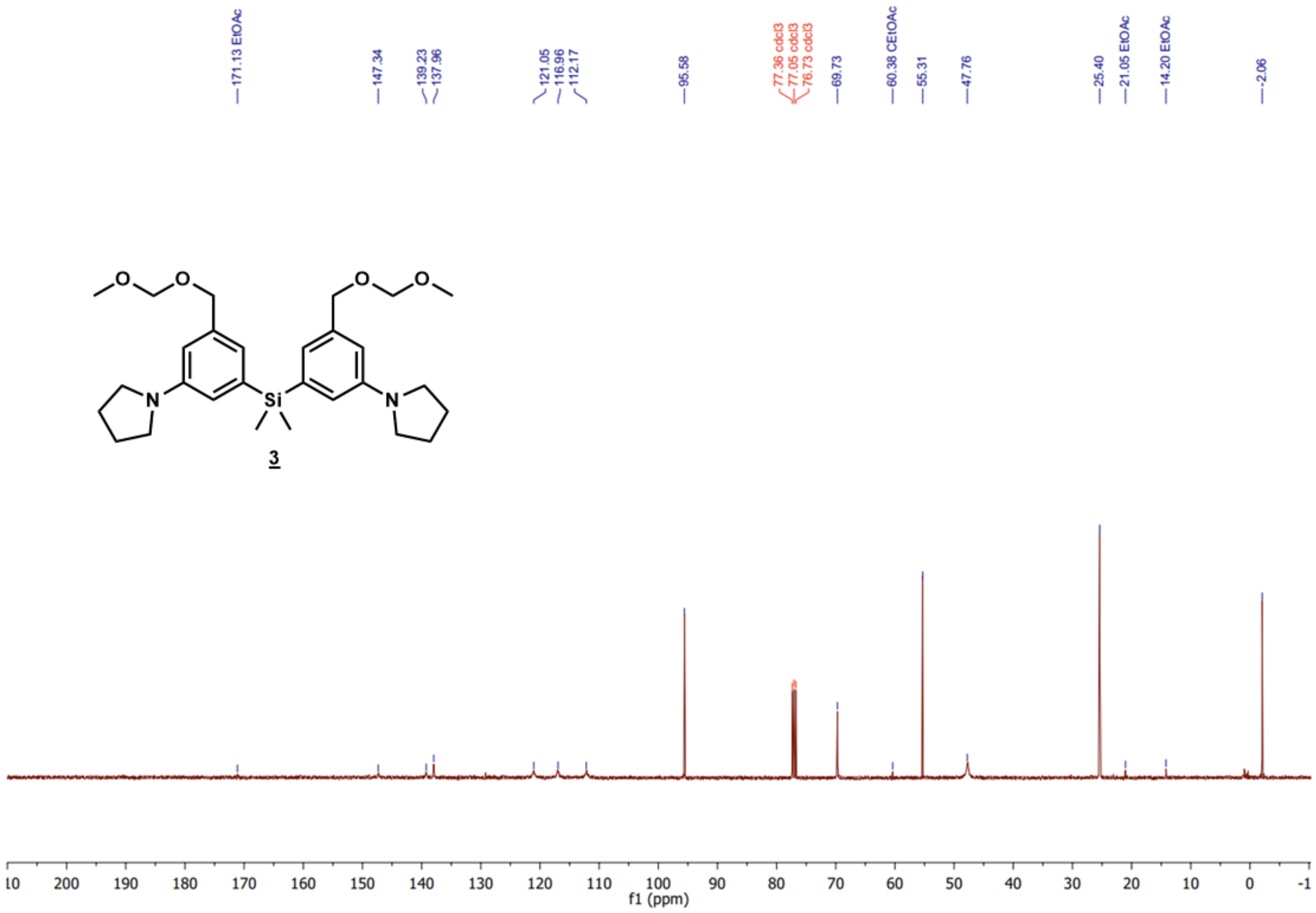

[1]H NMR of compound 4

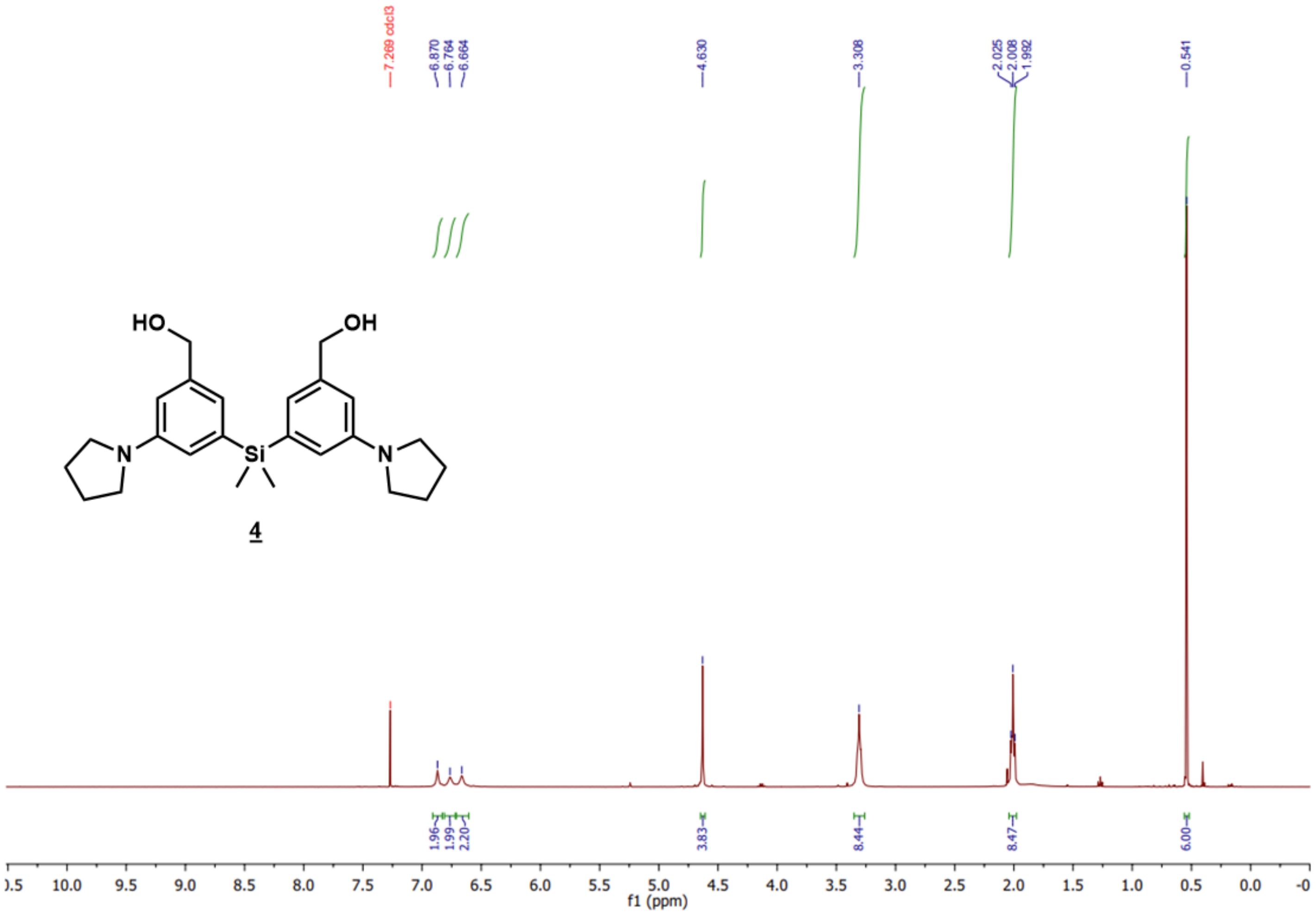

**$^{13}$C NMR of compound 4**

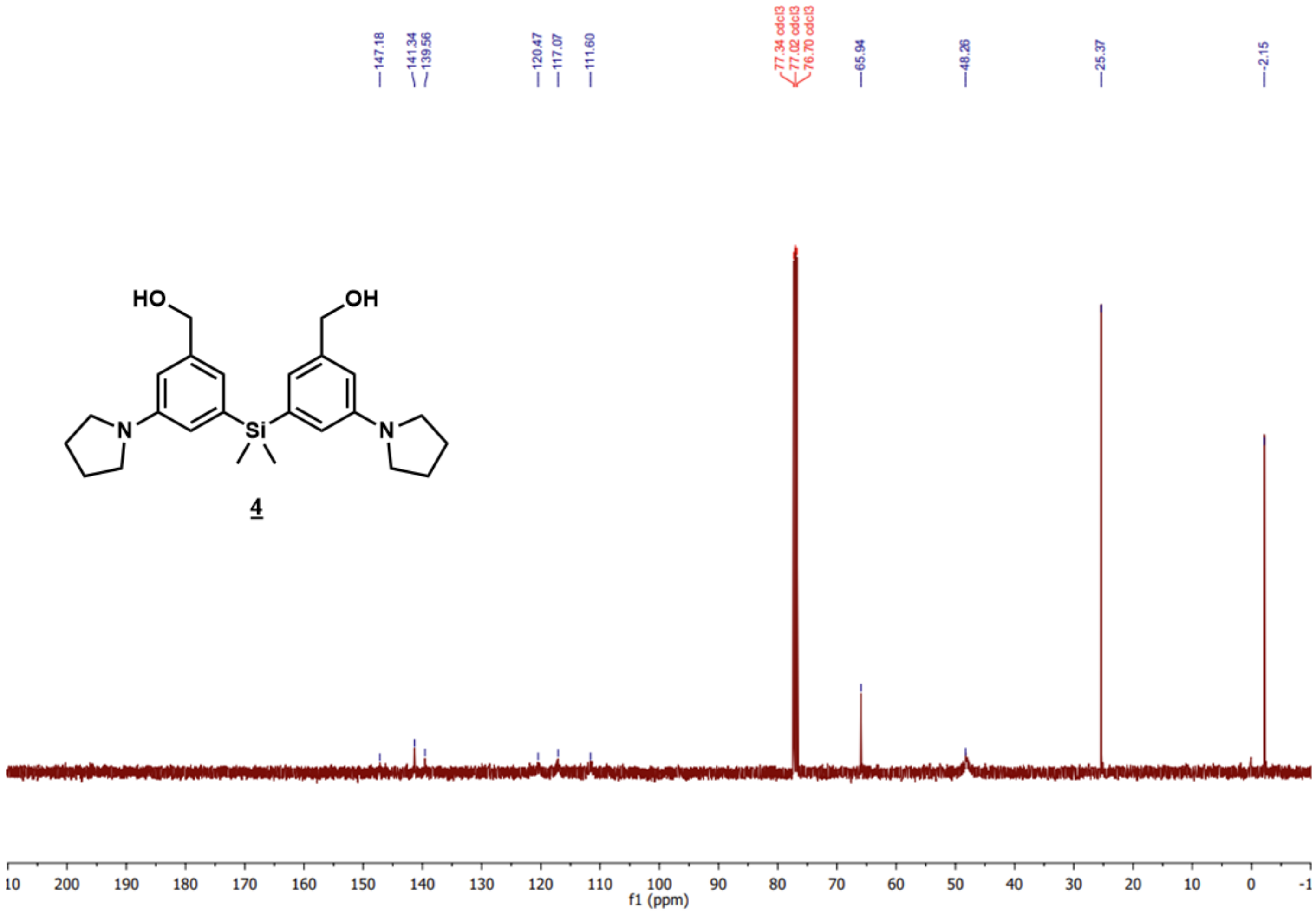

$^{1}$H NMR of compound 5

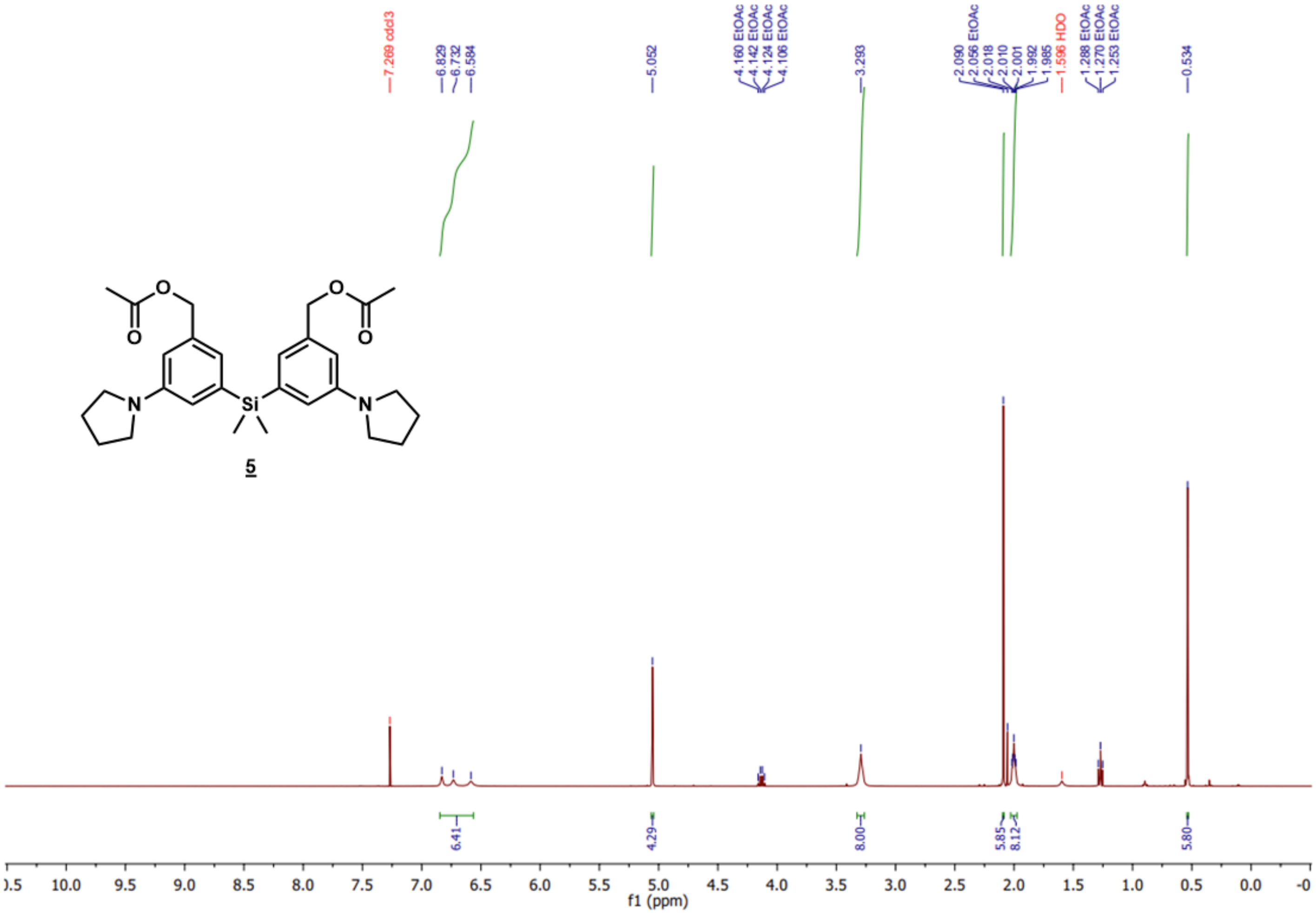

**13C NMR of compound 5**

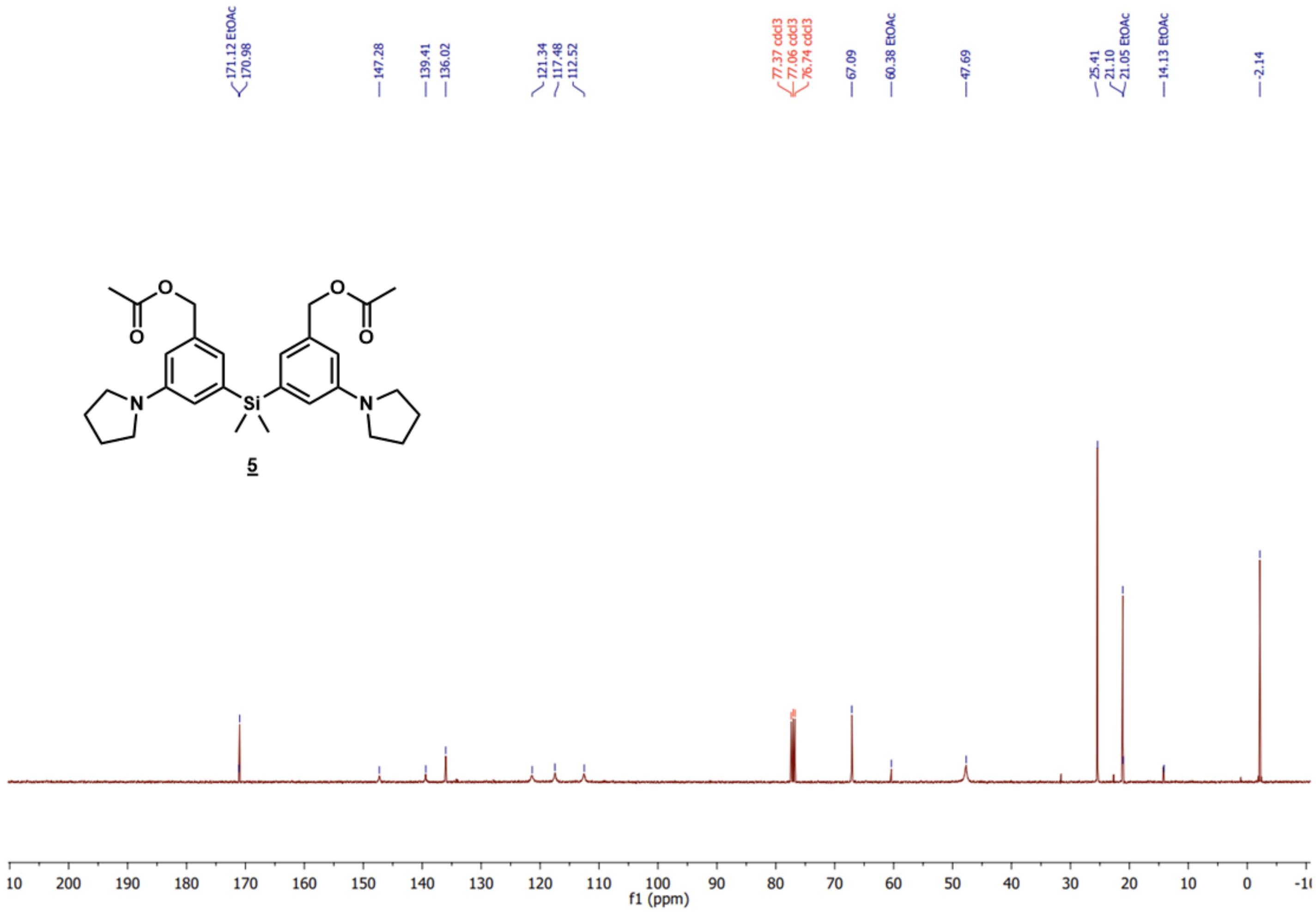

**$^1$H NMR of compound 6**

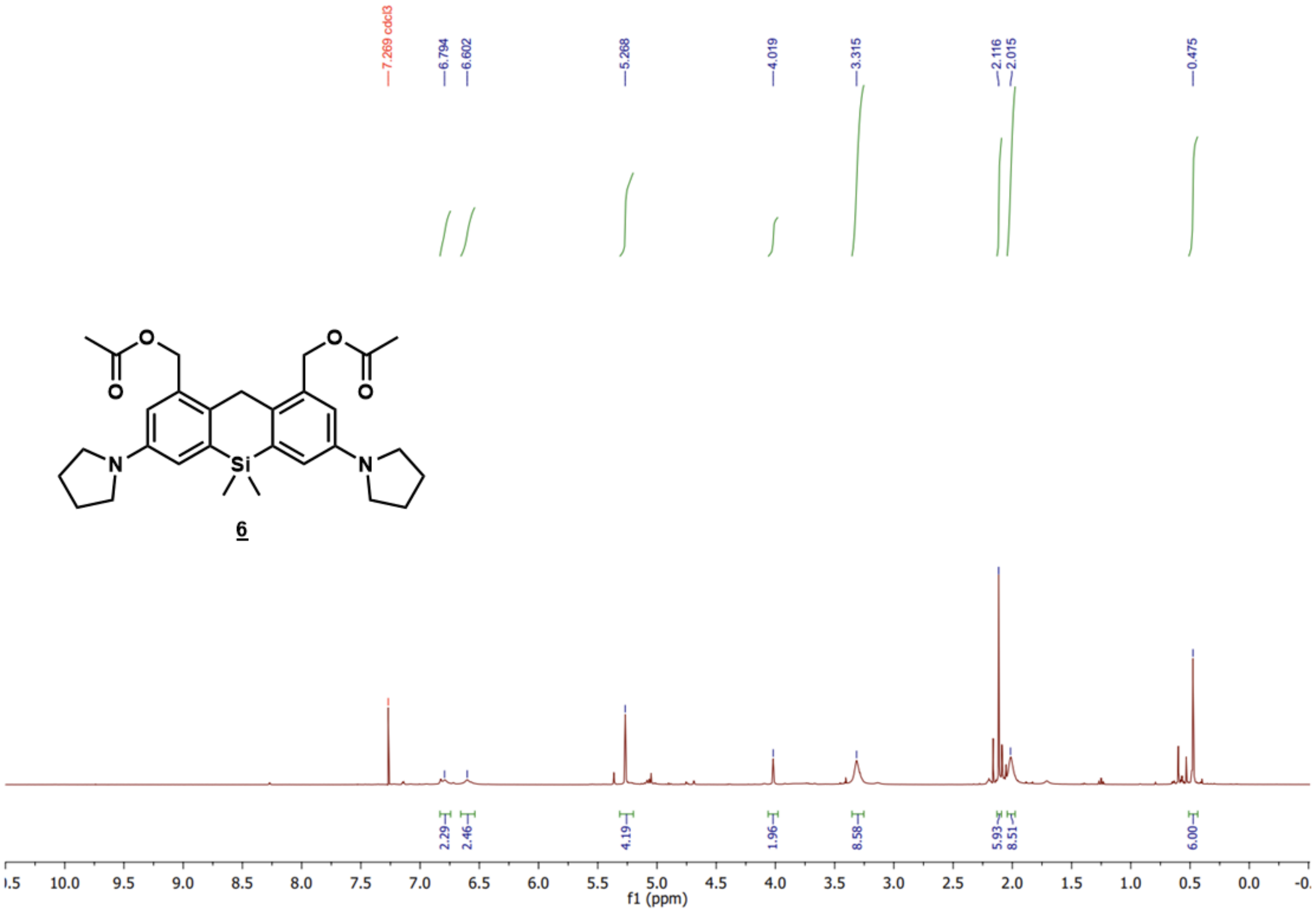

**$^1$H NMR of compound 7**

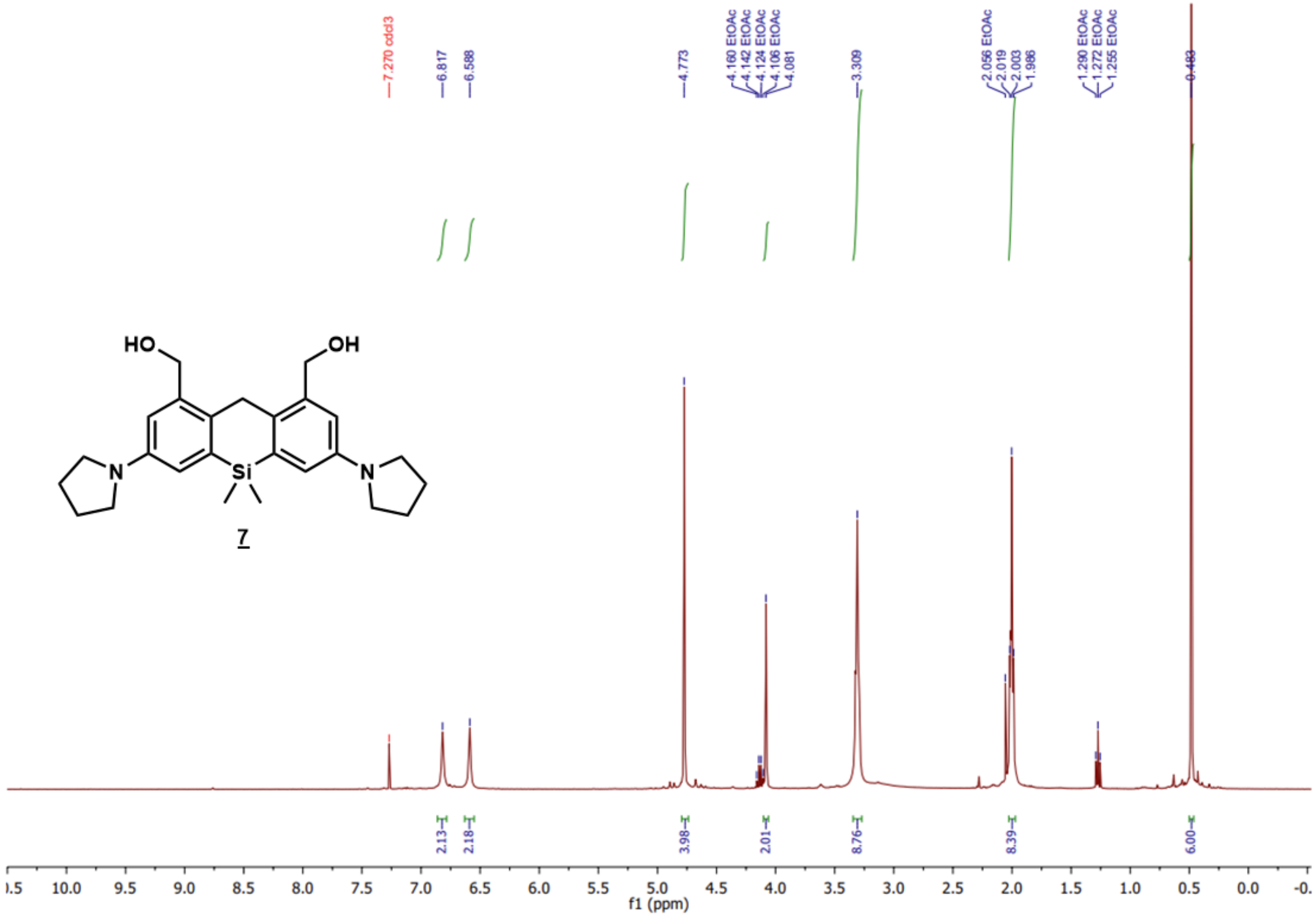

**$^{13}$C NMR of compound 7**

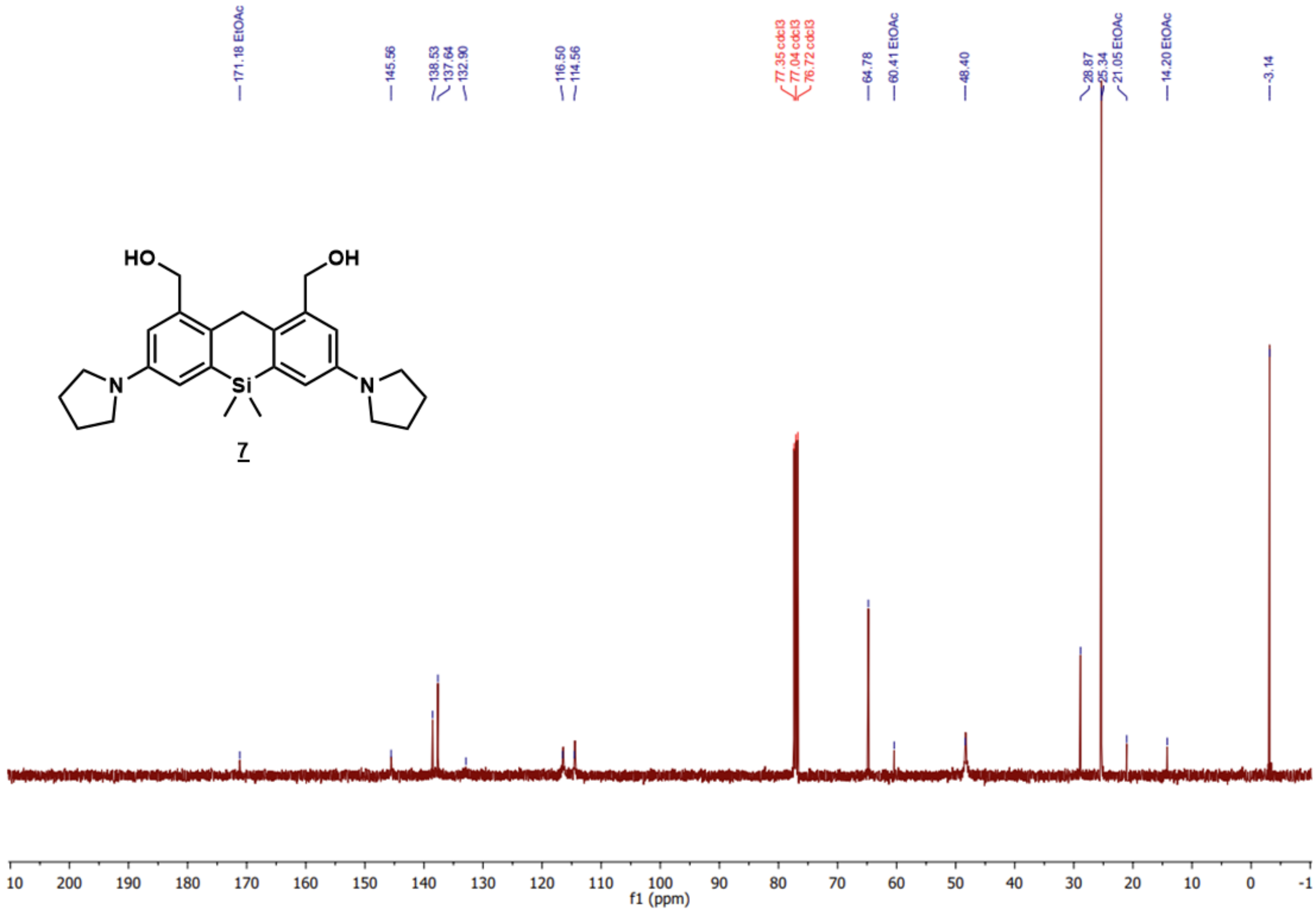

**[1]H NMR of compound 8**

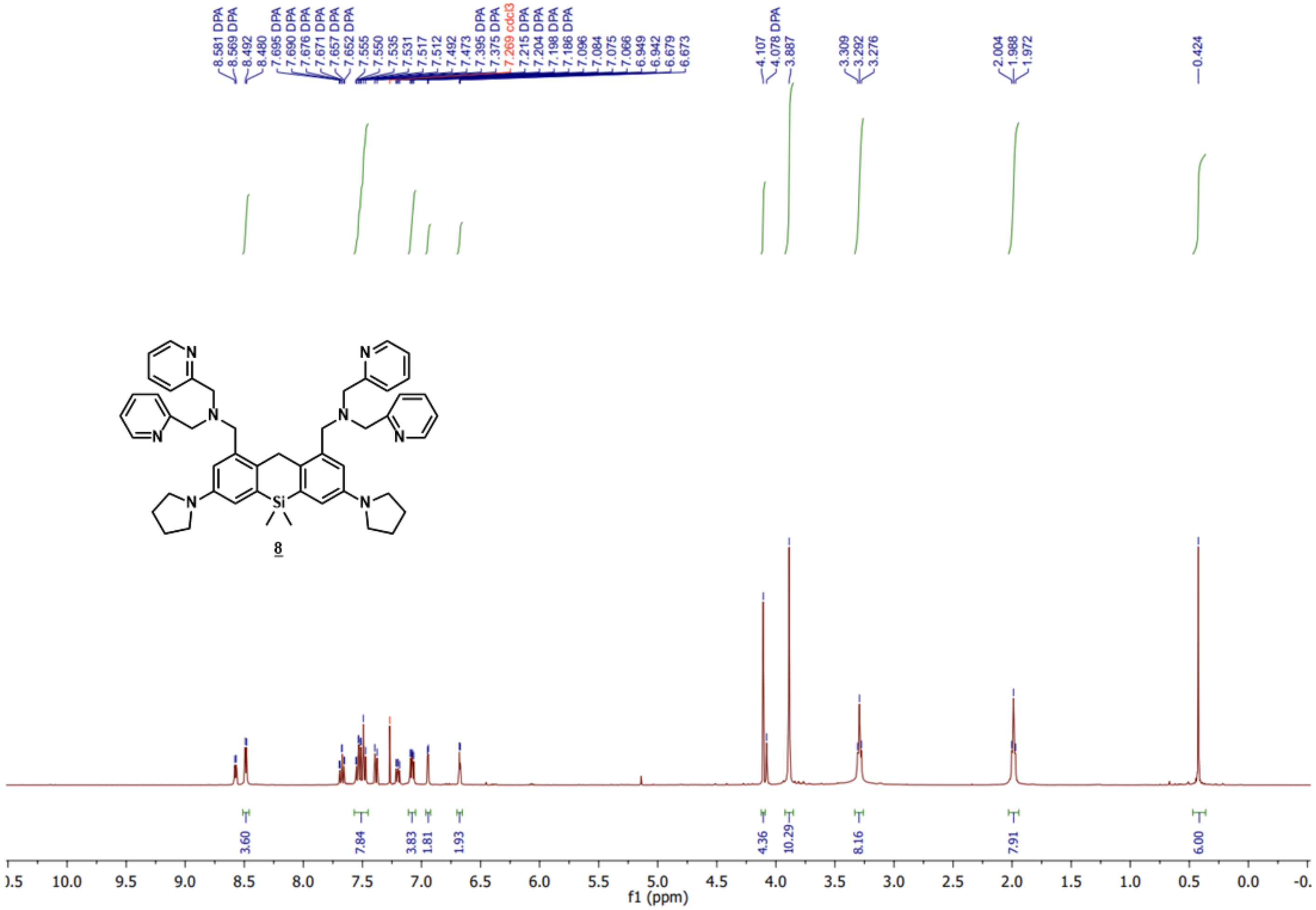

Trace of DPA was also observed.

**$^{13}$C NMR of compound 8**

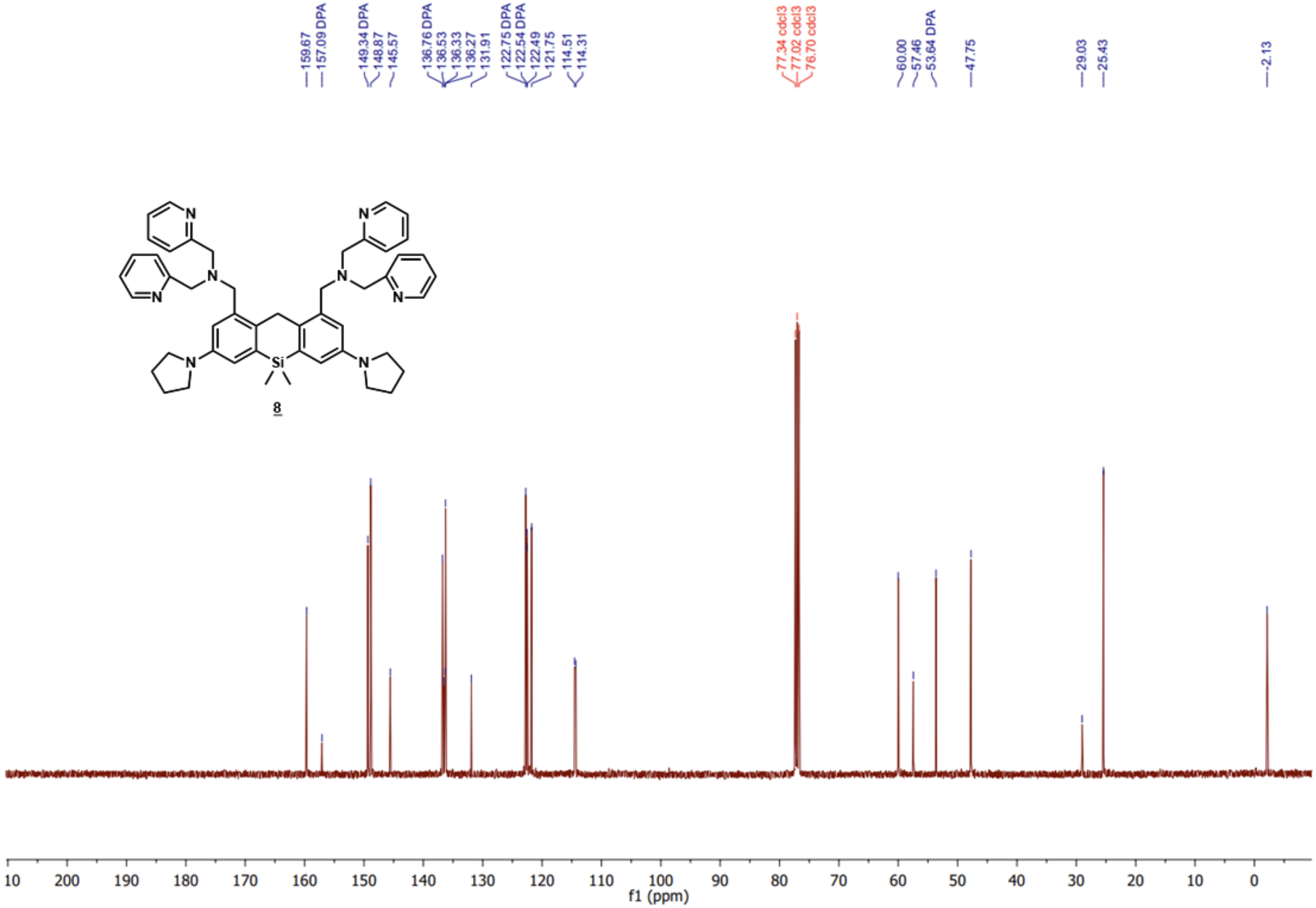

Trace of DPA was also observed.

[1]H NMR of DPA

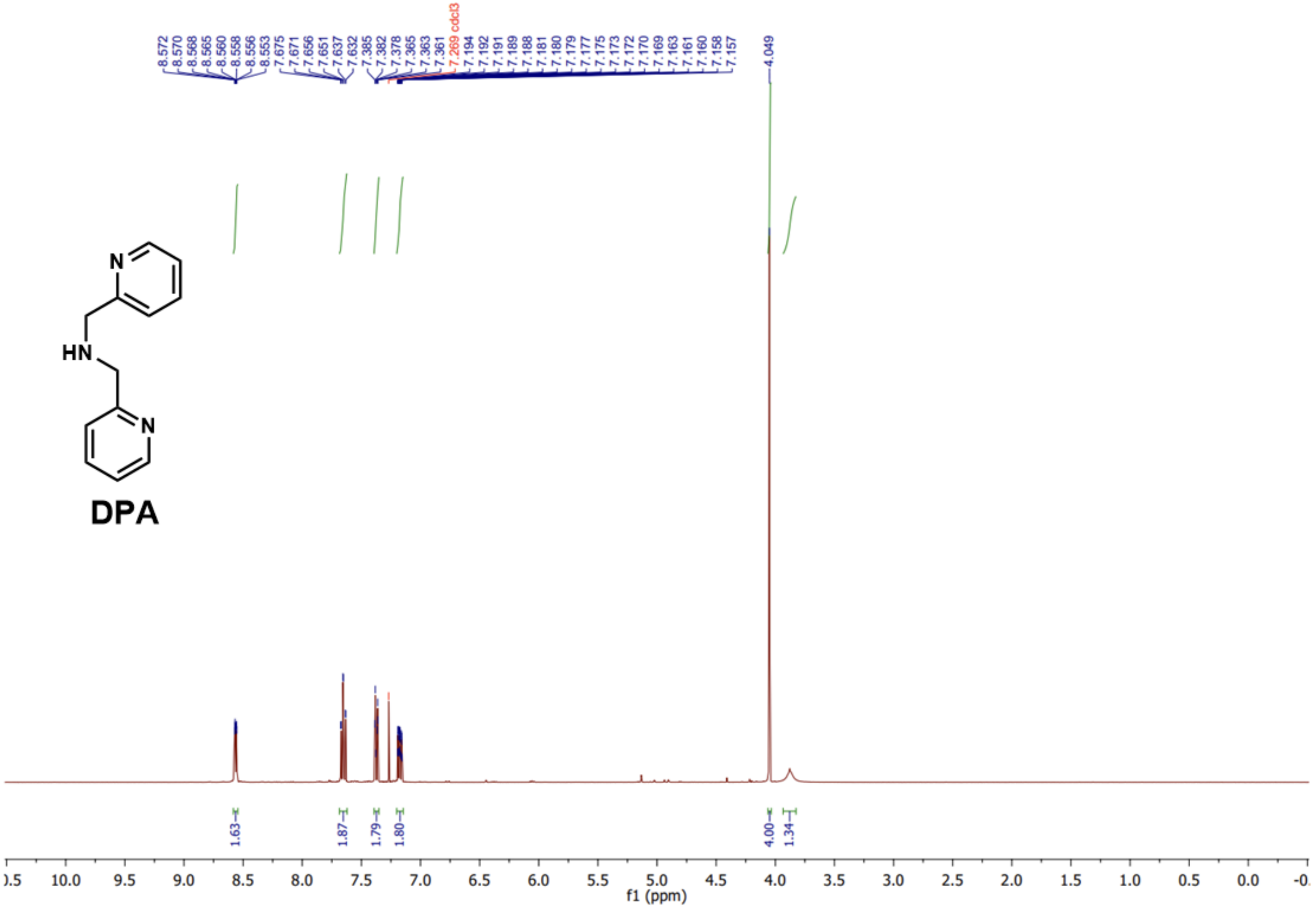

8.572
8.570
8.568
8.565
8.560
8.558
8.556
8.553
7.675
7.671
7.656
7.651
7.637
7.632
7.385
7.382
7.378
7.365
7.363
7.361
7.269 cdcl3
7.194
7.192
7.191
7.189
7.188
7.181
7.180
7.179
7.177
7.175
7.173
7.172
7.170
7.169
7.163
7.161
7.160
7.158
7.157
4.049
N
HN
N
DPA
1.63
1.87
1.79
1.80
4.00
1.34
f1 (ppm)

**$^{13}C$ NMR of DPA**

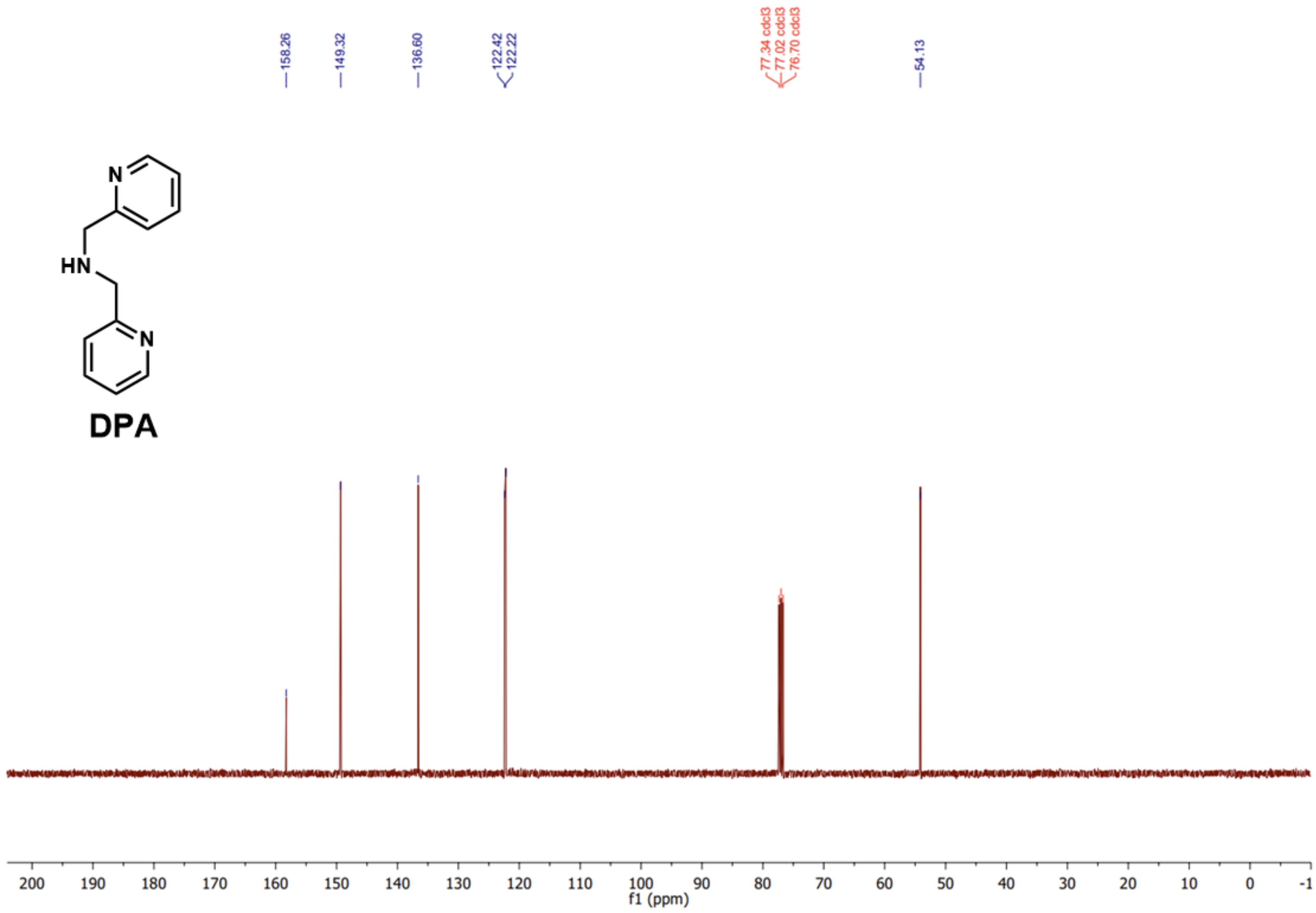

$^{1}$H NMR of SiX-DPA

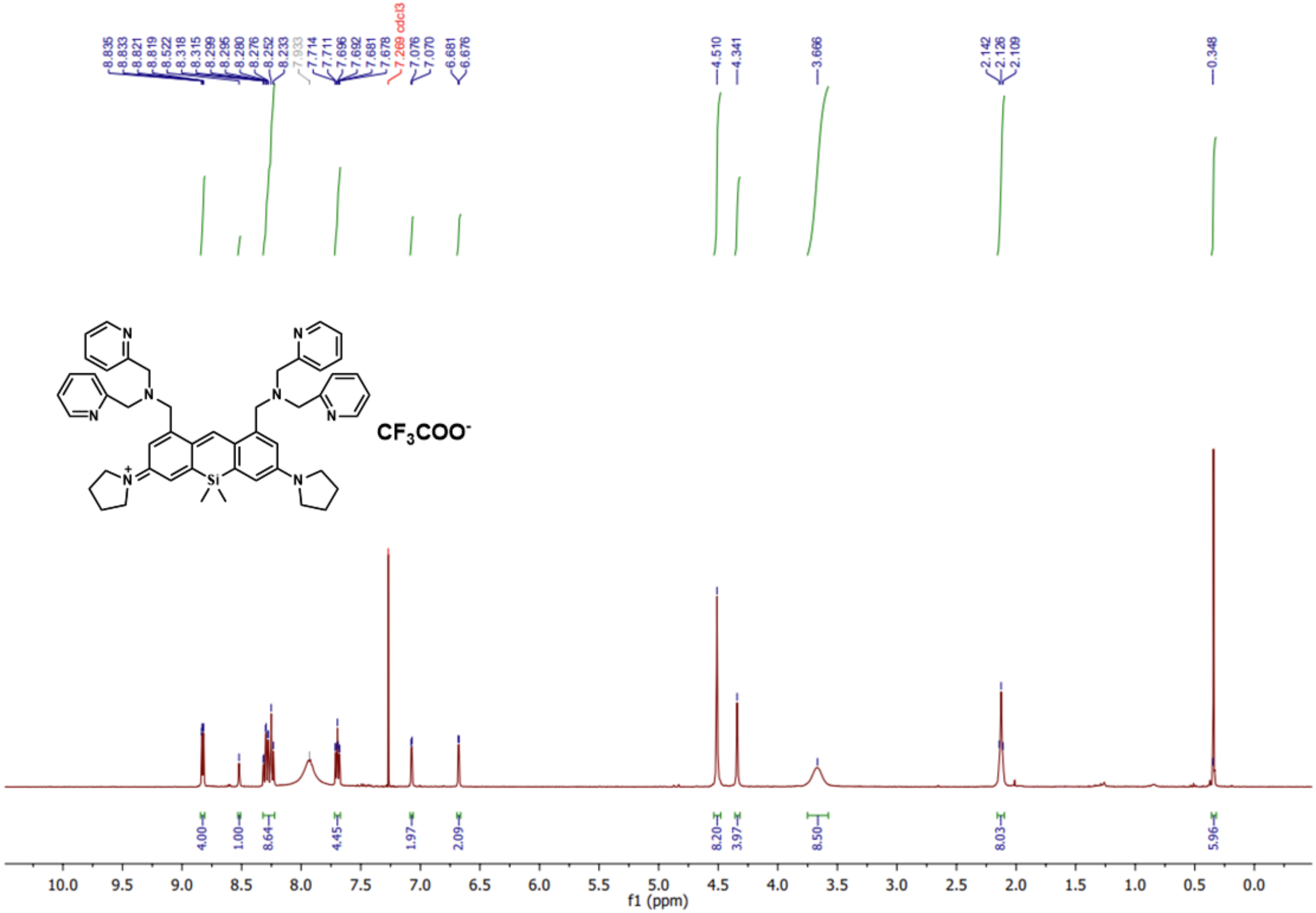

A broad signal centered at ~7.93 ppm was observed, which is likely due to hydrogen bonding (hydration) between the pyridine moiety of SiX-DPA and residual water molecules.

$^{13}C$ NMR of SiX-DPA

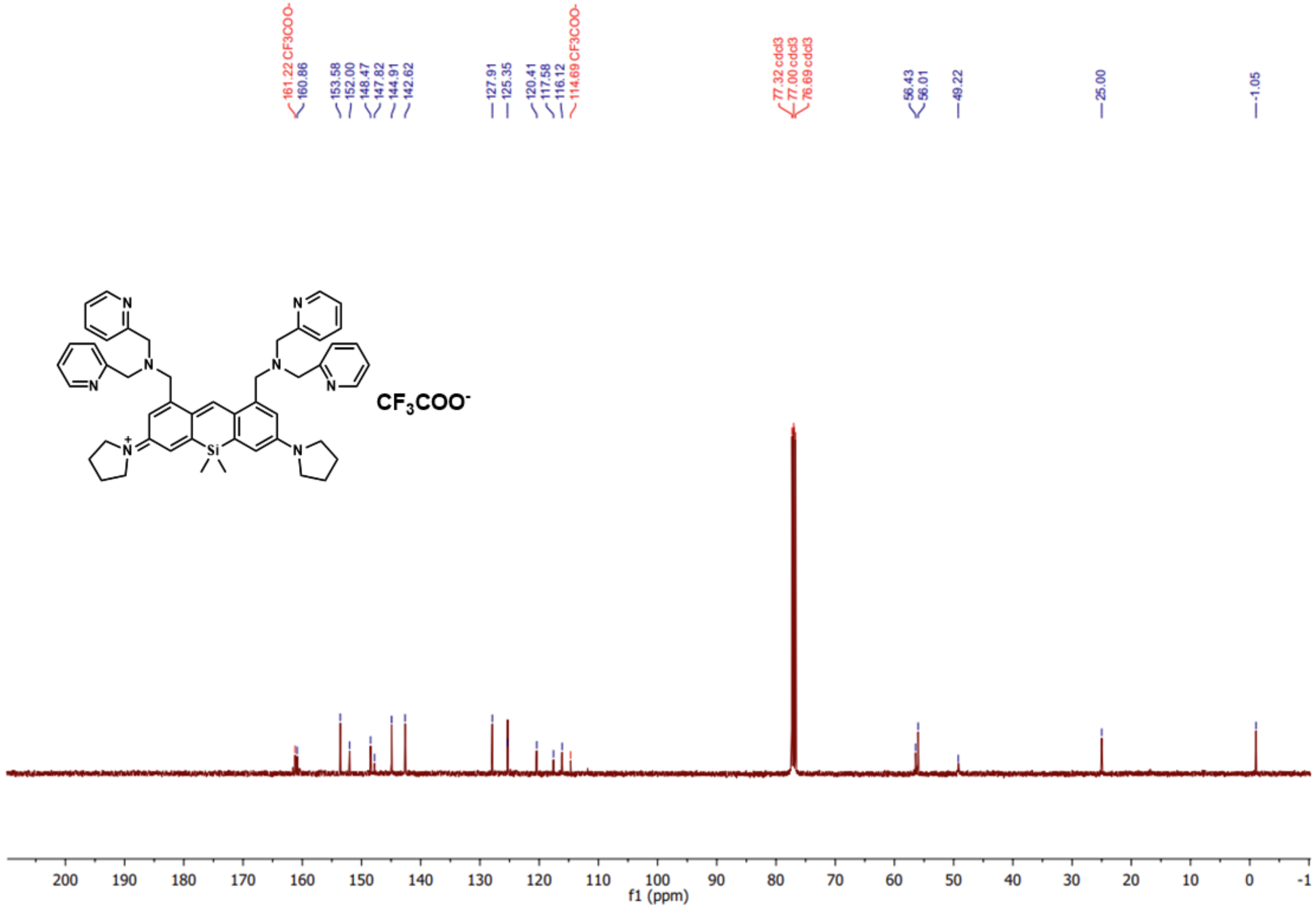

#### References

(1) Ojida, A.; Takashima, I.; Kohira, T.; Nonaka, H.; Hamachi, I. Turn-on Fluorescence Sensing of Nucleoside Polyphosphates Using a Xanthene-Based Zn(ii) Complex Chemosensor. *J. Am. Chem. Soc.* **2008**, *130* (36), 12095–12101. https://doi.org/10.1021/ja803262w.

(2) Martin, P.; van Mooy, B. A. S. Fluorometric Quantification of Polyphosphate in Environmental Plankton Samples: Extraction Protocols, Matrix Effects, and Nucleic Acid Interference. Appl. Environ. Microbiol. **2013**, *79* (1), 273-281. https://journals.asm.org/doi/10.1128/aem.02592-12.

(3) Christ, J. J.; Willbold, S.; Blank, L. M. Methods for the Analysis of Polyphosphate in the Life Sciences. *Anal. Chem.* **2020**, *92* (6), 4167–4176. https://doi.org/10.1021/acs.analchem.9b05144.

(4) Borghi, F.; Azevedo, C.; Johnson, E.; Burden, J. J.; Saiardi, A. A Mammalian Model Reveals Inorganic Polyphosphate Channeling into the Nucleolus and Induction of a Hyper-Condensate State. *Cell Rep. Methods* **2024**, *4* (7), 100814. https://doi.org/10.1016/j.crmeth.2024.100814.

(5) Diaz, J. M.; Ingall, E. D. Fluorometric Quantification of Natural Inorganic Polyphosphate. *Environ. Sci. Technol.* **2010**, *44* (12), 4665–4671. https://doi.org/10.1021/es100191h.

(6) Yang, X.; Gao, R.; Zhang, Q.; Yung, C. C. M.; Yin, H.; Li, J. Quantification of Polyphosphate in Environmental Planktonic Samples Using a Novel Fluorescence Dye JC-D7. *Environ. Sci. Technol.* **2024**. https://doi.org/10.1021/acs.est.4c04545.

(7) Smith, S. A.; Morrissey, J. H. Sensitive Fluorescence Detection of Polyphosphate in Polyacrylamide Gels Using 4′,6-Diamidino-2-Phenylindol. *ELECTROPHORESIS* **2007**, *28* (19), 3461–3465. https://doi.org/10.1002/elps.200700041.

(8) Fulmer, G. R.; Miller, A. J. M.; Sherden, N. H.; Gottlieb, H. E.; Nudelman, A.; Stoltz, B. M.; Bercaw, J. E.; Goldberg, K. I. NMR Chemical Shifts of Trace Impurities: Common Laboratory Solvents, Organics, and Gases in Deuterated Solvents Relevant to the Organometallic Chemist. *Organometallics* **2010**, *29* (9), 2176–2179. https://doi.org/10.1021/om100106e.

(9) Manoukian, L.; Stein, R. S.; Correa, J. A.; Frigon, D.; Omelon, S. Short-Chain Polyphosphates: Extraction Effects on Migration and Size Estimation of Saccharomyces Cerevisiae Extracts with Polyacrylamide Gel Electrophoresis. *ELECTROPHORESIS* **2023**, *44* (15–16), 1197–1205. https://doi.org/10.1002/elps.202300055.

(10) Smith, S. A.; Wang, Y.; Morrissey, J. H. DNA Ladders Can Be Used to Size Polyphosphate Resolved by Polyacrylamide Gel Electrophoresis. *Electrophoresis* **2018**, *39* (19), 2454–2459. https://doi.org/10.1002/elps.201800227.

(11) Manoukian, L.; Correa, J. A.; Stein, R. S.; Frigon, D.; Omelon, S. Extraction Processes Reduce Polyphosphate Ion Migration, Dispersion and Diffusion as Detected with Gel Electrophoresis and 31P DOSY NMR. *ELECTROPHORESIS* **2022**, *43* (20), 2014–2022. https://doi.org/10.1002/elps.202100364.

(12) Hofmann, H.; Soranno, A.; Borgia, A.; Gast, K.; Nettels, D.; Schuler, B. Polymer Scaling Laws of Unfolded and Intrinsically Disordered Proteins Quantified with Single-Molecule Spectroscopy. *Proc. Natl. Acad. Sci.* **2012**, *109* (40), 16155–16160. https://doi.org/10.1073/pnas.1207719109.

(13) Sun, L.-Z.; Qian, J.-L.; Cai, P.; Hu, H.-X.; Xu, X.; Luo, M.-B. Mg2+ Effects on the Single-Stranded DNA Conformations and Nanopore Translocation Dynamics. *Polymer* **2022**, *250*, 124895. https://doi.org/10.1016/j.polymer.2022.124895.

(14) Tan, K. Y.; Herr, A. E. Ferguson Analysis of Protein Electromigration during Single-Cell Electrophoresis in an Open Microfluidic Device. *The Analyst* **2020**, *145* (10), 3732–3741. https://doi.org/10.1039/c9an02553g.

(15) Chung, M.; Kim, D.; Herr, A. E. Polymer Sieving Matrices in Microanalytical Electrophoresis. *Analyst* **2014**, *139* (22), 5635–5654. https://doi.org/10.1039/C4AN01179A.

(16) Neville, D. M. Molecular Weight Determination of Protein-Dodecyl Sulfate Complexes by Gel Electrophoresis in a Discontinuous Buffer System. *J. Biol. Chem.* **1971**, *246* (20), 6328–6334. https://doi.org/10.1016/S0021-9258(18)61792-2.

(17) Andersson, K.; Fagerlind, M.; Daneholt, B. The Relationship between Molecular Size and Electrophoretic Mobility in Agarose Gels as Determined from a Single Population of Growing RNA Molecules. *Mol. Biol. Rep.* **1975**, *2* (3), 195–201. https://doi.org/10.1007/BF00356988.

(18) Pilatus, U.; Mayer, A.; Hildebrandt, A. Nuclear Polyphosphate as a Possible Source of Energy during the Sporulation of *Physarum Polycephalum*. *Arch. Biochem. Biophys.* **1989**, *275* (1), 215–223. https://doi.org/10.1016/0003-9861(89)90366-4.

(19) Schindelin, J.; Arganda-Carreras, I.; Frise, E.; Kaynig, V.; Longair, M.; Pietzsch, T.; Preibisch, S.; Rueden, C.; Saalfeld, S.; Schmid, B.; Tinevez, J.-Y.; White, D. J.; Hartenstein, V.; Eliceiri, K.; Tomancak, P.; Cardona, A. Fiji: An Open-Source Platform for Biological-Image Analysis. *Nat. Methods* **2012**, *9* (7), 676–682. https://doi.org/10.1038/nmeth.2019.
